# Development of SYK NanoBRET Cellular Target Engagement Assays for Gain–of–Function Variants

**DOI:** 10.1101/2024.06.12.598544

**Authors:** Jacob L. Capener, James D. Vasta, Vittorio L. Katis, Ani Michaud, Michael T. Beck, Sabrina C. D. Daglish, Sarit Cohen-Kedar, Efrat Shaham Barda, Stefanie Howell, Iris Dotan, Matthew B. Robers, Alison D. Axtman, Frances M. Bashore

## Abstract

Spleen tyrosine kinase (SYK) is a non-receptor tyrosine kinase that is activated by phosphorylation events downstream of FcR, B-cell and T-cell receptors, integrins, and C-type lectin receptors. When the tandem Src homology 2 (SH2) domains of SYK bind to phosphorylated immunoreceptor tyrosine-based activation motifs (pITAMs) contained within these immunoreceptors, or when SYK is phosphorylated in interdomain regions A and B, SYK is activated. SYK gain-of-function (GoF) variants were previously identified in six patients that had higher levels of phosphorylated SYK and phosphorylated downstream proteins JNK and ERK. Furthermore, the increased SYK activation resulted in the clinical manifestation of immune dysregulation, organ inflammation, and a predisposition for lymphoma. The knowledge that the SYK GoF variants have enhanced activity was leveraged to develop a SYK NanoBRET cellular target engagement assay in intact live cells with constructs for the SYK GoF variants. Herein, we developed a potent SYK-targeted NanoBRET tracer using a SYK donated chemical probe, MRL-SYKi, that enabled a NanoBRET cellular target engagement assay for SYK GoF variants, SYK(S550Y), SYK(S550F), and SYK(P342T). We determined that ATP-competitive SYK inhibitors bind potently to these SYK variants in intact live cells. Additionally, we demonstrated that MRL-SYKi can effectively reduce the catalytic activity of SYK variants, and the phosphorylation levels of SYK(S550Y) in an epithelial cell line (SW480) stably expressing SYK(S550Y).

## 1 Introduction

Spleen tyrosine kinase (SYK) is a non-receptor tyrosine kinase that contains a C-terminal kinase domain and two N-terminal tandem SH2 domains linked via interdomain loop regions A and B (Figure 1A) (Sada et al., 2001;Zhou et al., 2023). SYK remains in an autoinhibited conformation, and based on structural data, this is due to interactions between the interdomain regions and the C-terminal helix of the kinase (Kulathu et al., 2009;Grädler et al., 2013;Mansueto et al., 2019). SYK is activated by the interaction between its tandem SH2 domains and phosphorylated immunoreceptor tyrosine-based activation motifs (pITAMs), or by tyrosine phosphorylation (Figure 1A&C) (Mócsai et al., 2010;Antenucci et al., 2018;Mansueto et al., 2019). ITAMs are contained within immunoreceptors such as FcR, B-cell, T-cell, and C-type lectin receptors (Kiefer et al., 1998;Mócsai et al., 2010;Ben Mkaddem et al., 2019). Recently, SYK monoallelic gain-of-function (GoF) variants were identified from six patients with clinical presentation of immune deficiency, arthritis, dermatitis, colitis, and diffuse large B-cell lymphomas (Wang et al., 2021). The variants p.Pro342Thr (P342T) and p.Ala353Thr (A352T) are located in the interdomain B region, and p.Ser550Tyr (S550Y), p.Ser550Phe (S550F), and p.Met450Ile (M450I) are within the catalytic kinase domain (Figure 1B). The variants S550Y and S550F presented clinically in infants at 2 weeks, whereas P342T, A352T, and M450I presented in patients that were 12, 34, and 44 years old, respectively. These variants resulted in an increase in levels of Y525/526 phosphorylated SYK and downstream phosphorylation of c-Jun N-terminal kinase (JNK) and extracellular signal-regulated kinase (ERK) in HEK293 cells transfected with SYK variants. The S550 variants exhibited higher Y525/526 phosphorylation levels and enhanced downstream activation compared to the three other variants. Wang et al demonstrated that a promiscuous SYK inhibitor, R406, could decrease S550Y phosphorylation levels in a colonic epithelial cell line SW480 that stably expressed S550Y (Wang et al., 2021).

**Figure 1.**
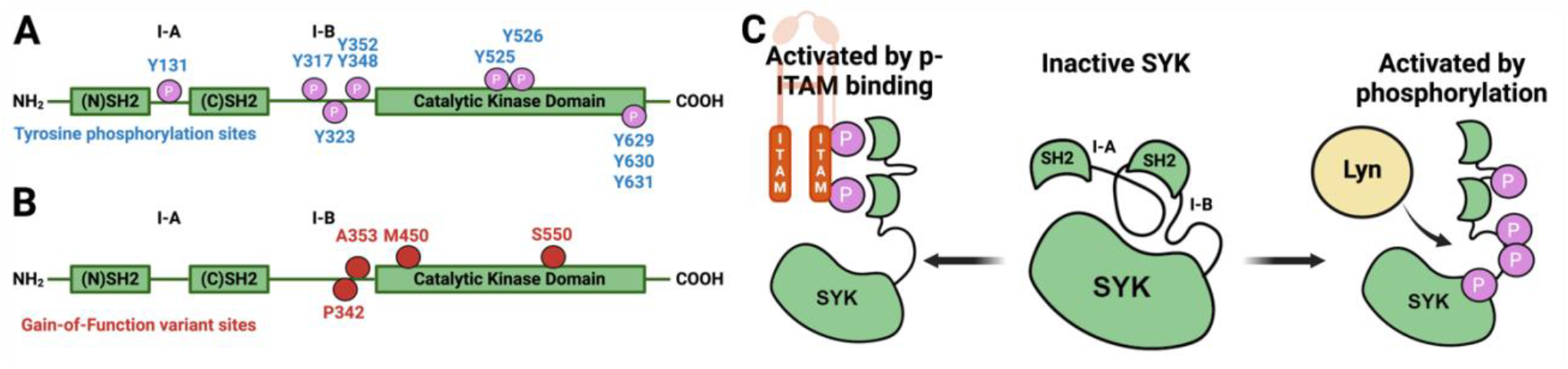
Diagrams that illustrate key sites on SYK. (**A**) SYK tyrosine phosphorylation sites. (**B**) SYK gain-of-function variant sites. (**C**) SYK activation mechanism through pITAM binding or via phosphorylation (autophosphorylation or through LYN kinase).

Small molecule ATP-competitive inhibitors of SYK include R406, entospletinib, cerdulatinib, P505-15, and MRL-SYKi (Braselmann et al., 2006;Altman et al., 2011;Coffey et al., 2012;Currie et al., 2014;Guo et al., 2017). These compounds potently inhibit SYK in enzymatic inhibition assays (Figure 2). R406 is a highly promiscuous kinase inhibitor that binds 79 kinases with a K^d^ <100 nM in the broad DiscoveRx KINOME*scan* panel of kinase binding assays (Braselmann et al., 2006;Rolf et al., 2015;Sharman et al., 2015). R406 is also a metabolite of the pro-drug fostamatinib, which is used as a treatment for chronic immune thrombocytopenia (ITP) due to SYK’s role in immunoreceptor signaling (Connell and Berliner, 2019). Fostamatinib is also in phase 2 and 3 clinical trials for rheumatoid arthritis, autoimmune hemolytic anemia, cutaneous lupus erythematosus, and COVID-19 (Cooper et al., 2023). Entospletinib (GS-9973) only binds one off-target kinase, TNK1, with a K^d^ <100 nM (Sharman et al., 2015;Cooper et al., 2023). SYK’s role in BCR signaling has led to ongoing clinical trials with entospletinib for a variety of hematologic malignancies, such as acute myelogenous leukemia, acute lymphocytic leukemia, and diffuse large B-cell lymphoma, among others (Liu and Mamorska-Dyga, 2017). Cerdulatinib (PRT062070), is a dual SYK/JAK1 inhibitor (JAK1 IC^50^ = 12 nM) in clinical trials for B-cell malignancies (Ma et al., 2015;Guo et al., 2017). Cerdulatinib also inhibits at least 22 off-target kinases with an IC^50^ <200 nM, therefore it also lacks selectivity for SYK (Ma et al., 2015;Liu and Mamorska-Dyga, 2017). P505-15 (PRT062607) also inhibits eight kinases by greater than 80% at 300 nM in enzymatic assays (FGR, MLK1, YES, FLT3, PAK5, LYN, SRC, and LCK) (Coffey et al., 2012;Spurgeon et al., 2013). When a dose–response study of inhibition was carried out for these eight kinases, the closest off-target was found to be 81-fold less potent (enzymatic IC^50^ = 81 nM) when compared to SYK. Lastly, MRL-SYKi is a donated chemical probe with greater than 30-fold selectivity versus the protein kinase family included in the KINOME*scan* panel and no off-target inhibition at 10 μM in a GPCR scan and panel of 113 enzymes and receptors (SGC-Frankfurt). Off-target kinases inhibited in enzymatic inhibition assays with a potency between 100 and 242 nM (greater or equal to 111-fold less potent) are SRC, NTRK1, NTRK3, FGR, FYN, FER, and JAK2. Furthermore, this probe has cellular activity at <100 nM and is recommended for use in cellular assays at 100 nM (Altman et al., 2011).

**Figure 2.**
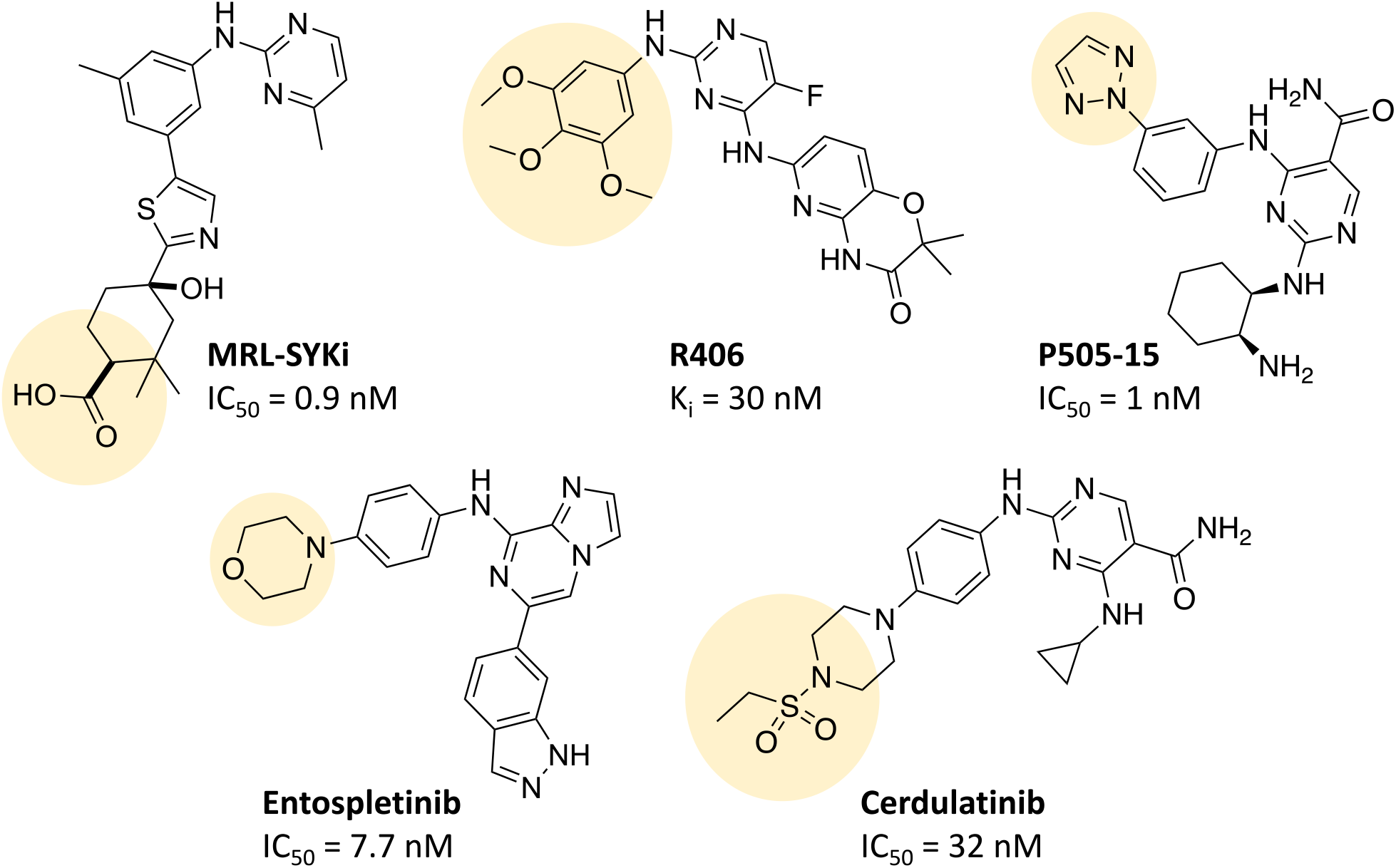
The chemical structures of SYK inhibitors and their solvent exposed sites (yellow). Corresponding enzymatic inhibition data from the literature is reported (SGC-Frankfurt;Braselmann et al., 2006;Altman et al., 2011;Coffey et al., 2012;Currie et al., 2014;Guo et al., 2017).

Due to a lack of cellular target engagement assays for SYK, assessing these inhibitors’ ability to bind directly to SYK in a cellular assay has not been possible. Currently, on-target cell activity for SYK is assessed in mast cell degranulation assays due to SYK’s pivotal role in FcεRI signaling (Siraganian et al., 2010). A peptide-based biosensor assay can also detect SYK kinase activation and inhibition within intact cells; however, this does not determine direct binding events (Lipchik et al., 2012). NanoBRET cellular target engagement assays are predominantly used for kinases to determine the ability of an inhibitor to displace a bioluminescence resonance energy transfer (BRET) probe from the NanoLuciferase (NL)-tagged protein in intact live cells, resulting in a cellular IC^50^ for binary target engagement (Machleidt et al., 2015;Vasta et al., 2018). BRET probes, often called NanoBRET tracers, contain a ligand for an NL-tagged protein and a BODIPY dye (NanoBRET 590 or BODIPY 576/589) attached with a linker. When the tracer binds to the protein, the BODIPY dye fluoresces upon its exposure to bioluminescence, acting as a BRET acceptor. For kinases, target engagement in intact live cells allows for biologically relevant measurements of binary target engagement and assessment of their cellular selectivity versus off-target kinases. The high ATP concentration in cells (>1 mM) often leads to a discrepancy between IC^50^ values in cells and cell-free enzymatic IC^50^ data. The latter can be determined using ATP concentrations at the K^m^ for the respective target or at a fixed concentration (Smyth and Collins, 2009;Vasta et al., 2018). Enzymatic IC^50^ data does not account for the cell permeability of the compound, post-translational modifications of the target protein, or regulator protein interactions, which can influence the NanoBRET IC^50^ in intact live cells.

Herein, we developed a NanoBRET cellular target engagement assay for the ATP binding site of SYK GoF variants using a tracer (**8**) designed from a donated chemical probe (MRL-SYKi) for SYK. Initially, we synthesized a suite of NanoBRET tracers for SYK with compounds from Figure 2 and identified tracer **8** as the most potent tracer in enzymatic SYK inhibition assays. NL-tagged SYK GoF variants, which were sufficiently activated as assessed by Y525/526 phosphorylation (SYK(S550Y), SYK(S550F), and SYK(P342T)) generated a BRET signal when combined with tracer **8**. Additionally, in digitonin permeabilized cells, we developed NanoBRET assays for SYK(WT) and all five SYK GoF variants. In the absence of an intact cell assay for SYK(WT), we also optimized a NanoLuc luciferase thermal shift assay (NaLTSA) assay that could detect the binding of MRL-SYKi to SYK(WT) via thermal stabilization. We then determined the cellular IC^50^ of literature SYK inhibitors with NanoBRET assays using these GoF variants and found that the SYK inhibitors were more potent against variants with higher levels of phosphorylation and kinase activity. The SYK donated chemical probe, MRL-SYKi, could decrease the catalytic activity of SYK variants SYK(S550F) and SYK(P342T). Additionally, MRL-SYKi decreased the phosphorylation of SYK(S550Y) GoF variant at Y525/526 in SW480 cells that stably express p.Ser550Tyr.

## 2 Results

### 2.1 Design and synthesis of NanoBRET tracers that potently inhibit SYK

NanoBRET tracers were synthesized from the potent SYK inhibitors entospletinib, R406, and MRL-SYKi. These SYK inhibitors were selected because of their potency and the availability of co-crystal structures (PDB ID: 4PUZ and 3FQS) that identified solvent-exposed sites on the compounds that could be chemically modified (Figure 2) (SGC-Frankfurt;Villaseñor et al., 2009;Currie et al., 2014). The SYK inhibitors were modified to contain a linker region and a BODIPY dye at the solvent-accessible sites identified in their co-crystal structures (Figure 2, yellow circle) (Villaseñor et al., 2009;Currie et al., 2014). In the case of entospletinib, the morpholine group was replaced with a piperazine (**1**), and for R406 the 3,4,5-trimethoxyaniline was replaced with 3-aminophenol. Parent inhibitor **3** was designed from an analogue of entospletinib that displayed weaker enzymatic inhibition of SYK (compound **59**: IC^50^ = 12.3 nM (Currie et al., 2014)). The 3,4-dimethoxyaniline was replaced with 4-(piperazin-1-yl)aniline resulting in an analogue of **1** containing a methylated indazole. Linkers were attached via the generation of an amide bond or an ether linkage, respectively (Supplementary Schemes 1&2). MRL-SYKi already contained a terminal carboxylic acid that appeared to be solvent-exposed, which could be rapidly functionalized with amines to form an amide bond (Supplementary Scheme 3). For tracers **2, 5**, and **7**, we used PEG3 linkers, and for tracers **4, 6**, and **8**, shorter alkyl linkers were used with each of the three parent SYK inhibitors (Table 1). The syntheses of these tracers are reported in the supplementary information. The synthesized tracers were tested in SYK enzymatic inhibition assays at Eurofins DiscoverX. The parent inhibitors **1, 3**, and MRL-SYKi were also tested in the same assay to determine if the modifications made to these inhibitors were detrimental to potency. Additionally, their IC^50^ data was a benchmark of optimal potency for inhibition of SYK – the goal was to produce tracer compounds that did not exhibit more than 10-fold reduction in inhibition from their respective parent inhibitors. The SYK inhibitors were potent (IC^50^ = 16, 97, and 30 nM, for **1, 3**, and MRL-SYKi, respectively), and therefore their synthetic modifications were appropriate for tracer derivatization. Entospletinib-derived tracers **2**, and **4** lacked potent inhibition (IC^50^ = 2400 nM and >10000 nM, respectively) despite their parent inhibitors, **1** and **3**, showing potent enzymatic inhibition. Tracer **6** contained a methylated indazole, and the modification resulted in the abolition of all SYK inhibition (Currie et al., 2014). Tracers **5** and **6**, with the R406-derived inhibitor, also exhibited poor potency in the enzymatic inhibition assays, regardless of the linker length used to attach the inhibitor to the BODIPY dye. Lastly, the tracers synthesized with MRL-SYKi generated the most potent SYK tracers, **7** and **8** (IC^50^ = 470 nM and 110 nM, respectively). These tracers only decreased potency 15.6 and 3.6-fold, respectively, compared to the parent inhibitor MRL-SYKi.

**Table 1.**
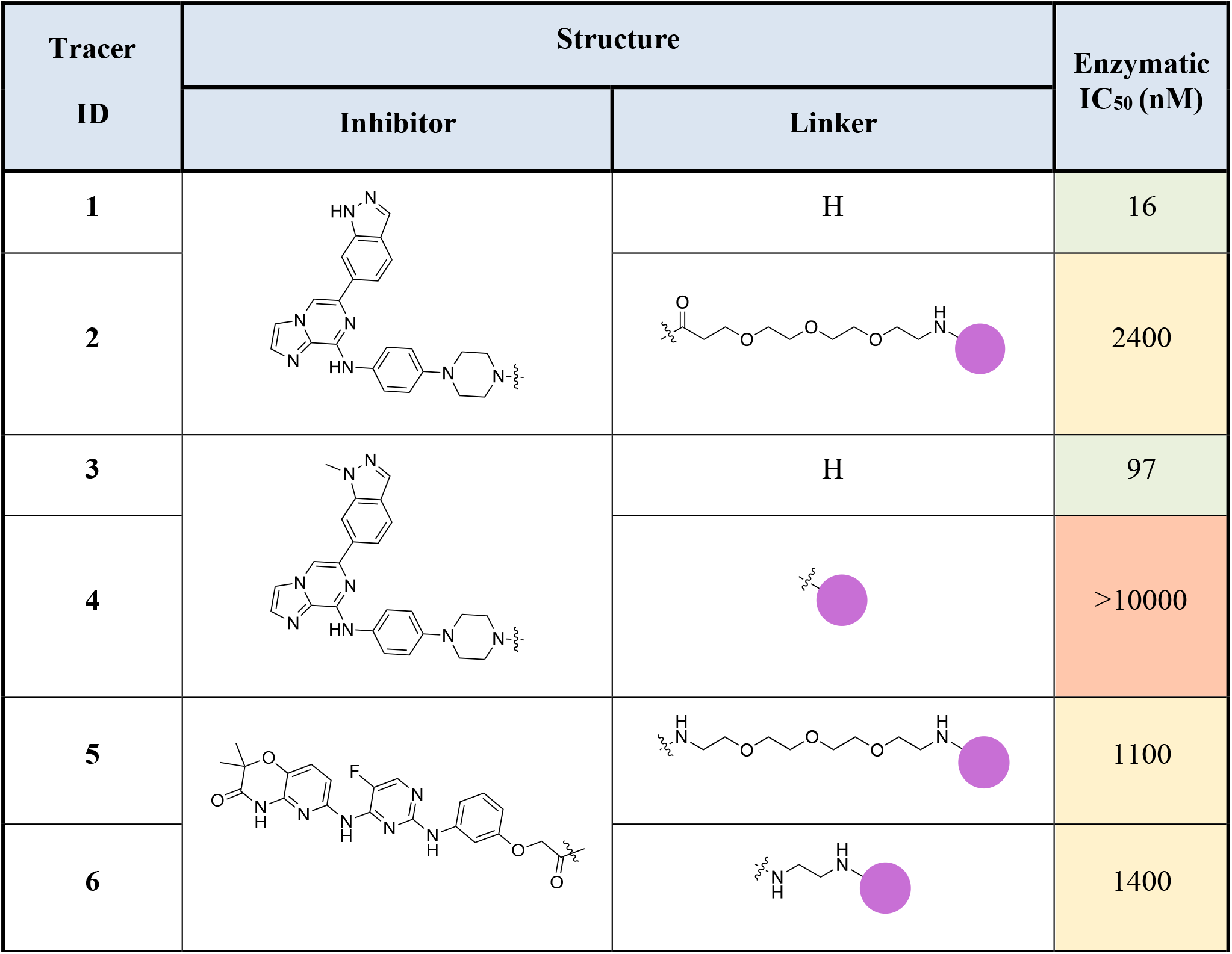
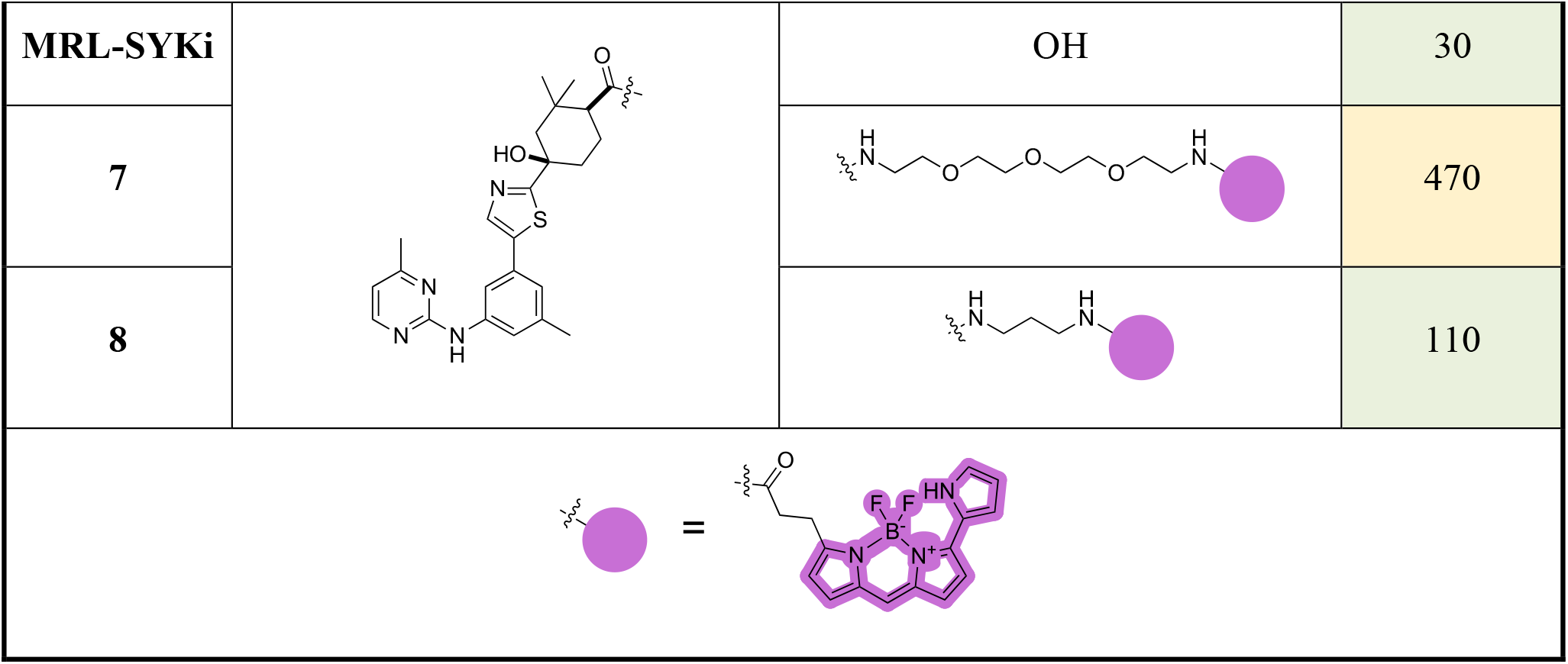
Structures of SYK-derived NanoBRET tracers and corresponding Eurofins DiscoverX enzymatic IC_50_ data for SYK.

| Tracer ID | Structure |  | Enzymatic IC<sub>50</sub> (nM) |
| --- | --- | --- | --- |
|  | Inhibitor | Linker |  |
| **1** |  | H | 16 |
| **2** |  |  | 2400 |
| **3** |  | H | 97 |
| **4** |  |  | >10000 |
| **5** |  |  | 1100 |
| **6** |  |  | 1400 |

| MRL-SYKi |  | OH | 30 |
| --- | --- | --- | --- |
| 7 |  |  | 470 |
| 8 |  |  | 110 |

### 2.2 SYK tracers enable BRET with full-length SYK(WT) after digitonin permeabilization

We performed NanoBRET tracer optimization to determine if the SYK tracers could effectively generate a BRET signal. HEK293 cells, transfected with either full-length kinase domain construct (N- or C-terminal NL tag) of WT SYK, were dosed with tracer concentrations of 0.5, 1, and 2 μM, either alone and with an excess (10 μM) of entospletinib or MRL-SYKi (Supplementary Figure 1&2). None of the tracers (Table 1) generated a BRET signal greater than 5 milliBRET units (mBU) in intact cells expressing either full-length SYK or the kinase domain of SYK, after background subtraction (Supplementary Figure 1&2). Additionally, the BRET signal generated was not disrupted by potent SYK inhibitors, entospletinib or MRL-SYKi, suggesting the observed low signal observed was a background signal and not due to specific binding at the ATP competitive binding site. To determine if the lack of signal was due to the poor permeability of the tracers, we performed the assay with the addition of digitonin, which permeabilizes the cells. Tracer **8** generated a BRET signal of approximately 15 mBU (background subtracted) in cells transfected with SYK(WT) with a C-terminal NL fusion (SYK(WT)-NL) that were permeabilized with digitonin (Supplementary Figure 1F). Under identical conditions no signal was observed with SYK containing an N-terminal NL fusion (NL-SYK(WT)), signifying the positioning of NL at the C-terminus was essential to generate a BRET signal. This result indicated that upon permeabilizing the cells, **8** could bind to the ATP binding pocket of SYK and the BODIPY dye was an appropriate distance from NL to generate a BRET signal. Tracers **5, 6**, and **7** also generated a BRET signal >10 mBU (background subtracted) when the cells were permeabilized with digitonin. Tracer **2** only generated a BRET signal window of 4 mBU, however, this could be competed away with entospletinib. Additionally, tracer **4**, which does not bind to SYK in enzymatic inhibition assays, did not generate a BRET signal. Next, N- and C-terminally-tagged NL SYK kinase domain constructs lacking the tandem SH2 domains were prepared. Tracers **7** and **8** failed to generate a BRET signal with or without digitonin (mBU <5) with either SYK kinase domain only construct (Supplementary Figure 2). To enable the NanoBRET assay using tracer **8** with cells permeabilized by digitonin, a tracer dose–response study was performed for each SYK(GoF) variant and SYK(WT) (Figure 3A), followed by tracer titration experiments (Figure 3B and Supplementary Figure 3). We designed SYK GoF variant constructs tagged with NL at the C-terminus corresponding to SYK(S550Y), (S550F), (P342T), (A353T), and (M450I). Initially, we determined the EC^50^ of tracer **8** for each SYK(GoF) variant and SYK(WT). The results are reported as a BRET fold change, which is the BRET ratio for the vehicle-treated sample divided by the BRET for the MRL-SYKi-treated (20 μM) sample. Tracer **8** fixed concentrations were then selected that were below the EC^50^ of the tracer but still generated a BRET signal large enough to give a useable assay-window (>3-fold over background). The resulting optimal fixed tracer concentrations for these assays were identified; SYK(WT): 31 nM, SYK(S550Y): 7.8 nM, SYK(S550F): 31 nM, SYK(P342T): 7.8 nM, SYK(A353T): 15.6 nM, and SYK(M450I): 63 nM. Using the optimal tracer concentrations for guidance, fixed concentrations of tracer **8** were selected and used to calculate the IC^50^ values for MRL-SYKi and rATP (Figure 3B and Supplementary Figure 3&4). rATP binds to SYK(WT) with an IC^50^ value of 16.8 ± 0.4 μM, compared to 25.6 ± 1.9 μM for SYK(S550Y) and 69.8 ± 1.0 μM for SYK(S550F) in this permeabilized cell assay. The IC^50^ values for SYK(S550Y), SYK(S550F), and SYK(P342T) were generally lower than WT, SYK(A353T) and SYK(M450I) for rATP. Intriguingly, when treated with MRL-SYKi, with the exception of SYK(S550F), MRL-SYKi was more potent for SYK(S550Y) (IC^50^ = 2.1 ± 1.0 nM) and SYK(P342T) (IC^50^ = 4.6 ± 2.6 nM) over SYK(WT) (IC^50^ = 10 ± 0 nM), SYK(A353T) (IC^50^ = 10 ± 5 nM) and SYK(M450I) (IC^50^ = 29 ± 6 nM).

**Figure 3.**
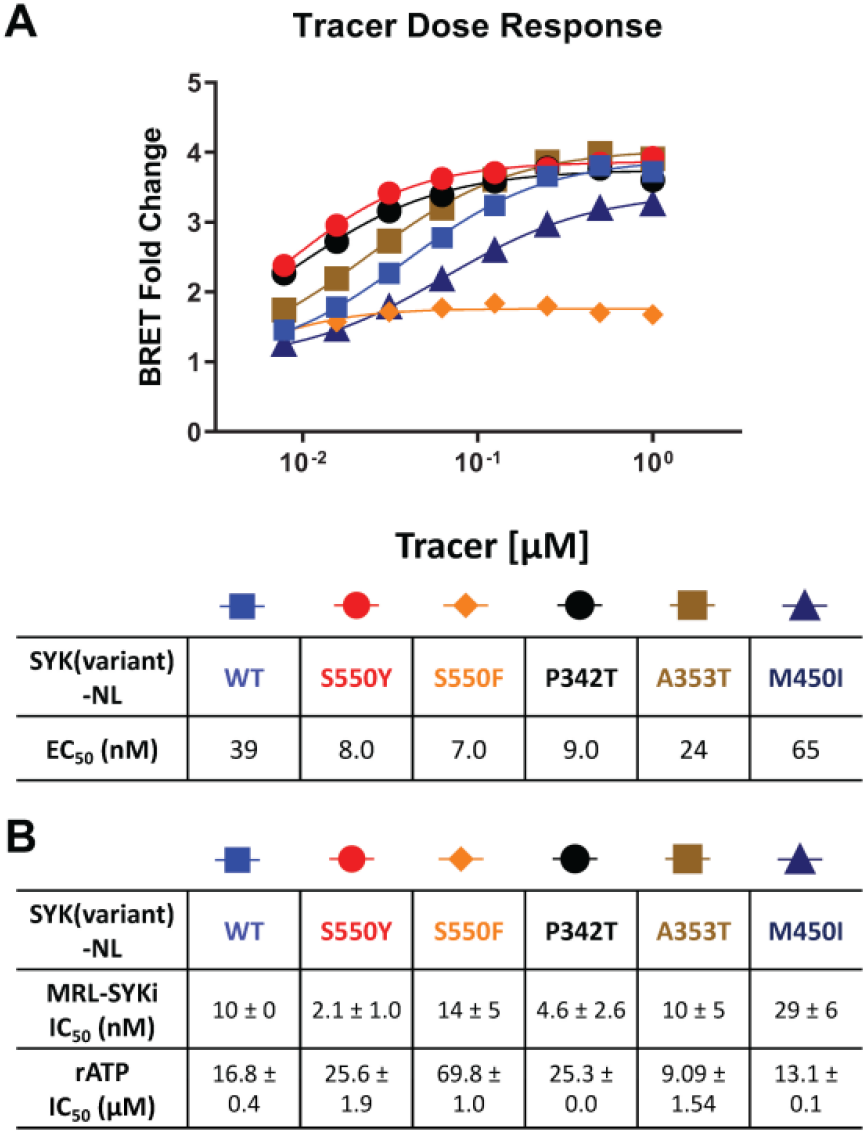
The NanoBRET assay was enabled with tracer **8** in digitonin permeabilized cells and was used to determine the IC_50_ of MRL-SYKi and rATP. (**A**) Competition experiments to generate the EC_50_ of **8** in dose–response format in digitonin permeabilized cells, relative to MRL-SYKi (20 μM) treated cells. Data reported are from a single experiment. (**B**) Dose-response of MRL-SYKi and rATP with SYK(GoF)-NL and SYK(WT)-NL at fixed concentrations of tracer **8**. Data are reported as an average of the IC_50_ ± SEM.

### 2.3 Generation of a SYK GoF variant NanoBRET assay with SYK tracers in intact cells

Next, we used SYK(GoF) variants to determine if tracer **8** could enable a NanoBRET cellular target engagement assay in intact live cells. For tracer optimization experiments, HEK293 cells transfected with each of these constructs were dosed with tracer **8** at 0.5 μM, 1 μM, and 2 μM. We observed BRET signal activation (Figure 4A), that could be competed away with 10 μM of the parent inhibitor MRL-SYKi, indicating this signal was specific to the ATP binding site of SYK. A BRET signal was also observed with tracer **7**, albeit slightly weaker than with **8** (, Figure 4A). Intriguingly, the BRET signal decreased upon permeabilizing cells with digitonin for the SYK(S550Y) variant (Figure 4B and Supplementary Figure 5E). We compared the potency of all SYK tracers in enzymatic SYK inhibition assays versus the BRET signal generated after dosing in SYK(S550Y)-NL transfected HEK293 cells and observed a general trend positively correlating the two when the tracer was dosed at 0.5 μM (Figure 4B). SYK tracers **2, 4, 5**, and **6** also showed the greatest BRET signal at this concentration: SYK(S550Y)-NL >SYK(S550F)-NL >SYK(P342T)-NL (Supplementary Figure 5A–D). There was not a sufficient BRET assay window generated with SYK(A353T) and SYK(M450I) variants, and it could not be competed away with the parent inhibitor. Competition experiments were performed with tracer **8** after transfection of SYK mutants (SYK(S550Y)-NL, SYK(S550F)-NL, and SYK(P342T)-NL) into HEK293 cells (Figure 4C). Tracer **8** was treated in a dose–response format using a top concentration of 3 μM due to solubility issues at >3 μM (Figure 4C). The BRET signal was competed away with an excess of the parent inhibitor MRL-SYKi (10 μM). The cellular EC^50^ of the tracer was <300 nM for each variant ((SYK(S550Y) EC^50^ = 159 nM, (SYK(S550F) EC^50^ = 273 nM, and (SYK(P342T) EC^50^ = 280 nM). To determine the optimal concentration of the tracer for the subsequent NanoBRET assays, a titration of the tracer in the presence of MRL-SYKi in a dose–response format (top concentration 10 μM) was performed (Figure 4D). Based on these dose–response curves, the concentration of tracer **8** for these assays would be fixed at 62.5 nM for SYK(S550Y), and 125 nM for SYK(S550F) and SYK(P342T). These concentrations provided a good assay window while also being below the cellular EC^50^ of the tracer for each variant.

**Figure 4.**
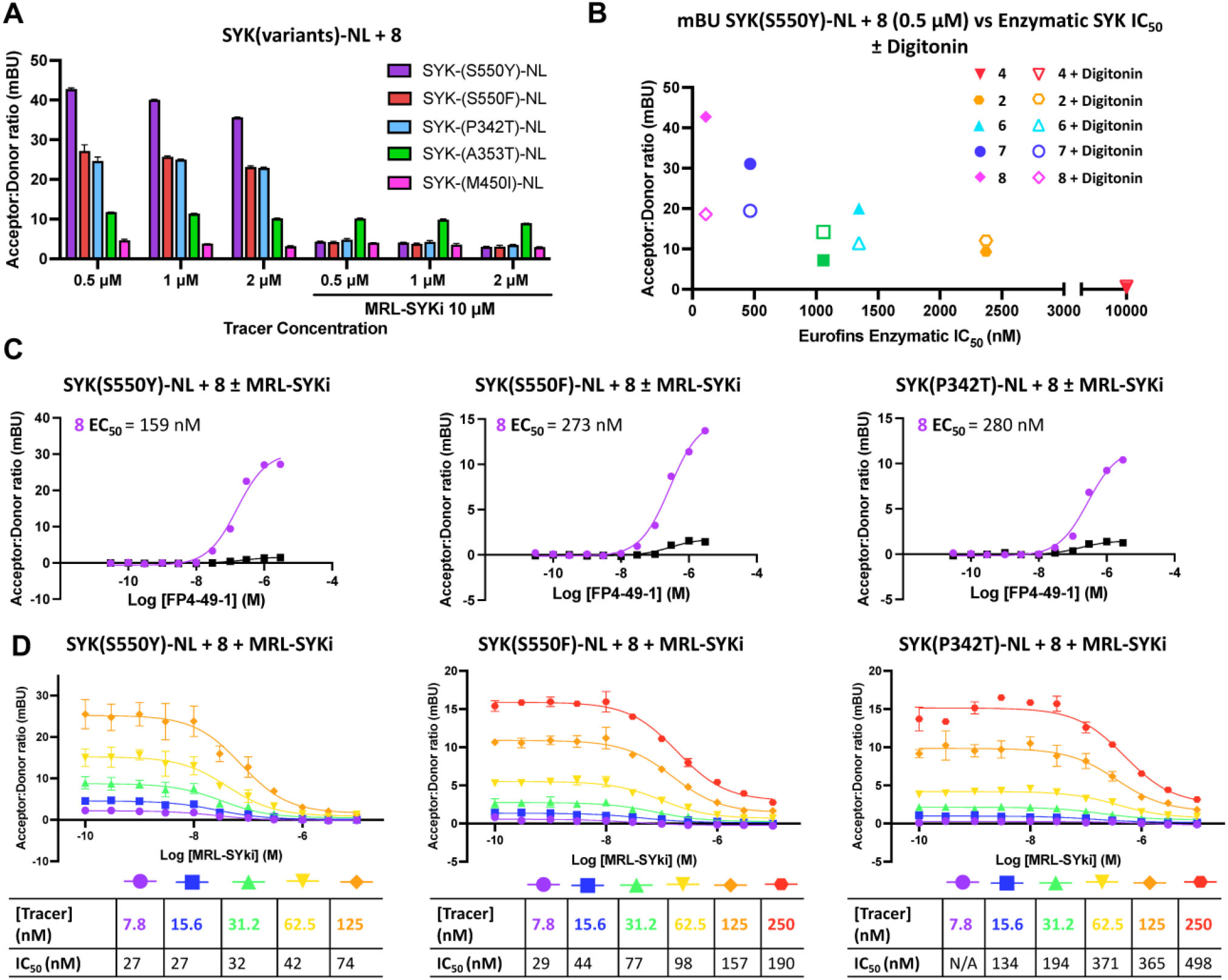
Potent SYK tracers coupled with SYK GoF variants enable BRET (as measured in mBU). (**A**) NanoBRET signal generated with SYK GoF variants and tracer **8**. Data are reported as technical duplicates ± standard deviation. (**B**) The correlation of Eurofins DiscoverX enzymatic inhibition with BRET signal at 0.5 μM of tracer. (**C**) Competition experiments with **8** and SYK(S550Y)-NL (left), SYK(S550F)-NL (middle), and SYK(P342T)-NL (right). Data represent a single experiment. (**D**) Tracer titrations with **8** and SYK(S550Y)-NL (left), SYK(S550F)-NL (middle), and SYK(P342T)-NL (right). Data are reported as duplicates ± standard deviation.

### 2.4 NanoLuciferase thermal shift assay (NaLTSA) determines inhibitor binding to SYK(WT)

Without a NanoBRET cellular target engagement assay for full-length SYK(WT), we optimized a thermal shift assay called NaLTSA. NaLTSA detects an increase in melting temperature (T^m^) via luminescence when the NL-tagged protein is stabilized by inhibitor binding (Dart et al., 2018). HEK293 cells were transfected with SYK(WT)-NL. Following a 20–24 hr incubation, cells were permeabilized with digitonin, treated with compound, heat treated, and finally, luminescence measured. The melting curve for vehicle-treated SYK(WT) corresponded with a melting temperature (T^m^) of 53.8 ± 0.1 °C. MRL-SYKi was added in a dose–response format at concentrations of 1.75, 3.25, 7.5, 15, and 30 μM (Figure 5 and Supplementary Figure 6). A T^m^ of 57.2 ± 0.3 °C was generated at 30 μM MRL-SYKi, leading to a temperature stabilization (ΔT^m^) of 3.5 °C compared to untreated SYK(WT). At 15 μM, a T^m^ of 55.5 ± 0.2 °C was generated with a ΔT^m^ of 1.7 °C compared to untreated SYK(WT), and at 7.5 μM a T^m^ of 54.2 ± 0.5 °C was generated with a ΔT^m^ of 0.5 °C compared to untreated SYK(WT). When comparing the average of the three biological replicates, no change in thermal stabilization temperature above the vehicle was detected at 3.25 or 1.75 μM of MRL-SYKi.

**Figure 5.**
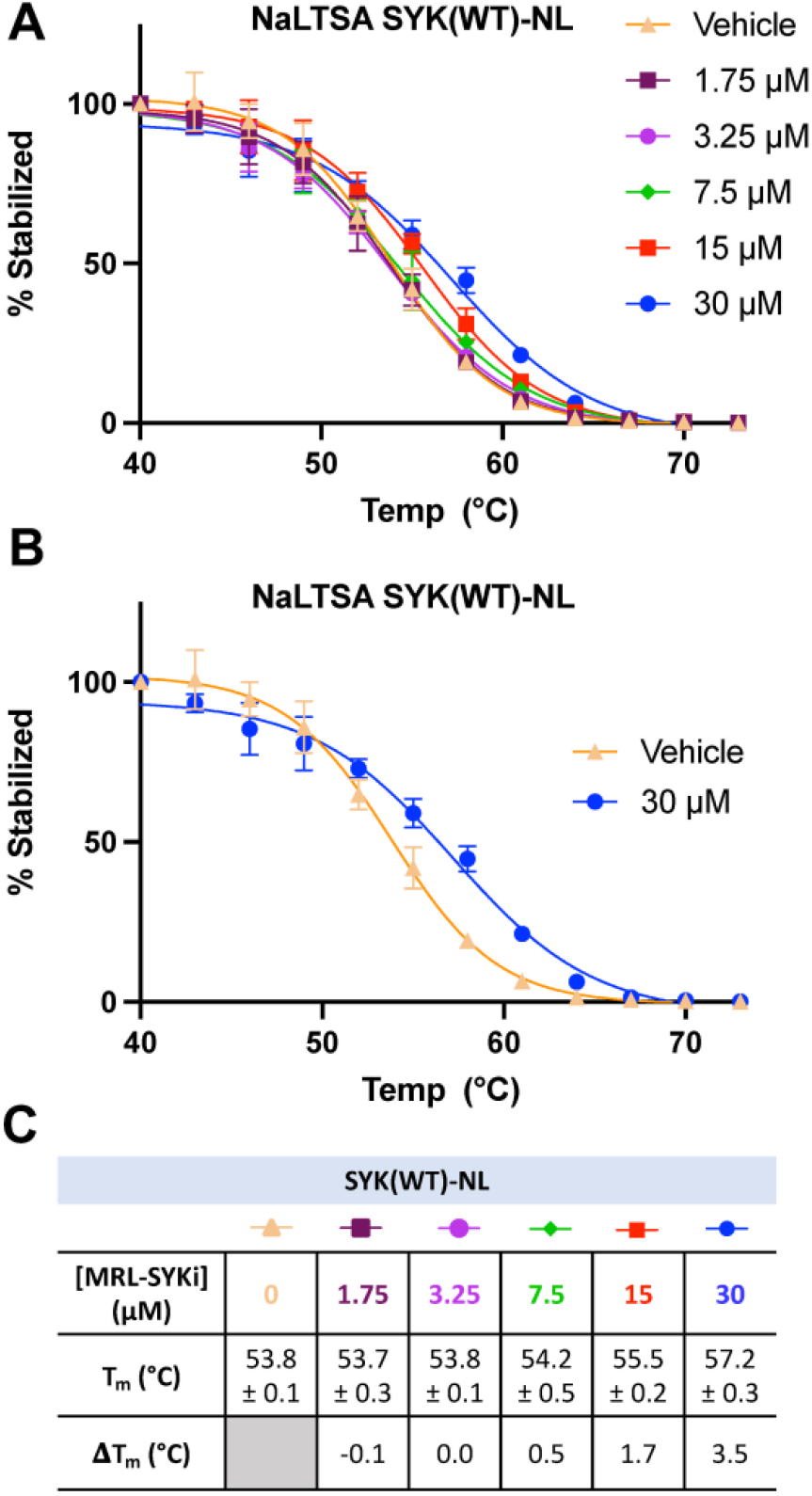
NaLTSA assay for SYK(WT)-NL with MRL-SYKi. (**A**) Melting curves with the addition of MRL-SYKi in dose–response format in HEK293 cells. Data reported are an average from biological triplicates each with two technical replicates. The data was fitted with a Boltzmann Sigmoid equation and error bars indicate the standard deviation (**B**) Visualization of the shift from vehicle with 30 μM of MRL-SYKi from (**A**). **(C**) Melting temperatures (T_m_) at each concentration of MRL-SYKi and the corresponding change in melting temperature (ΔT_m_). Data are reported as an average of the T_m_ ± SEM.

### 2.5 SYK variants have increased phosphorylation and kinase activity

The full-length SYK GoF variants SYK(S550F) and SYK(P342T) along with SYK(WT) were purified from insect cells (Sf9). SYK(S550Y) was unable to be purified due to complications in generating bacmid DNA. The kinase activity of these proteins was assessed using a continuous PhosphoSens sulfonamido-oxine (SOX) fluorophore peptide assay, in the presence or absence of phosphorylated ITAM peptide (Figure 6A&B). Among them, SYK(S550F) exhibited the highest kinase activity, which was further enhanced by the addition of pITAM. SYK(P342T) displayed higher kinase activity than WT SYK, although it was less active than SYK(S550Y). Consistently, the addition of pITAM peptide increased the kinase activity across all variants. Mass spectrometry of the purified proteins confirmed that the SYK GoF variants have higher phosphorylation levels compared to SYK(WT) (Figure 5C). SYK(S550F) had as many as 7 phosphorylation sites (phosphosites), SYK(P342T) had up to 4, while WT SYK was predominantly unphosphorylated, with 2 phosphosites detected. The kinase activity of these proteins was found to correlate well with their phosphorylation levels.

**Figure 6.**
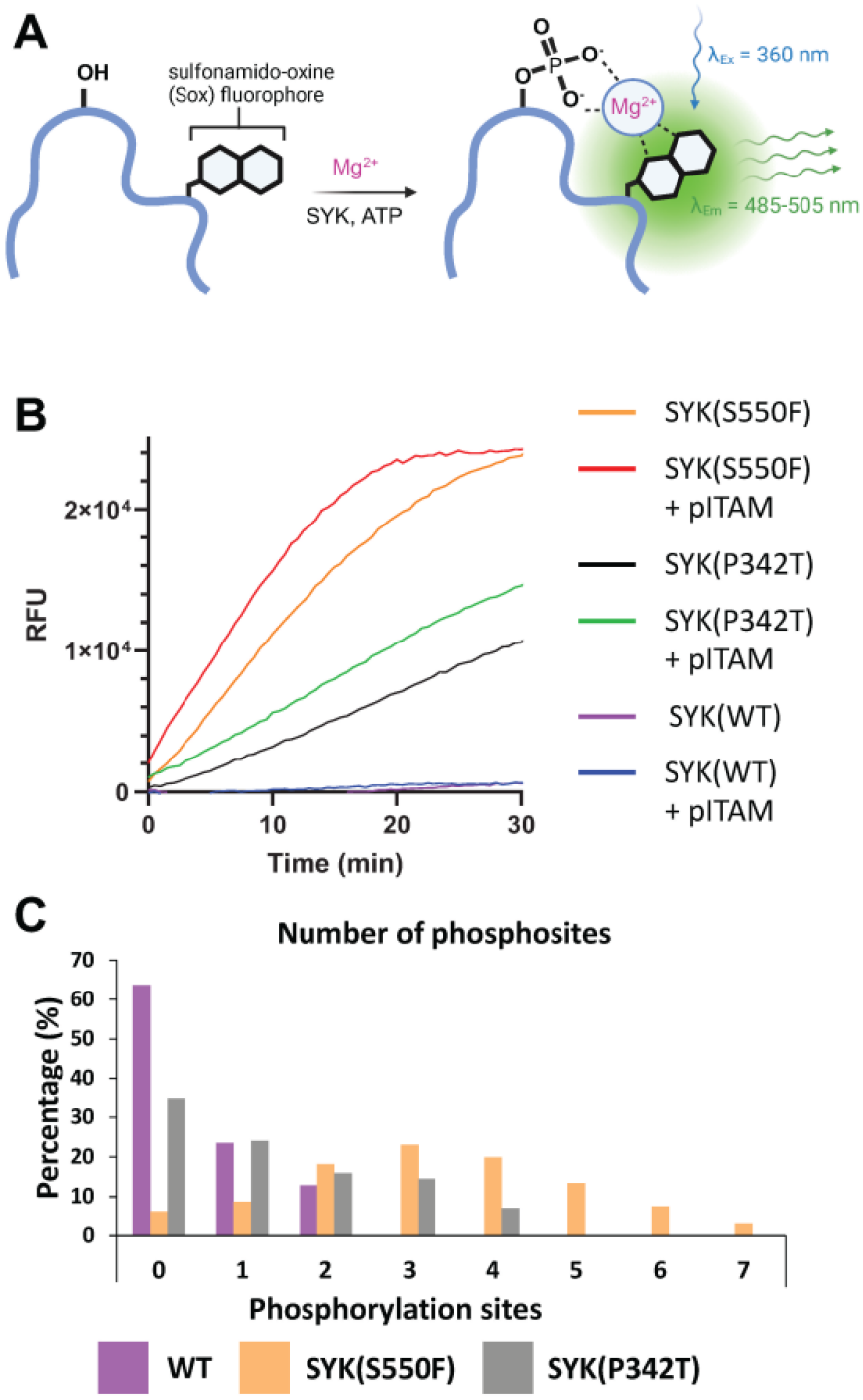
SYK GoF variants are more phosphorylated and have greater catalytic activity than SYK(WT). (**A**) PhosphoSens SOX kinase activity assay for SYK. Upon phosphorylation of the PhosphoSens peptide, chelation of the phosphate with a Mg^2+^ ion enhances fluorescence of the SOX moiety. (**B**) SYK(S550F) and SYK(P342T) GoF variants have greater kinase activity than SYK (WT). Kinase activity was continuously monitored over 30 min with or without the addition of a peptide containing the pITAM motif of FCER1G. (**C**) The number of phosphosites detected within purified SYK(WT), SYK(S550F), and SYK(P342T) proteins, as determined by mass spectrometry.

### 2.6 Inhibitors bind to SYK decreasing catalytic activity and phosphorylation levels

We investigated the cellular potency of literature SYK inhibitors in the NanoBRET cellular target engagement assays developed with the C-terminally tagged NL SYK variants. For compound screening with SYK variants, we used the optimized NanoBRET tracer concentrations determined in Figure 4C&D. We found that these SYK inhibitors bind more potently to SYK(S550Y) than the variants SYK(S550F) and SYK(P342T) (Figure 7A, Supplementary Figure 7). MRL-SYKi is the most selective of these inhibitors in enzymatic kinase inhibition assays and has a target engagement of <500 nM for all three variants in the NanoBRET SYK(GoF) cellular target engagement assays (SGC-Frankfurt). Cerdulatinib and P505-15 were both potent against all three SYK variants. R406 and Entospletinib were the least potent inhibitors in these cellular assays, especially against SYK(P342T). The apparent IC^50^ values generated for MRL-SYKi with tracer **8** (Figure 4D) were plotted in a linearized Cheng-Prusoff analysis to determine the K^D^ value for each (Figure 7B). The K^D^ for SYK variants SYK(S550Y), SYK(S550F), and SYK(P342T) were 27, 46, and 179 nM, respectively. Correspondingly, MRL-SYKi could decrease the catalytic activity of SYK(S550F) and SYK(P342T) in the PhosphoSens SOX peptide assay with purified proteins (Figure 7C, Supplementary Figure 8). Consistent with the findings from the NanoBRET assay, MRL-SYKi inhibited SYK(S550F) more potently than SYK(P342T), with IC^50^ = 63 nM and 95 nM, respectively. Additionally, we showed that tracer **8** inhibits SYK(WT) catalytic activity (Supplementary Figure 8B). Next, we used a colonic epithelial cell line, SW480, stably expressing the SYK(S550Y) variant to determine pSYK levels after MRL-SYKi inhibitor treatment (Figure 7D and Supplementary Figure 9). The cell line was dosed with MRL-SYKi at 100 nM and 10 μM for 20 hours. Both concentrations showed a decrease in pSYK(Y525/526) levels via western blot analysis compared to untreated cells. MRL-SYKi is advised for use in cells at 100 nM based on the reported potency and selectivity data for the inhibitor. Based on this recommendation and our NanoBRET assay potency, the decrease in pSYK(Y625/526) observed at 100 nM is most likely due to the inhibition of SYK in SW480 cells.

**Figure 7.**
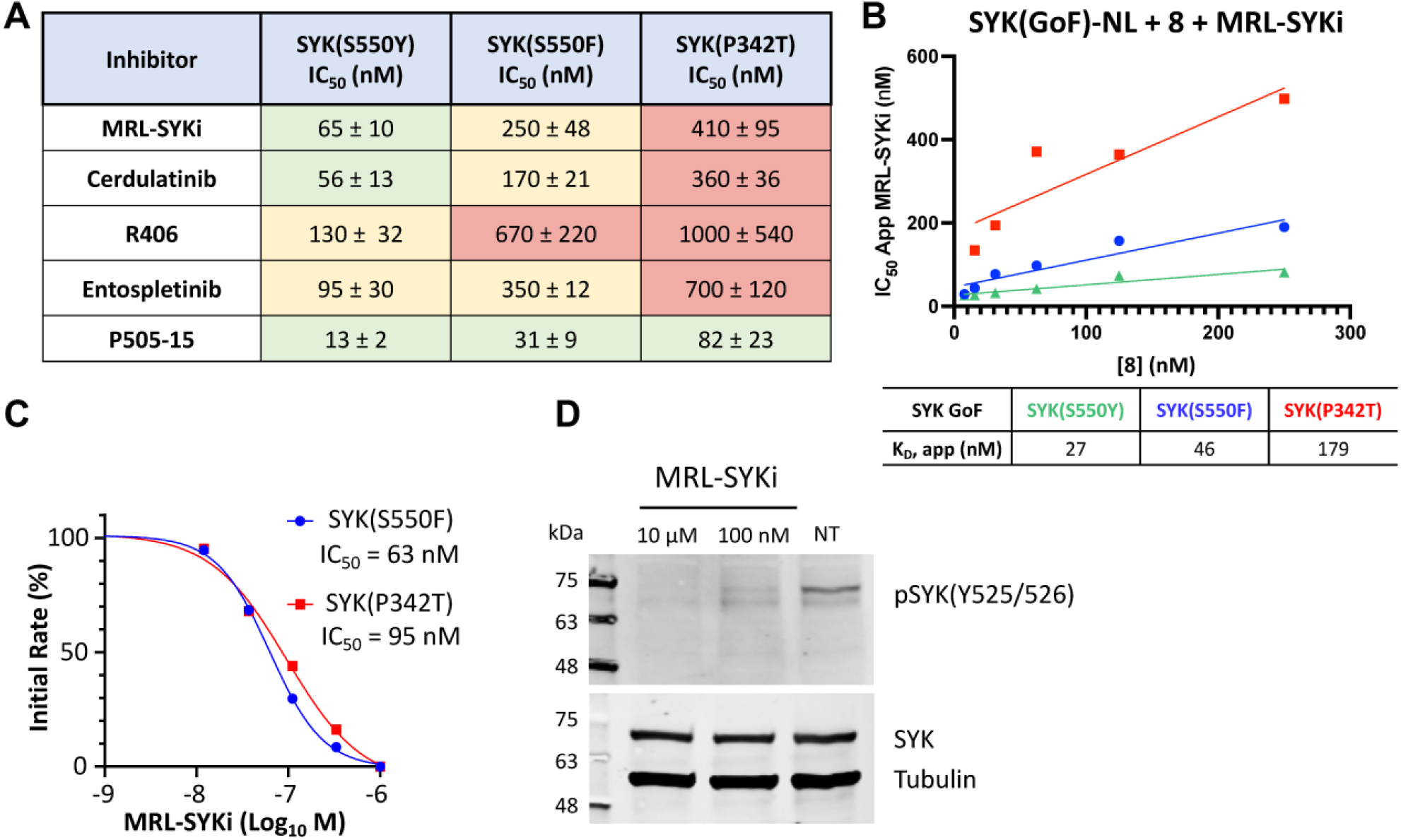
SYK inhibitors bind potently to SYK GoF variants, reducing SYK phosphorylation and SYK catalytic activity. (**A**) IC_50_ data for SYK inhibitors versus SYK GoF variants in HEK293 cells with NanoBRET tracer **8**. Data are reported as an average of triplicate IC_50_ data ± SEM. Data reported are from a single experiment and plotted using a log(inhibitor) vs. response (three-parameter) fit. (**B**) Quantitative analysis of BRET using Cheng-Prusoff relationship for SYK variants in HEK293 cells with NanoBRET tracer **8**. The apparent K_D_ value for MRL-SYKi was determined from the y-intercept by linear regression. Data reported are from a single experiment and plotted in GraphPad Prism with a simple linear regression. (**C**) The kinase activity of SYK(S550F) and SYK(P342T) is inhibited with MRL-SYKi in the PhosphoSens Assay. (**D**) MRL-SYKi decreases pSYK(Y525/526) levels in a SW480 cell line stably expressing SYK(S550Y) after treatment at 100 nM for 20 hrs.

## 3 Discussion

SYK is a target for autoimmune and inflammatory diseases due to its role downstream of activated T cell receptors, B cell receptors, and Fc receptors in immune signaling pathways (Zhou et al., 2023). The activity of SYK in cells is tightly regulated through phosphorylation, making the development of cellular target engagement assays targeting the ATP binding site challenging (Keshvara et al., 1998;Mócsai et al., 2010;Cottat et al., 2017;Kostyak et al., 2022). When SYK is activated in cells via tandem SH2 binding to pITAMs on immunoreceptors or autophosphorylation and phosphorylation via LYN kinase, SYK is released from its autoinhibited state (Bohnenberger et al., 2011;Grädler et al., 2013). Recently, five SYK GoF variants were identified in six patients that lead to the clinical manifestation of immune dysregulation, organ inflammation, and a predisposition for lymphoma (Wang et al., 2021). Despite SYK’s importance in disease and the existence of potent and selective SYK inhibitors, there was previously no assay to determine direct target engagement of SYK in intact live cells.

We designed the first NanoBRET cellular target engagement assays in intact live cells for three SYK GoF variants, SYK(S550Y), SYK(S550F), and SYK(P342T), that have increased phosphorylation levels and catalytic activity (Figure 6). We hypothesized that the GoF variants, due to their increased activity, allow the ATP binding site within SYK’s kinase domain to be accessed by inhibitors (Figure 2). Additionally, the use of the SYK GoF variants in combination with potent SYK-targeted NanoBRET tracers led to a greatly improved assay window. Generally, we found the tracers that exhibited an IC^50^ of <500 nM (tracers **7** and **8**) in enzymatic assays resulted in the largest assay window (Table 1). We were unable to generate a NanoBRET cellular target engagement assay for SYK(WT) or the variants SYK(A353T) and SYK(M450I) in intact live cells, likely due to their lower phosphorylation levels, as described by Wang et al, that result in lowered kinase activity (Wang et al., 2021). However, we found that permeabilizing the cells with digitonin generated a BRET signal with SYK(WT), as well as the five SYK GoF variants, when using our optimized SYK tracer **8** (Figure 3). Intriguingly, the BRET signal observed in digitonin permeabilized cells was much less than in intact live cells for SYK GoF variants SYK(S550Y), SYK(S550F), and SYK(P342T) (Figure 4B and Supplementary Figure 3&4). Additionally, we also optimized a binding assay for SYK(WT) using NaLTSA, which was used to demonstrate a temperature shift due to thermal stabilization of the NL-tagged SYK upon MRL-SYKi binding (Figure 5). In the NaLTSA assay, cells are permeabilized with digitonin before treatment with the compound, and then heated. Assays that employ digitonin to permeabilize cells appear to overcome the autoinhibition of SYK(WT) and allow the detection of binding at the ATP binding site; however, the reason for this is unclear. The NaLTSA assay registered a small but significant ΔT^m^ upon MRL-SYKi binding to SYK(WT) at concentrations of ≥15 μM, despite its potent inhibition of SYK in the NanoBRET assay (SYK(S550Y) IC^50^ = 65 ± 10 nM) and in enzymatic inhibition assays (SGC-Frankfurt). This suggests that while the NaLTSA assay can be used to determine inhibitor binding to SYK(WT), it is likely that only high-potency inhibitors will be identified using this method due to the small temperature shift observed for MRL-SYKi.

We characterized a suite of SYK inhibitors, which have reported nanomolar enzymatic inhibition, in the SYK GoF variant NanoBRET assays and confirmed they bind potently to each of the three SYK GoF variants (Figure 7). Generally, we observed that inhibitors more potently bound to SYK(S550Y), which has increased phosphorylation levels and is therefore likely to have a more exposed ATP binding site for inhibitor binding (Wang et al., 2021). The SYK inhibitors bind more potently to SYK variants with increased phosphorylation levels (SYK(S550Y) >SYK(S550F) >SYK(P342T)). The phosphorylation levels were quantified via western blot by Wang et al. This group showed that SYK(WT), SYK(A353T), and SYK(M450I) have minimal to no phosphorylation. We also compared the phosphorylation levels of recombinant purified SYK(WT), SYK(S550F), and SYK(P342T) and found a higher level of phosphorylation with the S550F and P342T variants compared to SYK(WT) (Figure 6). Furthermore, we showed that the catalytic activity of SYK was greater with SYK GoF variants compared to SYK(WT), and the magnitude of catalytic activity correlated with the levels of SYK phosphorylation (Figure 6). The rank order of SYK inhibitor potency based on the mean IC^50^ for SYK(S550Y) is 1) P505-15 (IC^50^ = 13 ± 2 nM), 2) MRL-SYKi (IC^50^ = 65 ± 10 nM), 3) cerdulatinib (IC^50^ = 56 ± 13 nM), 4) entospletinib (IC^50^ = 95 ± 30 nM), and 5) R406 (IC^50^ = 130 ± 32 nM) (Figure 7A). This rank order of potency is also observed for the S550F and the P342T variants.

We then investigated the effect on the catalytic activity of SYK(S550F) and SYK(P342T) upon the addition of the SYK donated chemical probe, MRL-SYKi. MRL-SYKi decreased the catalytic activity of the SYK GoF variants in a dose–response manner (IC^50^ = 63 nM and 95 nM, respectively) (Figure 7C). This data also indicates that potent inhibition of SYK GoF variants is correlated with their higher levels of phosphorylation and activity. Finally, we showed that MRL-SYKi could decrease SYK phosphorylation (pSYK(Y525/526)) in a colonic epithelial cell line, SW480, stably expressing SYK(S550Y) (Figure 7D). MRL-SYKi decreased pSYK(Y525/526) levels at 100 nM, and to a greater extent at 10 μM, after 20 hours of treatment. The donated chemical probe, MRL-SYKi is selective for SYK at 100 nM in biochemical selectivity assays, supporting that the decrease in pSYK(Y525/526) that is observed at 100 nM is a direct result of SYK inhibition in these cells.

Overall, we have optimized a novel NanoBRET cellular target engagement assay for three SYK GoF variants; (SYK(S550Y), SYK(S550F), and SYK(P342T). These three variants lead to the clinical presentation of intestinal, skin, and joint inflammation, recurrent infections, and hypogammaglobulinemia in infants at 2 weeks of age (Wang et al., 2021). Our assays can be used to assess the target engagement of SYK inhibitors at the ATP binding site and adds kinase assays that can be employed to profile the overall cellular selectivity of kinase inhibitors. Additionally, we demonstrated that SYK inhibitors, such as MRL-SYKi, bind potently to all three SYK GoF variants and can decrease both the catalytic activity and the phosphorylation levels of SYK.

## 4 Materials and Methods

### 4.1 Cell Culture

Human embryonic kidney (HEK293) cells were obtained from ATCC and cultured in Dulbecco’s Modified Eagle’s medium (DMEM, Gibco, #11965092) supplemented with 10% (v/v) fetal bovine serum (FBS, VWR Avantor Seradigm, #97068-085). HEK293 cells were incubated in 5% CO^2^ at 37 °C and passaged every 72 hr with trypsin (Gibco, #25300054) not allowing them to reach confluency.

### 4.2 Enzymatic Assays

Eurofins kinase enzymatic radiometric assays were executed at the K^m^ value for ATP in dose–response (9-pt curve) format for SYK, included in Table 1 as enzymatic IC^50^ values. Details of the substrate used, protein constructs, controls, and assay protocol for these kinase assays are available at the Eurofins website: https://www.eurofinsdiscoveryservices.com.

### 4.3 General Information for NanoBRET Assays

HEK293 cells were transfected with constructs of SYK (WT, KD, and GoF variants) tagged with NL on the N- or C-terminus as previously described (Wells et al., 2021). Constructs for NanoBRET measurements of SYK(WT) (NL-SYK(WT) and SYK(WT)-NL) and SYK GoF variants (SYK(S550Y)-NL, SYK(S550F)-NL, SYK(P342T)-NL, SYK(M450I)-NL, and SYK(A353T)-NL) were kindly provided by Promega. NL orientations used in the respective assays are indicated. Cells were plated in 96-well tissue culture-treated plates (Corning, #3917) at a cell density of 2 × 10^5^ cells/mL, with a total volume of 100 μL per well in DMEM + 10% FBS. After 16 h, the media was aspirated from the plate and replaced with room temperature Opti-MEM without phenol red (Gibco, #11058021), 100 μL in no tracer wells; 95 μL in tracer only wells; 90 μL in no tracer wells and with digitonin treated plates, 90 μL in no tracer wells; 85 μL in tracer only wells; 80 μL in no tracer wells. Plates were incubated for 25 min if digitonin was added or for 2 hr for the intact cell NanoBRET assay. NanoBRET plates were read after the addition of 50 μL of a stock solution (3X) containing NanoBRET NanoGlo substrate (Promega, #N2161), extracellular NanoLuc inhibitor (Promega, #N2161) when reading assay plates performed in intact live cells, and Opti-MEM without phenol red (Gibco, #11058021). For intact live cells in a 96-well plate, the 3X stock solution was prepared with 30 μL of NanoBRET NanoGlo substrate, 10 μL of extracellular NanoLuc inhibitor, and 4960 μL of Opti-MEM without phenol red. For digitonin permeabilized live cells in a 96-well plate, the 3X stock solution was prepared with 30 μL of NanoBRET NanoGlo substrate and 4970 μL of Opti-MEM without phenol red. Raw milliBRET units (mBU) were read on a GloMax Discover system (Promega) with a donor emission wavelength of 450 nm and an acceptor emission wavelength of 600 nm. mBU were calculated by dividing the acceptor emission values (600 nm) by the donor emission values (450 nm). For NanoBRET studies with SYK, background corrected mBU were calculated by subtracting the no tracer wells from the tracer-containing wells and multiplying by 1000.

### 4.3.1 Tracer Optimization Studies

HEK293 cells were transfected with constructs of SYK(WT) or SYK(KD) tagged with NL on the C- or N-terminus and SYK GoF variants tagged with NL on the C-terminus. Tracer **8** (20X) was prepared from a 400 μM stock solution in DMSO with tracer dilution buffer (Promega, #N291B) and 20% DMSO, with a final assay plate concentration of 1% DMSO. 5 μL of **8** (20X) was added to each well, with exception of the no tracer control wells, with final plate concentrations of 0.5, 1, and 2 μM. 10 μL of digitonin (Fisher, 10X) was added to specific wells to permeabilize the cells. The 10X digitonin stock was prepared by diluting a 40X DMSO solution of digitonin with room temperature Opti-MEM without phenol red, for a final assay plate concentration of 1% DMSO. SYK inhibitor stock solutions of MRL-SYKi (10X) were prepared from 10 mM DMSO stock solutions with room temperature Opti-MEM without phenol red, for final assay plate concentrations of 10 μM. Plates containing digitonin were incubated for 25 min and non-digitonin containing plates were incubated for 2 h. Two technical replicates were plotted in a bar chart with the standard deviation represented as error bars on Graphpad Prism.

### 4.3.2 Tracer Competition Experiments

HEK293 cells were transfected with SYK(S550Y)-NL, SYK(S550F)-NL, and SYK(P342T)-NL). MRL-SYKi was used to compete away the NanoBRET signal produced with tracer **8**. These compounds displace the tracer due to occupancy of the same binding site. Tracer **8** was prepared in DMSO and DMEM without phenol red in 11-point dose-response format with 3 μM as the top concentration, with a final assay plate concentration of 1% DMSO. 10X stock solutions of MRL-SYKi were prepared from 10 mM DMSO stock solution with room temperature Opti-MEM without phenol red, for final assay plate concentrations of 10 μM. 10 μL of **8** (10X) was added to each well, with the exception of the no tracer control wells, and 10 μL of 10X solutions of MRL-SYKi was added to wells for tracer competition. One biological replicate was plotted in GraphPad Prism with [inhibitor] vs. response (three parameters).

### 4.3.3 Tracer Titration

HEK293 cells were transfected with SYK(S550Y)-NL, SYK(S550F)-NL, and SYK(P342T)-NL). MRL-SYKi was tested in 11-point dose–response format with a top concentration of 10 μM. Tracer **8** (20X) stock solutions were prepared in tracer dilution buffer (Promega, #N291B) and 20% DMSO, for final assay plate concentrations of 7.8 nM, 15.6 nM, 21.3 nM, 62.5 nM, and 125 nM for SYK(S550Y)-NL or 7.8 nM, 15.6 nM, 21.3 nM, 62.5 nM, 125 nM, and 250 nM for SYK(S550F)-NL and SYK(P342T)-NL. The final assay plate concentrations included 1% DMSO. 5 μL of **8** (20X) was added to each well, except the no tracer control wells, and 10 μL of 10X threefold diluted solutions of MRL-SYKi was added to wells. Two biological replicates were plotted in GraphPad Prism with log[inhibitor] vs. response (three parameters) with the error bars indicating standard deviation.

### 4.3.4 SYK NanoBRET Assays - Inhibitor Screening

HEK293 cells were transfected with SYK(S550Y)-NL, SYK(S550F)-NL, and SYK(P342T)-NL). Based on the tracer titration results, assays were carried out as described by the manufacturer using 62.5 nM of tracer **8** for SYK(S550Y)-NL, 125 nM of tracer **8** for SYK(S550F)-NL and SYK(P342T)-NL. 20X stock solutions of the respective tracers were prepared in tracer dilution buffer (Promega, #N291B) and 20% DMSO for a final assay plate concentration of 62.5 nM (SYK(S550Y)-NL) or 125 nM (SYK(S550F)-NL and SYK(P342T)-NL), with a final assay plate concentration of 1% DMSO. Compounds were tested in 11-point dose–response format with a top concentration of 10 μM. Data are reported as IC50 ± standard error mean (SEM). One biological replicate was plotted in GraphPad Prism with log[inhibitor] vs. response (three parameters), the three IC50’s were averaged and the SEM was calculated.

### 4.4 Purification of Proteins

Full-length human SYK(WT) DNA was custom synthesized (GenScript) as a codon optimized form for expression in Sf9 cells. N-terminal 6His and GST tags were introduced, followed by a TEV protease cleavage site, and cloned into pFastbac1. SYK(S550F) and SYK(P342T) were generated by site-directed mutagenesis of SYK(WT) using the NEB Q5 site directed mutagenesis kit. Plasmids were transformed into *E. coli* DH10Bac for bacmid generation. Recombinant bacmid DNA was used to transfect Sf9 cells to generate viral stocks. For large-scale protein production, 2L of Sf9 cells at 2 x 10^6^ cells/ml were infected with a P2 virus stock and allowed to incubate with shaking for 3 days at 27 °C. Harvested cells were lysed in 200 ml lysis buffer containing 50 mM HEPES (pH 7.5), 500 mM NaCl, 10 mM imidazole, 10% glycerol, 1 mM TCEP. The protein was bound to equilibrated Ni-iminodiacetic acid resin for 1 hr before two batch washes in lysis buffer. Beads were subsequently loaded onto a drip column, followed by washing with lysis buffer containing 30 mM imidazole, before finally eluting bound protein with lysis buffer containing 300 mM imidazole. The 6His and GST tags were removed by the addition of a 1:10 mass ratio of 6His-TEV protease while undergoing dialysis in lysis buffer lacking imidazole for 16 hr at 4 °C. TEV protease and the cleaved 6His tag was subsequently removed with Ni-iminodiacetic acid resin equilibrated in lysis buffer. SYK protein was further purified by size exclusion chromatography in buffer containing 10 mM HEPES (pH 7.5), 150 mM NaCl, 10% glycerol, 1 mM TCEP, prior to concentrating to 1-7 mg/ml. The molecular mass of purified protein was confirmed by intact mass spectrometry.

### 4.5 Mass Spectrometry

Proteins were diluted to 1 mg/ml in buffer containing 50 mM HEPES (pH 7.5) and 200 mM NaCl before being diluted 1:20 in 0.1% formic acid. Reversed-phase chromatography was performed in-line prior to mass spectrometry using an Agilent 1100 HPLC system (Agilent Technologies inc. – Palo Alto, CA, USA). Protein samples in formic acid were injected (50 μL) on to a 2.1 mm × 12.5 mm Zorbax 5um 300SB-C3 guard column maintained at 40 °C. The employed solvent system comprised of 0.1% formic acid in ultra-high purity water (solvent A) and 0.1% formic acid in methanol (solvent B). Chromatography was performed as follows: Initial conditions were 90 % A and 10 % B and a flow rate of 1.0 ml/min. After 15 s at 10 % B, a two-stage linear gradient from 10 % B to 80 % B was applied, over 45 s and then from 80% B to 95% B over 3 s. Elution then proceeded isocratically at 95 % B for 1 min 12 s, followed by equilibration at initial conditions for a further 45 s. Protein intact mass was determined using a 1969 MSD-ToF electrospray ionisation orthogonal time-of-flight mass spectrometer (Agilent Technologies). The instrument was configured with the standard ESI source and operated in positive ion mode. The ion source was operated with the capillary voltage at 4000 V, nebulizer pressure at 60 psig, drying gas at 350°C and drying gas flow rate at 12 L/min.

### 4.6 SOX Peptide Activity Assay

SYK kinase activity was evaluated using a continuous kinase assay using a PhosphoSens (AssayQuant) SOX peptide, according to the protocol provided, with modifications. The assays were conducted in 20 μl volumes within 384-well black non-binding plates. Both SYK(WT) and SYK GoF mutant proteins were tested at a final concentration of either 2 nM (Figure 6B, 7C and Supplementary Figure 8A) or 8 nM (Supplementary Figure 8B) in assay buffer containing 54 mM HEPES (pH 7.5), 1.2 mM DTT, 0.55 mM EGTA, 0.012% Brij-35, 10 mM MgCl^2^, 1% glycerol, and 200 μg/ml BSA. To this mixture, 10 μM PhosphoSens peptide (AQT0794) was added. Where indicated, the reaction was supplemented with 10 μM of a peptide containing the dually phosphorylated ITAM motif of FCER1G [DGV(pY)TGLSTRNQET(pY)ETLKH], which was custom synthesized by LifeTein. The assay mixture was incubated, together with inhibitors when noted, for 30 min. Prior to initiating the kinase reaction with 1 mM ATP, the assay mixture was incubated for 30 minutes at room temperature. Fluorescence readings were taken at 30-second intervals for a duration ranging from 30 minutes to an hour, using a PheraStar FX device set to kinetic mode, employing an appropriate optics module (FI 350 460). The data was represented as a plot of relative fluorescence over time, deducting the fluorescence of control reactions (which lacked kinase) from the fluorescence recorded after introducing the kinase. The IC^50^ values for MRL-SYKi were calculated by measuring the initial rate of kinase activity at various inhibitor concentrations, followed by performing a four-parameter non-linear regression analysis (GraphPad).

### 4.7 NaLTSA Assay

All NaLTSA assays were performed using an altered version of existing published protocols (Dart et al., 2018). A transfection complex of DNA at 10 μg/mL was created, consisting of 9 μg/mL diluent DNA and 1 μg/mL SYK NL fusion DNA in serum-free Opti-MEM without phenol red (Gibco, #11058021). The SYK NL fusion DNA was encoded in a pFC32K expression vector (Promega) with a C terminal NL attachment. Promega kindly provided the full-length human construct for SYK (SYK-NL). After the DNA solution was made, 30 μL of FuGENE HD (Promega) was added for every mL of DNA solution. The transfection solution was immediately vortexed and incubated at room temperature for 20 minutes to allow for lipid:DNA complex formation. The transfection complex was then mixed with a 20X volume solution of HEK293 cells in DMEM/FBS to arrive at a final concentration of 200,000 cells/mL. The transfected cell solution was then plated onto T75 plates and incubated at 37°C in 5% CO^2^. After a 20–24-hour incubation, the cells were washed with 8 mL of PBS and harvested with 0.05% Trypsin-EDTA (GIBCO). The trypsinized cells were resuspended in 9 mL of DMEM and added to a 15 mL centrifuge tube, before being centrifuged at 1200x rpm for five minutes. The supernatant was aspirated, and the pelleted cells were resuspended and then diluted to a concentration of 2.25 x 10^5^ in Opti-MEM without phenol red. Next, a protease inhibitor cocktail (Promega, #G6521), reconstituted in DMSO, was used to make a 100X stock solution with all test compounds. 1 μL of the 100X protease inhibitor/test compound solution was added to each well in a 3 x 32-well PCR Reaction Plate. Then, 10 x 500 μg/mL stock solution of digitonin (Sigma) was made in Opti-MEM, and 10 μL of this solution was added to each well of the PCR reaction plate. Next, 89 μL of the cell solution was added to each well to bring the total volume to 100 μL per well. The solution was mixed by pipetting. The plates were then covered with an adhesive seal and incubated at room temperature for 30 minutes. After the incubation, the plates were placed in a thermal cycler (Applied Biosystems ProFlex PCR System, Thermo Fisher) and treated with temperatures ranging from 40-73°C with 3-degree increments for a period of 3 minutes. The reaction was cooled at room temperature for 3 minutes, which was followed by the addition of 25 μL of a 5X solution of NanoGlo substrate (Promega, #N2161). The cells were then transferred to a white non-binding surface (NBS) 96-well plate (Corning, #3917). The plates’ total luminescence was then read using a GloMax Discover luminometer (Promega). To generate T^m^ values, the raw luminescence data was converted to percent stabilized by normalizing it to the 40 °C luminescence values. The resulting values were then fitted to gather apparent T^m^ values using the Boltzmann Sigmoid Equation. All NaLTSA experiments were run in technical duplicate and repeated independently three times.

### 4.8 SYK Phosphorylation Western Blots

SW480 cells stably expressing S550Y variant were maintained as previously described (Wang et al., 2021). For SYK S550Y inhibition, cells were seeded in 12-well plate, at 10^6^ cells /well. The next day, medium was replaced and MRL-SYK inhibitor was added at 10 μM and 100 nM for 20h. Protein lysates were prepared and analyzed on Western blot as previously described (Wang et al., 2021).

## 6 Author Contributions

Conceptualization, FMB; Formal Analysis, JLC, JDV, VLK, AM, MTB, SCDD, SC-K, ESB, SH, ID, FMB; Funding Acquisition, ADA; Investigation, JLC, JDV, VLK, AM, MTB, SCDD, SC-K, ESB, SH, FMB; Methodology, JLC, JDV, VLK, AM, MTB, SCDD, SC-K, ESB, SH, ID, FMB; Project administration, ID, MBR, ADA, FMB; Supervision, ID, MBR, ADA, FMB; Validation, JLC, JDV, VLK, AM, MTB, SCDD, SC-K, ESB, SH, ID, FMB; Visualization, FMB; Writing – original draft, FMB; Writing – review and editing, JLC, JDV, VLK, SC-K, ID, MBR, ADA, FMB.

## 8 Acknowledgments

We thank the SGC Frankfurt for providing the donated chemical probe, MRL-SYKi. Biorender was used to generate the images in Figure 1, and Figure 6.

## 9 Data Availability Statement

All of the reagents generated are available from the corresponding author.

