## Supplementary Material for "Development of SYK NanoBRET Cellular Target Engagement Assays for Gain–of–Function Variants"

#### Table of Contents

**1 Supplementary Figures and Tables**

**1.1 Supplementary Figures ........ S2–S9**

**1.2 Chemistry Experimental ........ S9**

**1.3 General Information for Chemical Synthesis ........ S9**

**1.4 Abbreviations ........ S9**

**1.5 Chemistry Schemes ........ S10–S11**

**1.6 Chemistry Experimental Data ........ S12–S20**

**1.7 Purity Traces and Spectra ........ S21–S38**

### 1 Supplementary Figures and Tables

#### 1.1 Supplementary Figures

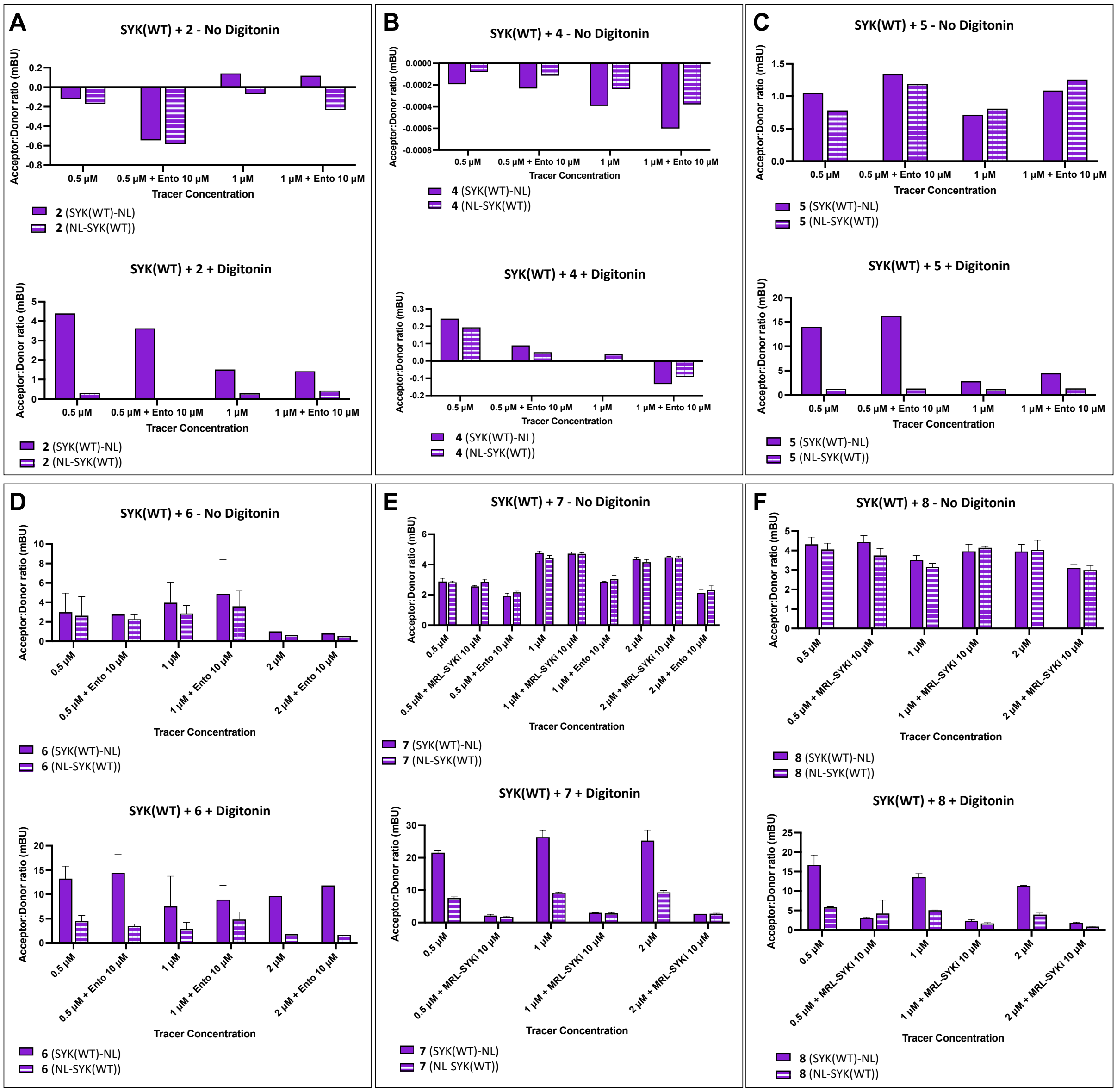


**Supplementary Figure 1.** SYK NanoBRET tracers **2**, **5**, **6**, **7**, and **8** generate a BRET signal (measured in mBU) with SYK(WT)-NL upon digitonin permeabilization (**A**, **C**–**F**), but not without digitonin. (**B**) Tracer **4** does not generate a BRET signal ± digitonin. Data reported are from a single experiment (**A**–**C**). Data are reported in biological duplicate ± standard deviation (**D**). Data are reported with 1–2 biological replicates ± standard deviation (**E**, **F**).

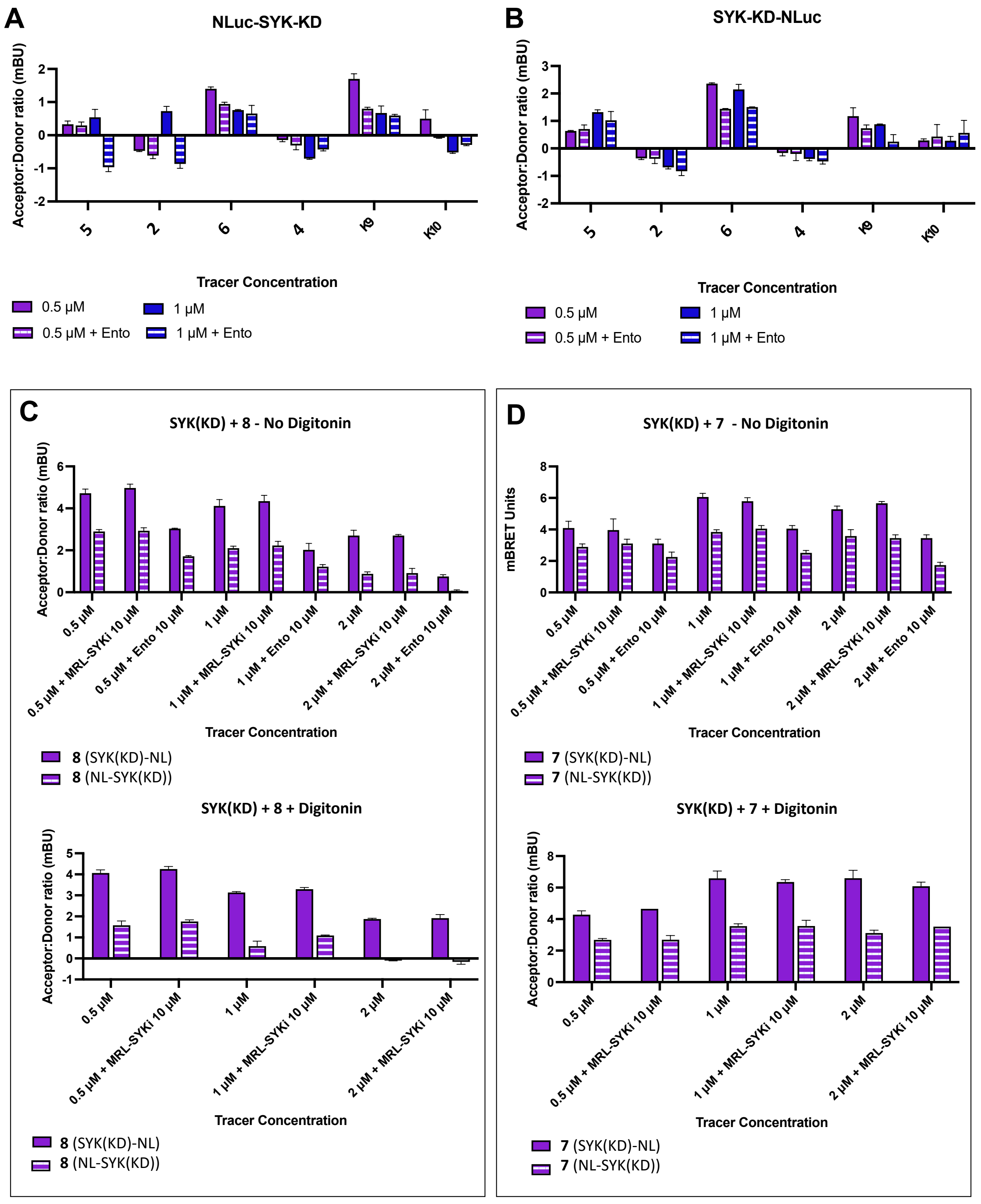


**Supplementary Figure 2.** SYK NanoBRET tracers **2**, **4**, **5**, **6**, **7**, **8, K9**, and **K10** did not generate a BRET signal (mBU) with SYK(KD)-NL or NL-SYK(KD) ± digitonin (**A**–**D**). Data are reported as technical duplicates ± standard deviation (**A**–**D**).

A

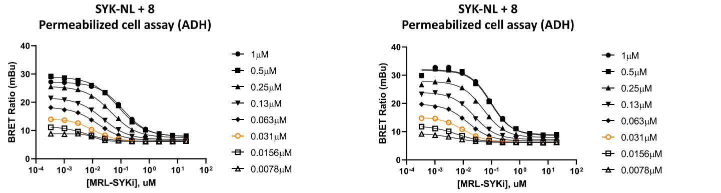


| 1μM | 0.5μM | 0.25μM | 0.13μM | 0.063μM | 0.031μM | 0.0156μM | 0.0078μM |
|---|---|---|---|---|---|---|---|
| 0.1160 | 0.08067 | 0.04454 | 0.02297 | 0.01500 | 0.01014 | 0.006042 | 0.01218 |

| 1μM | 0.5μM | 0.25μM | 0.13μM | 0.063μM | 0.031μM | 0.0156μM | 0.0078μM |
|---|---|---|---|---|---|---|---|
| 0.09703 | 0.09084 | 0.05142 | 0.02412 | 0.01449 | 0.009823 | 0.005907 | 0.004837 |

B

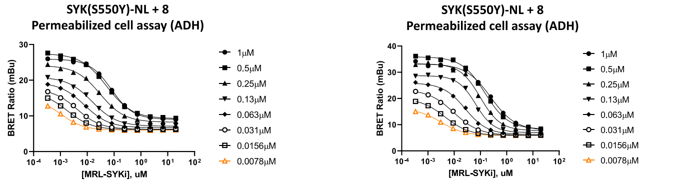


| 1μM | 0.5μM | 0.25μM | 0.13μM | 0.063μM | 0.031μM | 0.0156μM | 0.0078μM |
|---|---|---|---|---|---|---|---|
| 0.07911 | 0.05251 | 0.02999 | 0.01529 | 0.007911 | 0.004136 | 0.001850 | 0.001100 |

| 1μM | 0.5μM | 0.25μM | 0.13μM | 0.063μM | 0.031μM | 0.0156μM | 0.0078μM |
|---|---|---|---|---|---|---|---|
| 0.2462 | 0.1495 | 0.09825 | 0.06566 | 0.02624 | 0.009585 | 0.004440 | 0.003152 |

C

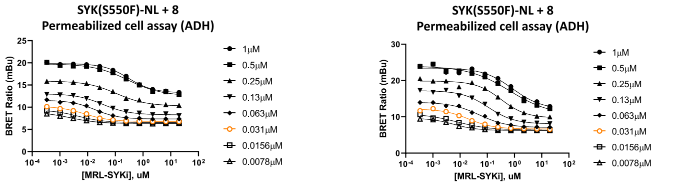


| 1μM | 0.5μM | 0.25μM | 0.13μM | 0.063μM | 0.031μM | 0.0156μM | 0.0078μM |
|---|---|---|---|---|---|---|---|
| 0.2991 | 0.2809 | 0.1102 | 0.03496 | 0.01849 | 0.007820 | 0.004570 | 0.002387 |

| 1μM | 0.5μM | 0.25μM | 0.13μM | 0.063μM | 0.031μM | 0.0156μM | 0.0078μM |
|---|---|---|---|---|---|---|---|
| 0.8527 | 0.6568 | 0.3262 | 0.1093 | 0.05096 | 0.02068 | 0.007451 | 0.005868 |

D

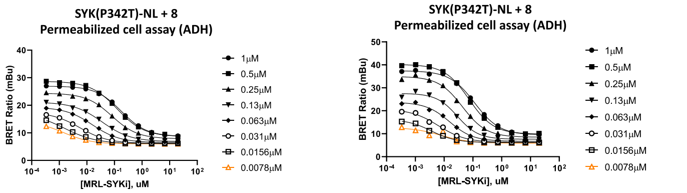


| 1μM | 0.5μM | 0.25μM | 0.13μM | 0.063μM | 0.031μM | 0.0156μM | 0.0078μM |
|---|---|---|---|---|---|---|---|
| 0.1783 | 0.1261 | 0.08052 | 0.03527 | 0.01596 | 0.005601 | 0.001563 | 0.001436 |

| 1μM | 0.5μM | 0.25μM | 0.13μM | 0.063μM | 0.031μM | 0.0156μM | 0.0078μM |
|---|---|---|---|---|---|---|---|
| 0.1149 | 0.07369 | 0.04362 | 0.02902 | 0.01546 | 0.006368 | 0.003587 | 0.007826 |

E

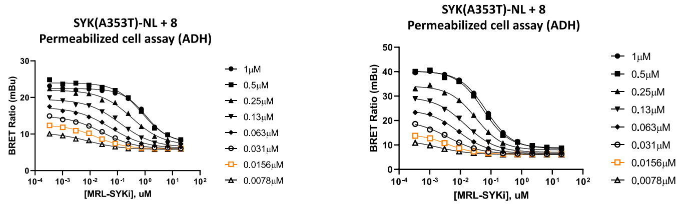


| 1μM | 0.5μM | 0.25μM | 0.13μM | 0.063μM | 0.031μM | 0.0156μM | 0.0078μM |
|---|---|---|---|---|---|---|---|
| 1.059 | 0.7680 | 0.3282 | 0.1236 | 0.05597 | 0.02784 | 0.01689 | 0.008468 |

| 1μM | 0.5μM | 0.25μM | 0.13μM | 0.063μM | 0.031μM | 0.0156μM | 0.0078μM |
|---|---|---|---|---|---|---|---|
| 0.06818 | 0.05489 | 0.03206 | 0.01450 | 0.008647 | 0.004024 | 0.003702 | 0.001751 |

F

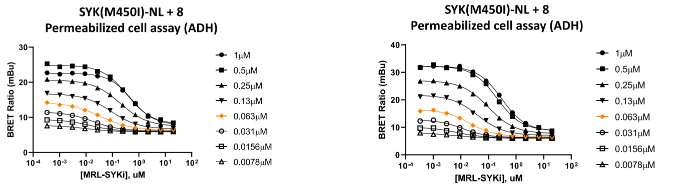


| 1μM | 0.5μM | 0.25μM | 0.13μM | 0.063μM | 0.031μM | 0.0156μM | 0.0078μM |
|---|---|---|---|---|---|---|---|
| 0.5212 | 0.3575 | 0.1953 | 0.08893 | 0.03536 | 0.01871 | 0.01463 | 0.01022 |

| 1μM | 0.5μM | 0.25μM | 0.13μM | 0.063μM | 0.031μM | 0.0156μM | 0.0078μM |
|---|---|---|---|---|---|---|---|
| 0.3014 | 0.2071 | 0.09867 | 0.04076 | 0.02122 | 0.01191 | 0.007194 | 0.004140 |

**Supplementary Figure 3.** SYK NanoBRET tracer **8** titrations in cells permeabilized with digitonin. An 11-point dose–response study of MRL-SYKi from a top concentration of 20 μM was used to compete away the BRET signal. Two biological replicates (left and right) and the results plotted using a log(inhibitor) vs. response (three parameter) fit. (**A**) SYK(WT)-NL (**B**) SYK(S550Y)-NL (**C**) SYK(S550F)-NL (**D**) SYK(P342T)-NL (**E**) SYK(A353T)-NL (**F**) SYK(M450I)-NL.

**A**

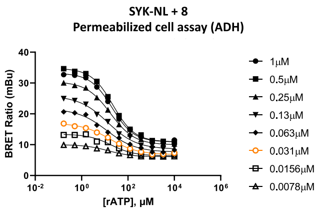


| 1μM | 0.5μM | 0.25μM | 0.13μM | 0.063μM | 0.031μM | 0.0156μM | 0.0078μM |
|---|---|---|---|---|---|---|---|
| 20.52 | 22.55 | 17.39 | 13.52 | 12.08 | 16.26 | 19.15 | 13.95 |

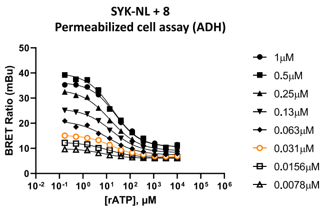


| 1μM | 0.5μM | 0.25μM | 0.13μM | 0.063μM | 0.031μM | 0.0156μM | 0.0078μM |
|---|---|---|---|---|---|---|---|
| 23.96 | 18.45 | 14.52 | 16.09 | 11.41 | 17.31 | 11.77 | 6.581 |

**B**

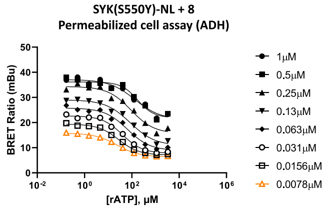


| 1μM | 0.5μM | 0.25μM | 0.13μM | 0.063μM | 0.031μM | 0.0156μM | 0.0078μM |
|---|---|---|---|---|---|---|---|
| 135.2 | 173.9 | 92.01 | 85.63 | 64.83 | 39.78 | 27.60 | 23.28 |

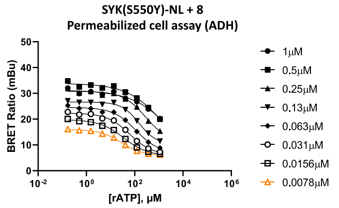


| 1μM | 0.5μM | 0.25μM | 0.13μM | 0.063μM | 0.031μM | 0.0156μM | 0.0078μM |
|---|---|---|---|---|---|---|---|
| 14302 | 1686 | 280.6 | 152.2 | 111.2 | 61.27 | 34.14 | 27.95 |

**C**

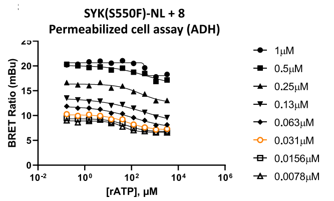


| 1μM | 0.5μM | 0.25μM | 0.13μM | 0.063μM | 0.031μM | 0.0156μM | 0.0078μM |
|---|---|---|---|---|---|---|---|
| 382.9 | 427.9 | 238.3 | 386.2 | 111.2 | 68.52 | 59.77 | 42.12 |

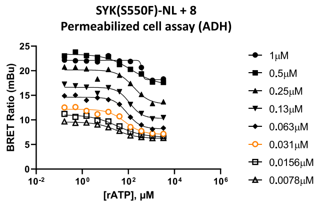


| 1μM | 0.5μM | 0.25μM | 0.13μM | 0.063μM | 0.031μM | 0.0156μM | 0.0078μM |
|---|---|---|---|---|---|---|---|
| 389.1 | 135.5 | 168.6 | 114.1 | 89.22 | 71.03 | 33.83 | 30.06 |

**D**

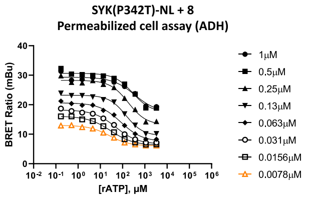


| 1μM | 0.5μM | 0.25μM | 0.13μM | 0.063μM | 0.031μM | 0.0156μM | 0.0078μM |
|---|---|---|---|---|---|---|---|
| 352.1 | 282.0 | 180.2 | 143.1 | 84.70 | 48.34 | 29.34 | 25.24 |

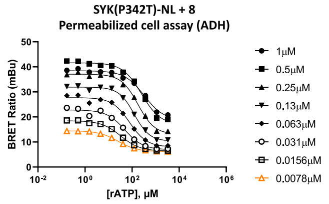


| 1μM | 0.5μM | 0.25μM | 0.13μM | 0.063μM | 0.031μM | 0.0156μM | 0.0078μM |
|---|---|---|---|---|---|---|---|
| 329.8 | 250.7 | 177.1 | 123.4 | 77.58 | 49.06 | 32.80 | 25.31 |

**E**

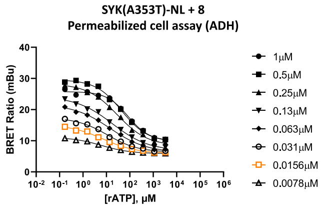


| 1μM | 0.5μM | 0.25μM | 0.13μM | 0.063μM | 0.031μM | 0.0156μM | 0.0078μM |
|---|---|---|---|---|---|---|---|
| 84.28 | 45.43 | 29.35 | 18.12 | 14.56 | 11.10 | 7.210 | 6.099 |

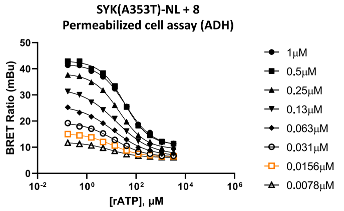


| 1μM | 0.5μM | 0.25μM | 0.13μM | 0.063μM | 0.031μM | 0.0156μM | 0.0078μM |
|---|---|---|---|---|---|---|---|
| 37.55 | 33.70 | 20.46 | 15.50 | 12.16 | 12.96 | 10.97 | 9.064 |

**F**

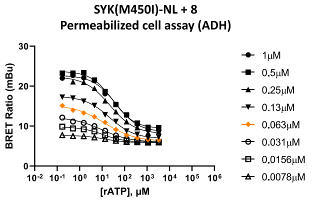


| 1μM | 0.5μM | 0.25μM | 0.13μM | 0.063μM | 0.031μM | 0.0156μM | 0.0078μM |
|---|---|---|---|---|---|---|---|
| 40.84 | 29.61 | 22.12 | 21.43 | 13.24 | 9.791 | 11.05 | 13.08 |

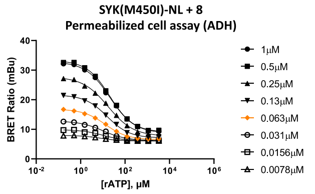


| 1μM | 0.5μM | 0.25μM | 0.13μM | 0.063μM | 0.031μM | 0.0156μM | 0.0078μM |
|---|---|---|---|---|---|---|---|
| 20.80 | 20.75 | 17.23 | 15.50 | 12.92 | 10.23 | 10.06 | 7.824 |

**Supplementary Figure 4.** SYK NanoBRET tracer **8** titrations in cells permeabilized with digitonin. An 11-point dose–response study of ATP from a top concentration of 10 mM was used to compete away the BRET signal. Two biological replicates (left and right) and the results plotted in GraphPad Prism using a log(inhibitor) vs. response (three parameter) fit. (**A**) SYK(WT)-NL (**B**) SYK(S550Y)-NL (**C**) SYK(S550F)-NL (**D**) SYK(P342T)-NL (**E**) SYK(A353T)-NL (**F**) SYK(M450I)-NL.

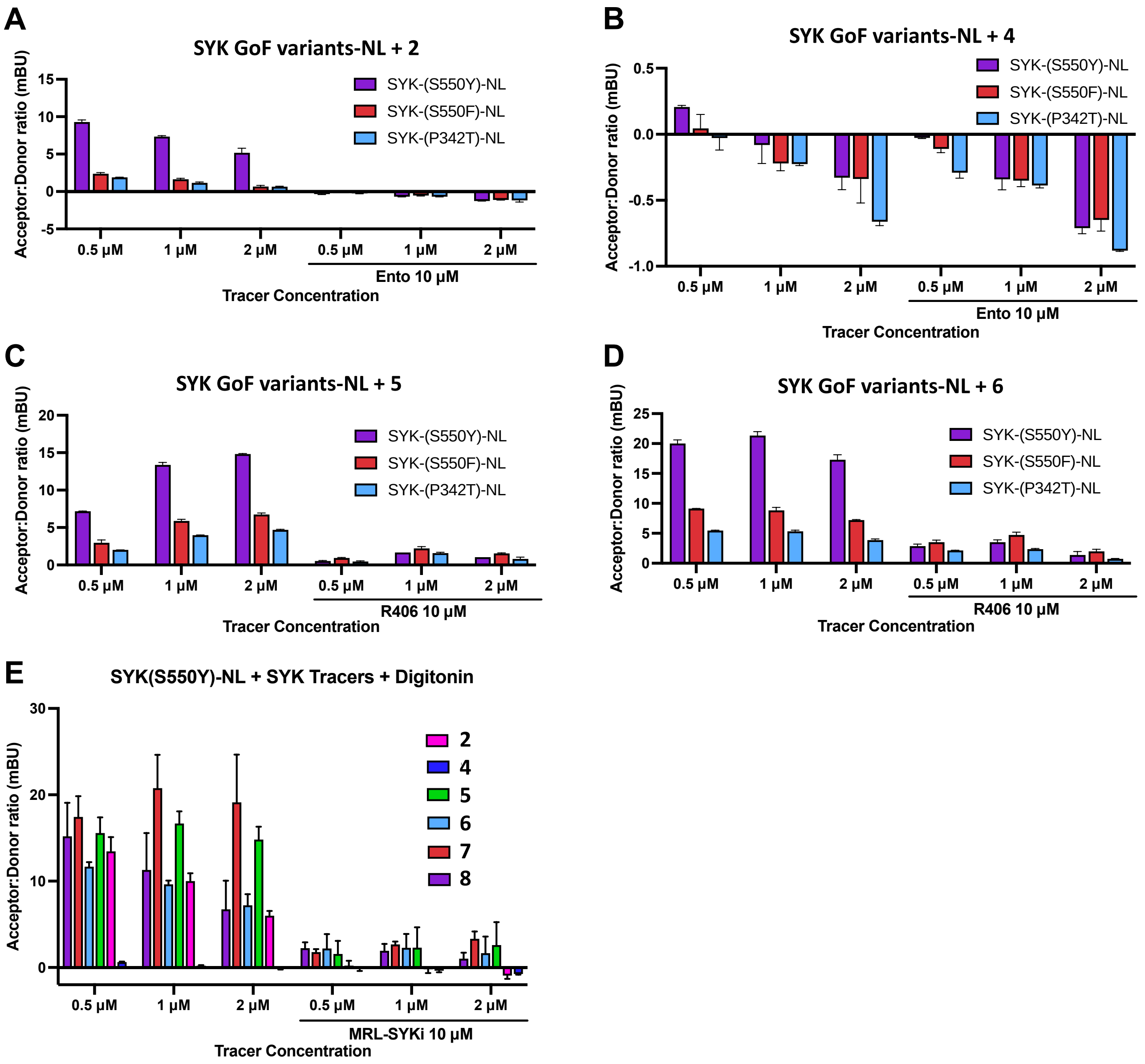


**Supplementary Figure 5.** SYK NanoBRET tracers **2** (**A**), **5** (**C**), **6** (**D**) generated BRET signal (mBU) with SYK GoF variants in the NanoBRET assay that can be competed away with SYK inhibitors. BRET signal is greater with SYK(S550Y) than SYK(S550F) and least with SYK(P342T). (**B**) Tracer **4** does not generate a BRET signal with any GoF variants, which correlates with its inability to bind to SYK in enzymatic inhibition assays. (**E**) Tracers **2**, **5**, **6**, **7**, and **8** all generate a BRET signal with SYK(S550Y) + digitonin. The mBU are decreased compared to intact cells. Data are reported in biological duplicate ± standard deviation (**A**–**D**). Data are reported in biological duplicates each with two technical replicates ± standard deviation (**E**).

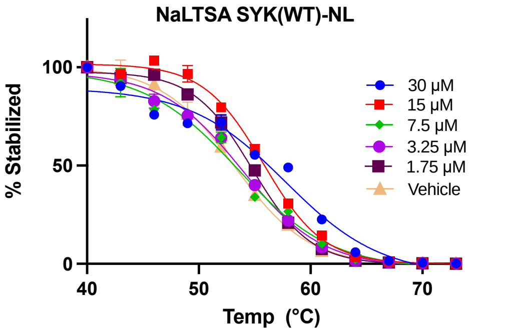


| | 30 μM | 15 μM | 7.5 μM | 3.25 μM | 1.75 μM | Vehicle |
|---|---|---|---|---|---|---|
| Melting Temp ° C | 57.8 | 55.8 | 53.5 | 53.9 | 54.7 | 53.2 |

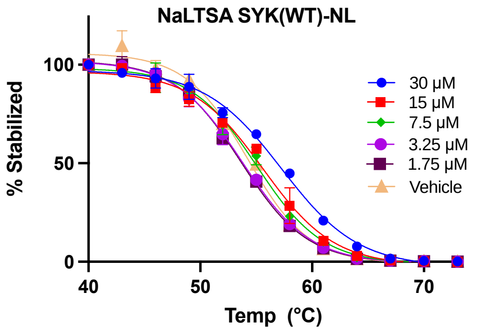


| | 30 μM | 15 μM | 7.5 μM | 3.25 μM | 1.75 μM | Vehicle |
|---|---|---|---|---|---|---|
| Melting Temp ° C | 57.3 | 55.6 | 55.1 | 53.8 | 53.6 | 54.2 |

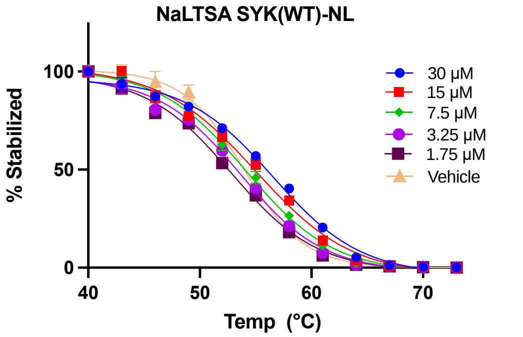


| | 30 μM | 15 μM | 7.5 μM | 3.25 μM | 1.75 μM | Vehicle |
|---|---|---|---|---|---|---|
| Melting Temp ° C | 56.6 | 55.0 | 54.1 | 53.7 | 52.8 | 53.9 |

**Supplementary Figure 6.** Three biological replicates for the NaLTSA assay with SYK(WT)-NL, evaluating MRL-SYKi in dose–response format, corresponding to Figure 5 in the main text. The data was fitted with a Boltzmann Sigmoid equation and error bars indicate the standard deviation.

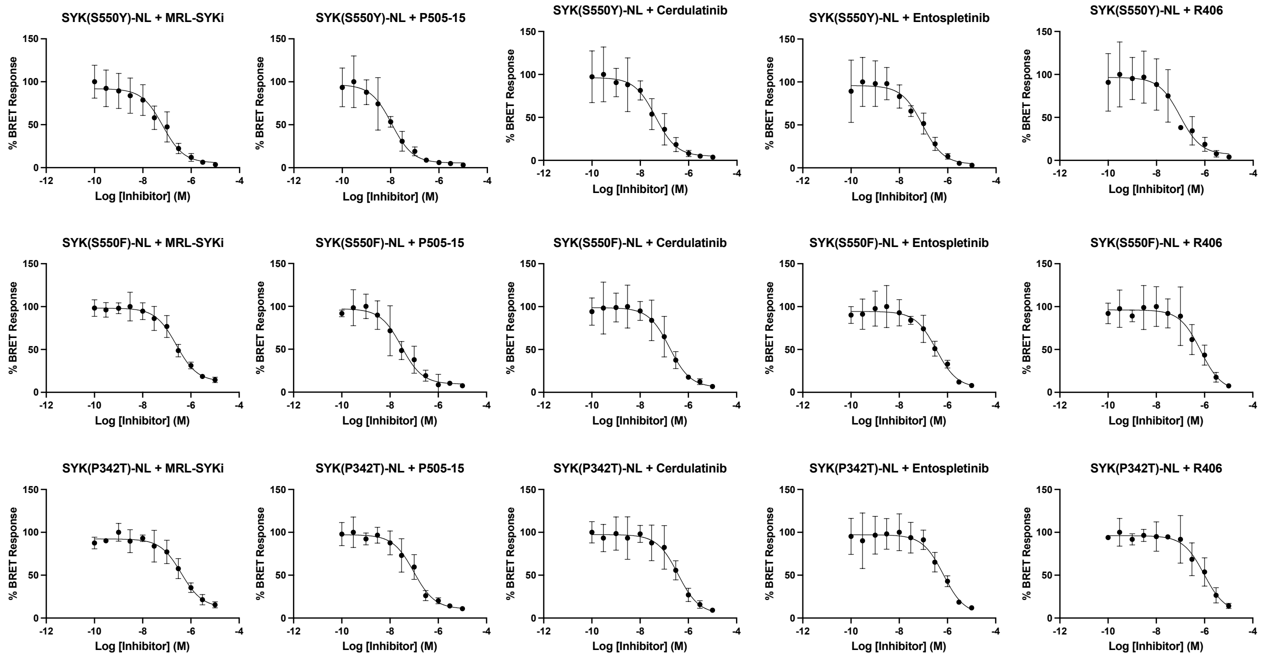


**Supplementary Figure 7.** SYK inhibitor dose–response curves with SYK(GoF)-NL variants. Data are reported as three biological replicates ± standard deviation and plotted in GraphPad Prism using a log(inhibitor) vs. response (three parameter) fit.

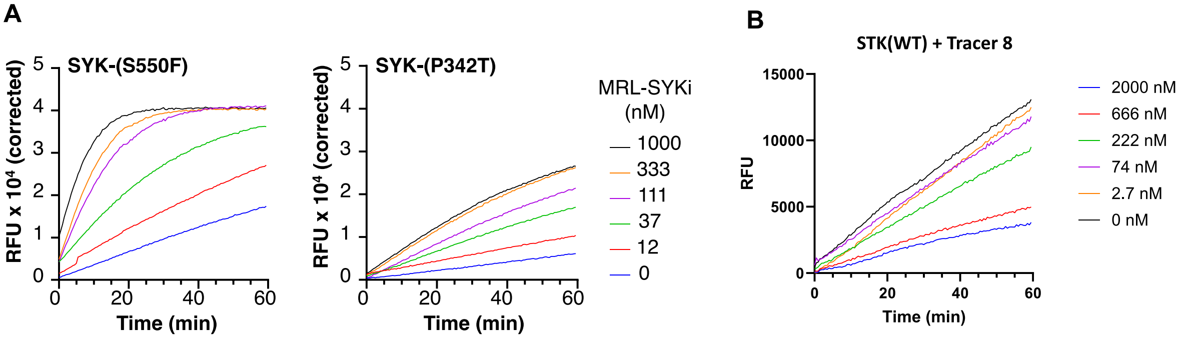


**Supplementary Figure 8.** SYK PhosphoSens Assay with SYK SOX-peptide substrate. (**A**) Raw data for the dose–response curves of MRL-SYKi with SYK(S550F) and SYK(P342T) in Figure 5D within the main text. (**B**) Dose–response evaluation of tracer **8** with SYK(WT) indicating inhibition of SYK catalytic activity.

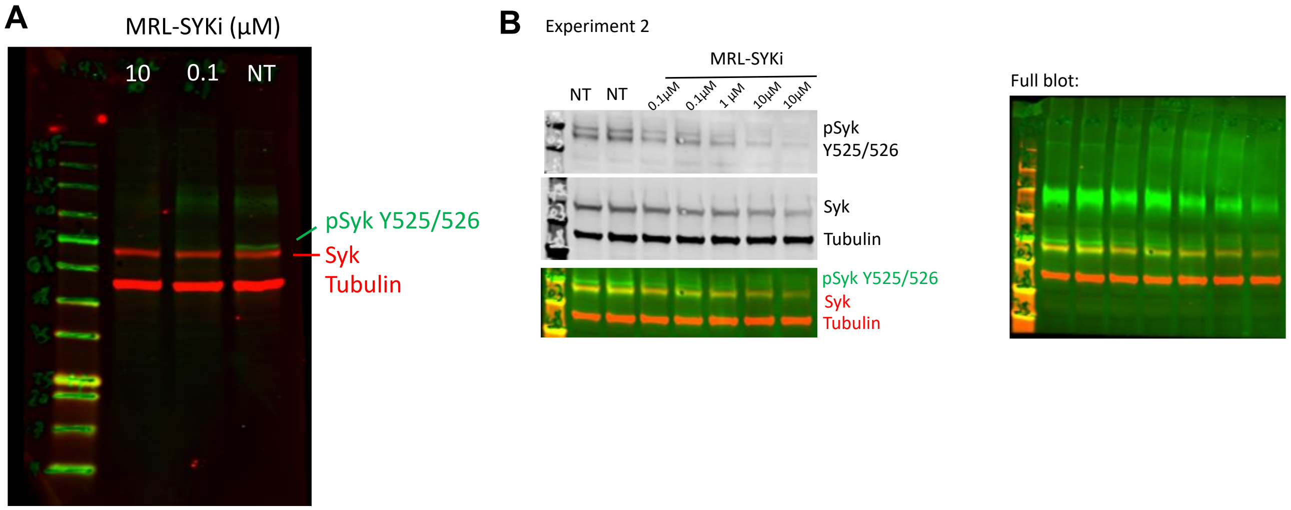


**Supplementary Figure 9.** Phosphorylation levels at SYK(Y525/526) decrease upon MRL-SYKi inhibitor treatment after 20 hrs at 100 nM and 10 µM in a stably expressing SYK(S550Y) SW480 cell line. (**A**) Full western blot corresponding to the western blot included in Figure 5E in the main text (biological replicate number 1). (**B**) Biological replicate number 2 of MRL-SYKi inhibitor treatment.

#### 2 Chemistry Experimental

##### 2.1 General Information for Chemical Synthesis

Reagents were obtained from verified suppliers and employed without purification. Temperatures are in degrees Celsius (°C); solvent was removed using a rotary evaporator; thin layer chromatography and LC–MS were used to monitor reaction progress. $^{1}$H NMR and additional analytical data was collected for intermediates and final compounds to verify their identity and evaluate their purity. $^{1}$H and $^{13}$C NMR spectra were obtained in DMSO-$d_6$, MeOD-$d_4$, or $CDCl_3$ and recorded using Bruker spectrometers. Magnet strength is listed in each experimental write-up along with peak positions in parts per million (ppm). Peaks in NMR spectra are calibrated versus the shift of the indicated deuterated solvent and coupling constants ($J$ values) are reported in hertz (Hz). Peak multiplicities are included as follows: singlet (s), doublet (d), doublet of doublets (dd), triplet (t), pentet (p), and multiplet (m). The purity of all final compounds was assessed via HPLC.

##### 2.2 Abbreviations

DIPEA (*N*-ethyl-*N*-isopropylpropan-2-amine); DMF (*N*,*N*-dimethylformamide), MeOH (methanol), EtOH (ethanol), Ar (argon), TBTU (2-(1*H*-benzo[*d*][1,2,3]triazol-1-yl)-1,1,3,3-tetramethyluronium tetrafluoroborate), rt (room temperature), $H_2O$ (water), TFA (trifluoroacetic acid), THF (tetrahydrofuran).

##### 2.3 Chemistry Schemes

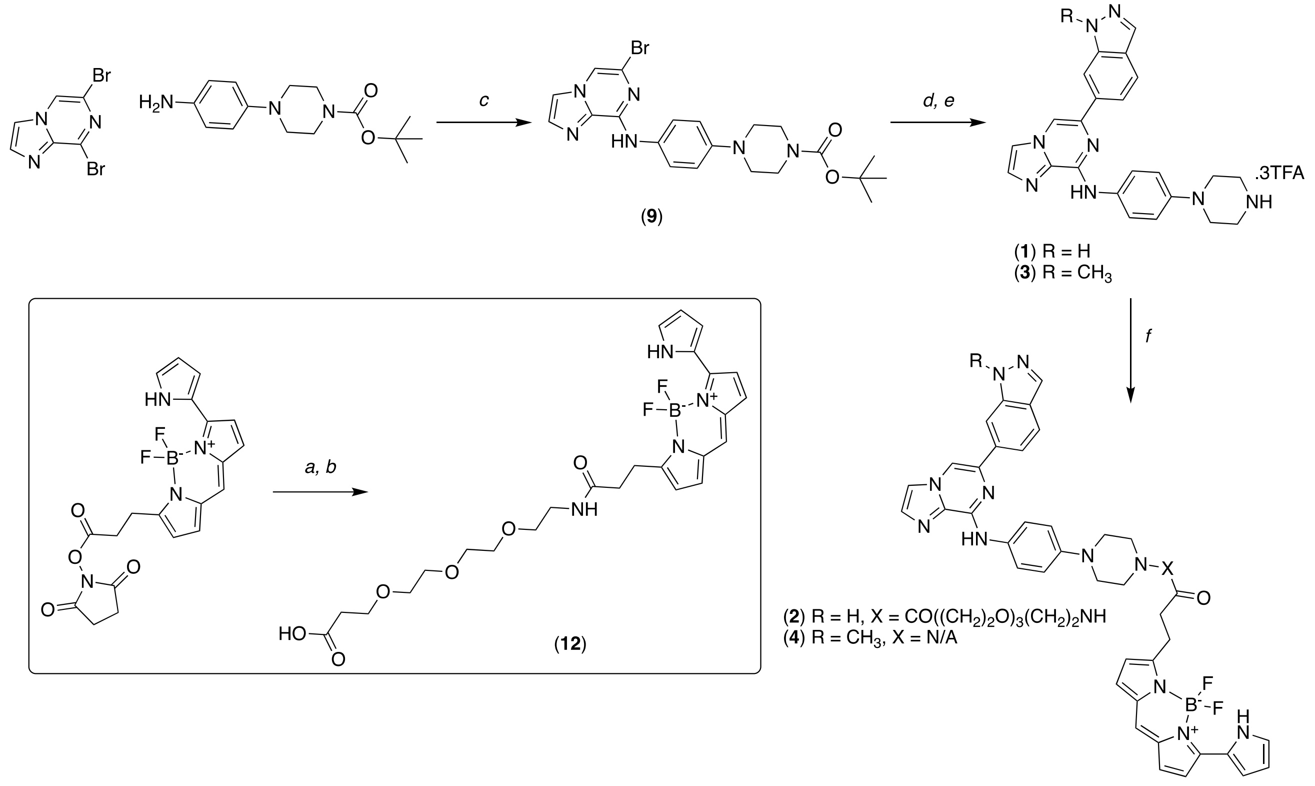


**Scheme 1.** Synthesis of NanoBRET tracers **2** and **4**. Reagents and conditions: a) *tert*-butyl 3-(2-(2-(2-aminoethoxy)ethoxy)ethoxy)propanoate, 2,5-dioxopyrrolidin-1-yl 3-(5,5-difluoro-7-(1*H*-pyrrol-2-yl)-5*H*-5$\lambda^4$,6$\lambda^4$-dipyrrolo[1,2-c:2',1'-*f*][1,3,2]diazaborinin-3-yl)propanoate, DIPEA, DMF. b) TFA, $CH_2Cl_2$, rt, 16 hrs. c) (**9**): 6,8-dibromoimidazo[1,2-*a*]pyrazine, *tert*-butyl 4-(4-aminophenyl)piperazine-1-carboxylate, DIPEA, propan-2-ol, 85 °C, 16 hrs. d) (1*H*-indazol-6-yl)boronic acid or (1-methyl-1*H*-indazol-6-yl)boronic acid , **9**, $Na_2CO_3$, $PdCl_2$(dppf), $H_2O$, 1,4-dioxane (0.25 M), 100 °C, $N_2$, 16 hrs. e) TFA, $CH_2Cl_2$, rt, 16 hrs. f) (**2**): **12**, **1**, DIPEA, TBTU, DMF, rt, 16 hrs. (**4**): 2,5-dioxopyrrolidin-1-yl 3-(5,5-difluoro-7-(1*H*-pyrrol-2-yl)-5*H*-5$\lambda^4$,6$\lambda^4$-dipyrrolo[1,2-*c*:2',1'-*f*][1,3,2]diazaborinin-3-yl)propanoate, **3**, DIPEA, DMF, rt, 10 mins.

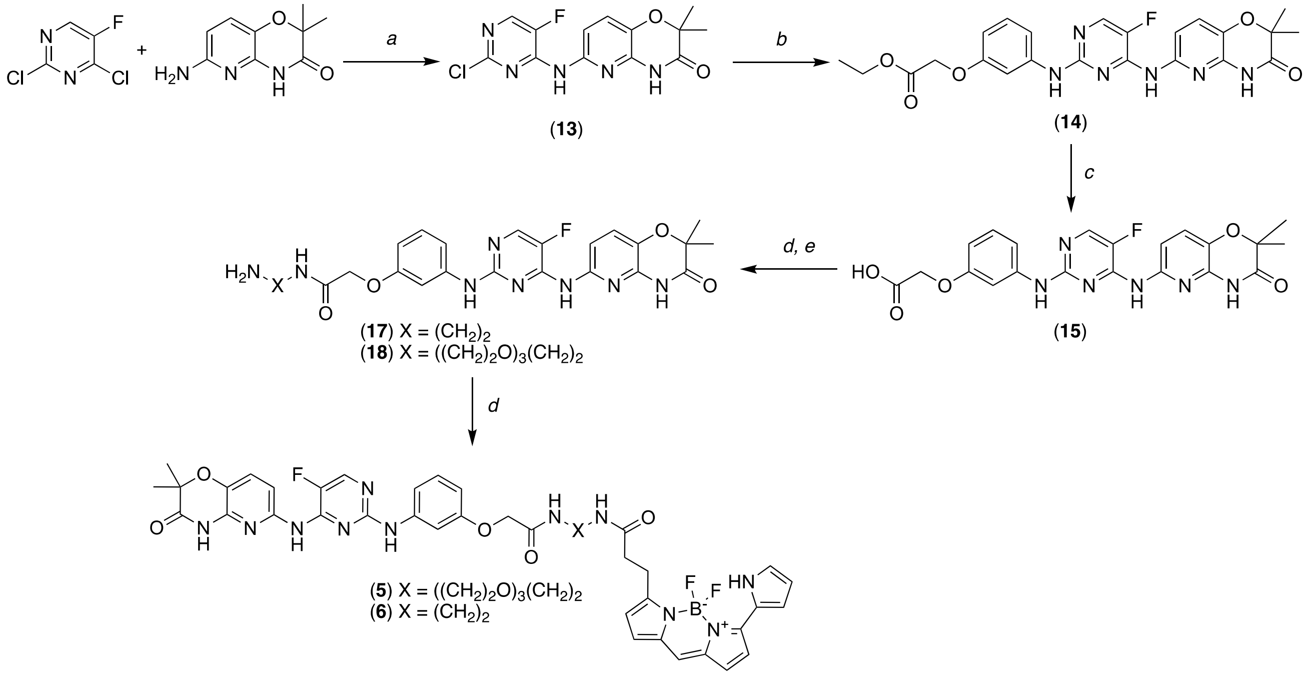


**Scheme 2.** Synthesis of NanoBRET tracers **5** and **6**. Reagents and conditions: a) 2,4-dichloro-5-fluoropyrimidine, 6-amino-2,2-dimethyl-2*H*-pyrido[3,2-*b*][1,4]oxazin-3(4*H*)-one, MeOH, $H_2O$, 80 °C, 16 hrs. b) ethyl 2-(3-aminophenoxy)acetate, **13**, EtOH, Ar, 80 °C, 16 hrs. c) **14**, LiOH, THF, $H_2O$, 50 ºC, 16 hrs. d) (**17**): **15**, DIPEA, TBTU, *tert*-butyl (2-aminoethyl)carbamate, DMF, 72 hrs. (**18**): **15**, DIPEA, TBTU, *tert*-butyl (2-(2-(2-(2-aminoethoxy)ethoxy)ethoxy)ethyl)carbamate, DMF, 16 hrs. e) TFA, $CH_2Cl_2$, rt, 16 hrs. d) **17** or **18**, 2,5-dioxopyrrolidin-1-yl 3-(5,5-difluoro-7-(1*H*-pyrrol-2-yl)-5*H*-5$\lambda^4$,6$\lambda^4$-dipyrrolo[1,2-*c*:2',1'-*f*][1,3,2]diazaborinin-3-yl)propanoate, DIPEA, DMF, rt, 30 mins.

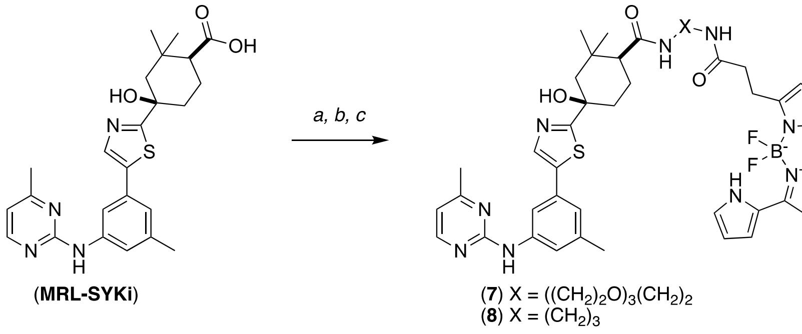


**Scheme 3.** Synthesis of NanoBRET tracers **7** and **8**. Reagents and conditions: a) (**7**): MRL-SYKi, *tert*-butyl (2-(2-(2-(2-aminoethoxy)ethoxy)ethoxy)ethyl)carbamate, DIPEA, HATU, DMF, rt,16 hrs. b) TFA, $CH_2Cl_2$, rt, 1 hr. c) (1*S*,4*R*)-*N*-(2-(2-(2-(2-aminoethoxy)ethoxy)ethoxy)ethyl)-4-hydroxy-2,2-dimethyl-4-(5-(3-methyl-5-((4-methylpyrimidin-2-yl)amino)phenyl)thiazol-2-yl)cyclohexane-1-carboxamide 2,2,2-trifluoroacetate, 2,5-dioxopyrrolidin-1-yl 3-(5,5-difluoro-7-(1*H*-pyrrol-2-yl)-5*H*-5λ4,6λ4-dipyrrolo[1,2-*c*:2',1'-*f*][1,3,2]diazaborinin-3-yl)propanoate, DIPEA, DMF, rt, 30 mins. (**8**): a) MRL-SYKi *tert*-butyl (3-aminopropyl)carbamate, DIPEA, HATU, DMF, rt,16 hrs. b) TFA, $CH_2Cl_2$, rt, 1 hr. c) (1*S*,4*R*)-*N*-(3-aminopropyl)-4-hydroxy-2,2-dimethyl-4-(5-(3-methyl-5-((4-methylpyrimidin-2-yl)amino)phenyl)thiazol-2-yl)cyclohexane-1-carboxamide 2,2,2-trifluoroacetate, 2,5-dioxopyrrolidin-1-yl 3-(5,5-difluoro-7-(1*H*-pyrrol-2-yl)-5*H*-5λ4,6λ4-dipyrrolo[1,2-*c*:2',1'-*f*][1,3,2]diazaborinin-3-yl)propanoate, DIPEA, DMF, rt, 30 mins.

#### 2.4 Chemistry Experimental Data

***Tert*-butyl 4-(4-((6-bromoimidazo[1,2-*a*]pyrazin-8-yl)amino)phenyl)piperazine-1-carboxylate (9)**

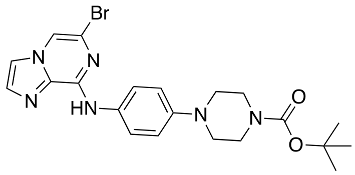


To a flask was added 6,8-dibromoimidazo[1,2-*a*]pyrazine (250 mg, 1.00 equiv., 903 μmol), *tert*-butyl 4-(4-aminophenyl)piperazine-1-carboxylate (263 mg, 1.05 equiv., 948 μmol), and DIPEA (236 μL, 1.50 equiv., 1.35 mmol) in propan-2-ol (3.61 mL, 0.25 molar, 1.00 equiv., 903 μmol) and the reaction was heated to 85 °C for 16 hrs. The reaction was concentrated *in vacuo* and purified via column chromatography (silica, 0-100% EtOAc in hexane) to yield the desired product as a yellow solid (218 mg, 51%). $^{1}H$ NMR (400 MHz, $CDCl_3$) δ 9.25 (s, 1H), 7.86 (d, $J$ = 8.9 Hz, 2H), 7.74 (s, 1H), 7.66 – 7.56 (m, 2H), 7.14 (s, 2H), 3.70 (s, 4H), 3.18 (s, 4H), 1.48 (s, 9H). LCMS calculated for $C_{21}H_{26}BrN_6O_2$ $[M + H]^+$: 473.12. Found: 473.08.

***Tert*-butyl 4-(4-((6-(1*H*-indazol-6-yl)imidazo[1,2-*a*]pyrazin-8-yl)amino)phenyl)piperazine-1-carboxylate (10)**

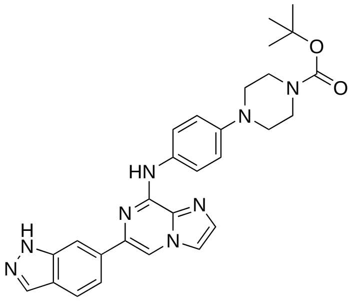


To a microwave vial was added (1*H*-indazol-6-yl)boronic acid (112 mg, 1.5 equiv., 690 μmol), *tert*-butyl 4-(4-((6-bromoimidazo[1,2-*a*]pyrazin-8-yl)amino)phenyl)piperazine-1-carboxylate (218 mg, 1.0 equiv., 460 μmol), $Na_2CO_3$ (73 mg, 1.5 equiv., 690 μmol), and $PdCl_2$(dppf) (34 mg, 0.1 equiv., 46 μmol) in $H_2O$ (460 μL) and 1,4-dioxane (1.84 mL, 0.25 M) and the reaction was heated to 100 °C under $N_2$ for 16 hrs. The reaction was cooled to rt, filtered through celite, and concentrated *in vacuo*. The crude material was purified by column chromatography (silica, 0-10% MeOH in $CH_2Cl_2$) to yield the desired product as a yellow solid (141 mg, 60%). $^{1}H$ NMR (400 MHz, DMSO) δ 13.18 (s, 1H), 9.54 (s, 1H), 8.67 (s, 1H), 8.18 (s, 1H), 8.09 (s, 1H), 8.03 (d, $J$ = 9.1 Hz, 2H), 7.99 (d, $J$ = 1.0 Hz, 1H), 7.84 (d, $J$ = 8.6 Hz, 1H), 7.72 (dd, $J$ = 8.6, 1.3 Hz, 1H), 7.64 (d, $J$ = 1.0 Hz, 1H), 7.02 (d, $J$ = 9.1 Hz, 2H), 3.53 – 3.45 (m, 4H), 3.12 – 3.05 (m, 4H), 1.43 (s, 9H). $^{13}C$ NMR (214 MHz, DMSO) δ 153.91, 146.46, 144.83, 140.51, 136.46, 135.18, 133.47, 132.79, 132.31, 132.14, 122.58, 121.15, 120.67, 118.31, 116.47, 108.72, 107.12, 79.00, 49.18, 28.09.

**6-(1-methyl-1*H*-indazol-6-yl)-*N*-(4-(piperazin-1-yl)phenyl)imidazo[1,2-*a*]pyrazin-8-amine (3)**

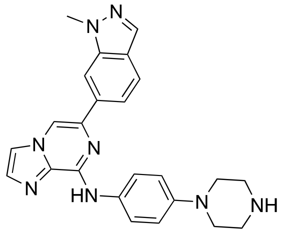


To a microwave vial was added *tert*-butyl 4-(4-((6-bromoimidazo[1,2-*a*]pyrazin-8-yl)amino)phenyl)piperazine-1-carboxylate (50 mg, 1.0 equiv., 0.11 mmol), (1-methyl-1*H*-indazol-6-yl)boronic acid (37 mg, 2.0 equiv., 0.21 mmol), $Na_2CO_3$ (25 mg, 2.2 equiv., 0.23 mmol), 1,4-dioxane (0.74 mL) and $H_2O$ (0.3 mL). The solution was degassed with nitrogen for 5 mins before $Pd(PPh_3)_4$ (12 mg, 0.1 equiv., 11 μmol) was added. The reaction was heated at 100˚C for 19 hrs. The reaction was cooled, dissolved in EtOAc, and filtered *in vacuo* through celite. The crude material was purified by column chromatography (silica, 0-100% EtOAc in hexane) to yield *tert*-butyl 4-(4-((6-(1-methyl-1*H*-indazol-6-yl)imidazo[1,2-*a*]pyrazin-8-yl)amino)phenyl)piperazine-1-carboxylate, which was used in the next step. To a flask was added *tert*-butyl 4-(4-((6-(1-methyl-1*H*-indazol-6-yl)imidazo[1,2-*a*]pyrazin-8-yl)amino)phenyl)piperazine-1-carboxylate (29 mg, 1.0 equiv., 55 μmol) with 20% TFA in $CH_2Cl_2$. The reaction was stirred at rt for 36 hrs, concentrated *in vacuo*, and purified by preparative-HPLC (10-100% MeOH in $H_2O$ (+ 0.01% TFA) to yield the desired product as a white solid. $^1$H NMR (400 MHz, MeOD-$d_4$) δ 8.52 (d, $J$ = 1.6 Hz, 1H), 8.11 (s, 1H), 8.04 – 7.96 (m, 2H), 7.91 – 7.87 (m, 2H), 7.83 (d, $J$ = 1.5 Hz, 1H), 7.78 – 7.75 (m, 1H), 7.73 – 7.70 (m, 1H), 7.10 – 7.02 (m, 2H), 4.06 (d, $J$ = 1.3 Hz, 3H), 3.39 (s, 8H). $^{13}$C NMR (214 MHz, MeOD-$d_4$) δ 147.69, 144.85, 141.73, 140.52, 136.74, 134.85, 133.72, 130.62, 125.27, 122.29, 122.22, 118.74, 117.96, 116.72, 115.39, 109.94, 108.00, 54.78, 48.46, 44.93, 35.58. LCMS calculated for $C_{24}H_{25}N_8$ $[M + H]^+$: 425.21. Found: 425.53.

**3-(5,5-difluoro-7-(1*H*-pyrrol-2-yl)-5*H*-5λ⁴,6λ⁴-dipyrrolo[1,2-*c*:2',1'-*f*][1,3,2]diazaborinin-3-yl)-1-(4-(4-((6-(1-methyl-1*H*-indazol-6-yl)imidazo[1,2-*a*]pyrazin-8-yl)amino)phenyl)piperazin-1-yl)propan-1-one (4)**

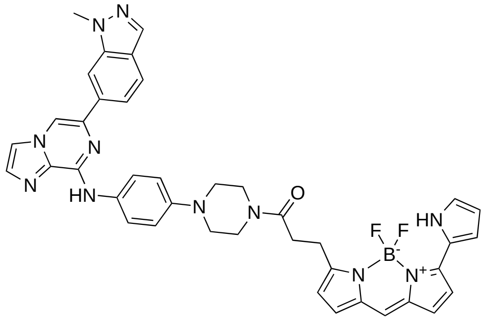


To a flask was added 2,5-dioxopyrrolidin-1-yl 3-(5,5-difluoro-7-(1*H*-pyrrol-2-yl)-5*H*-5$\lambda^4$,6$\lambda^4$-dipyrrolo[1,2-*c*:2',1'-*f*][1,3,2]diazaborinin-3-yl)propanoate (5 mg, 1.0 equiv., 0.01 mmol), 6-(1-methyl-1*H*-indazol-6-yl)-*N*-(4-(piperazin-1-yl)phenyl)imidazo[1,2-*a*]pyrazin-8-amine (5 mg, 1.0 equiv, 0.01 mmol), and DIPEA (10 μL, 6.0 equiv., 0.06 mmol) in DMF (100 μL, 0.1M). The reaction was stirred at rt for 10 mins. The reaction was concentrated *in vacuo* and purified by preparative-HPLC (10-100% MeOH in $H_2O$ (+ 0.01% TFA) to yield the desired product as a purple solid (0.9 mg, 10%). $^1$H NMR (400 MHz, $CD_3OD$) δ 8.55 (s, 1H), 8.19 (s, 1H), 8.02 (s, 1H), 8.00 – 7.93 (m, 3H), 7.84 – 7.76 (m, 2H), 7.75 (s, 1H), 7.23 (s, 1H), 7.21 – 7.16 (m, 3H), 7.14 (d, $J$ = 8.9 Hz, 2H), 6.99 (d, $J$ = 4.6 Hz, 1H),

6.92 (d, *J* = 4.0 Hz, 1H), 6.36 (d, *J* = 3.9 Hz, 1H), 6.34 – 6.31 (m, 1H), 4.09 (s, 3H), 3.85 – 3.80 (m, 2H), 3.77 – 3.72 (m, 2H), 3.24 – 3.20 (m, 2H), 3.19 – 3.25 (m, 2H), 2.95 – 2.84 (m, 2H). LCMS calculated for $C_{40}H_{37}BF_2N_{11}O$ $[M + H]^+$: 736.32. Found: 736.0.

***Tert*-butyl 1-(5,5-difluoro-7-(1*H*-pyrrol-2-yl)-5*H*-5λ$^4$,6λ$^4$-dipyrrolo[1,2-*c*:2',1'-*f*][1,3,2]diazaborinin-3-yl)-3-oxo-7,10,13-trioxa-4-azahexadecan-16-oate (11)**

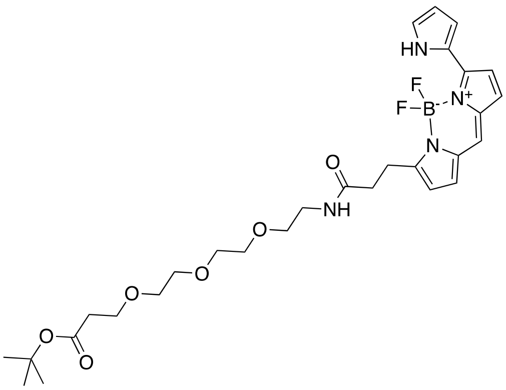


To a flask was added *tert*-butyl 3-(2-(2-(2-aminoethoxy)ethoxy)ethoxy)propanoate (6.5 mg, 1.0 equiv., 23 µmol), 2,5-dioxopyrrolidin-1-yl 3-(5,5-difluoro-7-(1*H*-pyrrol-2-yl)-5*H*-5λ$^4$,6λ$^4$-dipyrrolo[1,2-c:2',1'-*f*][1,3,2]diazaborinin-3-yl)propanoate (10 mg, 1.0 equiv., 23 µmol), and DIPEA (25 µL, 6.0 equiv., 0.14 mmol) in DMF (0.47 mL, 0.05 M). The reaction was concentrated *in vacuo* and the crude material was purified by column chromatography (silica, 0-10% MeOH in $CH_2Cl_2$) to yield the desired product as a purple solid (14 mg, 100 %). $^1$H NMR (400 MHz, MeOD-*d*$_4$) δ 7.99 (s, 1H), 7.24 (dd, *J* = 16.1, 6.9 Hz, 4H), 7.03 (d, *J* = 4.7 Hz, 1H), 6.93 (s, 1H), 6.39 – 6.34 (m, 1H), 3.74 – 3.51 (m, 14H), 3.40 (d, *J* = 5.6 Hz, 2H), 2.71 – 2.63 (m, 2H), 2.53 – 2.41 (m, 2H), 1.45 (dd, *J* = 10.4, 1.3 Hz, 9H).

**1-(5,5-difluoro-7-(1*H*-pyrrol-2-yl)-5*H*-5λ$^4$,6λ$^4$-dipyrrolo[1,2-*c*:2',1'-*f*][1,3,2]diazaborinin-3-yl)-3-oxo-7,10,13-trioxa-4-azahexadecan-16-oic acid (12)**

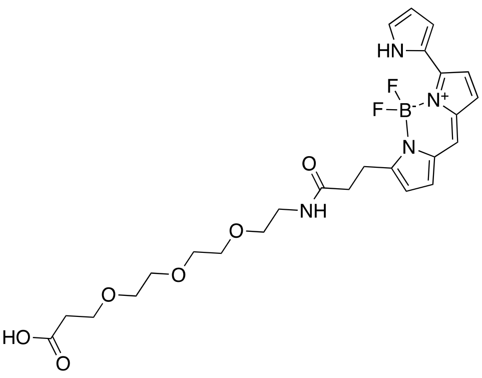


To a flask was added *tert*-butyl 1-(5,5-difluoro-7-(1*H*-pyrrol-2-yl)-5*H*-5λ$^4$,6λ$^4$-dipyrrolo[1,2-*c*:2',1'-*f*][1,3,2]diazaborinin-3-yl)-3-oxo-7,10,13-trioxa-4-azahexadecan-16-oate (**11**) (14 mg, 1.0 equiv., 24 µmol) in 20% TFA in $CH_2Cl_2$ (1 mL) and the reaction stirred at rt for 16 hrs. The reaction was concentrated *in vacuo* and used in the next step without further purification (7.1 mg).

**_N_-(2-(2-(2-(3-(4-(4-((6-(1_H_-indazol-6-yl)imidazo[1,2-_a_]pyrazin-8-yl)amino)phenyl)piperazin-1-yl)-3-oxopropoxy)ethoxy)ethoxy)ethyl)-3-(5,5-difluoro-7-(1_H_-pyrrol-2-yl)-5_H_-5λ$^4$,6λ$^4$-dipyrrolo[1,2-_c_:2',1'-_f_][1,3,2]diazaborinin-3-yl)propenamide (2)**

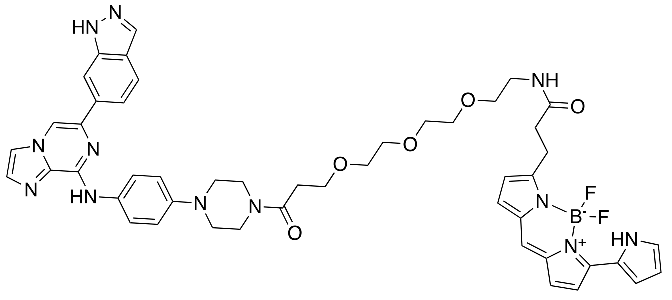


To a flask was added 1-(5,5-difluoro-7-(1_H_-pyrrol-2-yl)-5_H_-5λ$^4$,6λ$^4$-dipyrrolo[1,2-_c_:2',1'-_f_][1,3,2]diazaborinin-3-yl)-3-oxo-7,10,13-trioxa-4-azahexadecan-16-oic acid (**12**) (7.1 mg, 1.0 equiv., 13 µmol), DIPEA (7.4 µL, 3.2 equiv., 43 µmol), and TBTU (6.4 mg, 1.5 equiv., 20 µmol) in DMF (130 µL, 0.1M) and the reaction stirred at rt for 30 mins. (**1**) (10 mg, 1.0 equiv., 13 µmol) was added and the reaction was stirred at rt for 16 hrs. The reaction was concentrated and purified via preparative-HPLC (10-100% MeOH in $H_2O$ (+ 0.01% TFA) to yield the desired product as a purple solid (0.5 mg, 3%). $^1$H NMR (400 MHz, $CDCl_3$) δ 10.28 (s, 1H), 9.98 (s, 1H), 8.25 (s, 1H), 8.11 (s, 1H), 8.00 (s, 1H), 7.92 (d, _J_ = 9.0 Hz, 2H), 7.81 (d, _J_ = 8.5 Hz, 1H), 7.70 (d, _J_ = 1.7 Hz, 1H), 7.62 (d, _J_ = 1.7 Hz, 1H), 7.52 (d, _J_ = 4.5 Hz, 1H), 7.12 (d, _J_ = 8.8 Hz, 2H), 7.05 (s, 1H), 6.95 (d, _J_ = 4.6 Hz, 1H), 6.91 – 6.86 (s, 2H), 6.76 (dd, _J_ = 7.2, 4.3 Hz, 2H), 6.30 (s, 1H), 6.24 (d, _J_ = 4.0 Hz, 1H), 3.86 (s, 2H), 3.81 – 3.72 (m, 4H), 3.58 (d, _J_ = 13.0 Hz, 8H), 3.52 (t, _J_ = 4.9 Hz, 2H), 3.44 (d, _J_ = 5.1 Hz, 2H), 3.29 (t, _J_ = 7.6 Hz, 2H), 3.24 (s, 2H), 3.19 (s, 2H), 2.65 – 2.57 (m, 4H). LCMS calculated for $C_{48}H_{52}BF_2N_{12}O_5$ [M + H]$^+$: 925.42. Found: 925.30

**6-((2-chloro-5-fluoropyrimidin-4-yl)amino)-2,2-dimethyl-2_H_-pyrido[3,2-_b_][1,4]oxazin-3(4_H_)-one (13)**

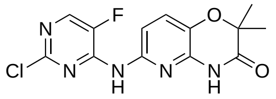


To a flask was added 2,4-dichloro-5-fluoropyrimidine (432 mg, 1.0 equiv., 2.6 mmol), 6-amino-2,2-dimethyl-2_H_-pyrido[3,2-_b_][1,4]oxazin-3(4_H_)-one (500 mg, 1.0 equiv., 2.6 mmol), MeOH (4.31 mL), and $H_2O$ (933 µL) and the reaction was heated at 80 °C for 16 hrs. The reaction was cooled and the precipitate was filtered under vacuum with MeOH to yield the desired product as a white solid (543 mg, 64.9 %). $^1$H NMR (400 MHz, DMSO) δ 11.15 (s, 1H), 10.11 (s, 1H), 8.36 (d, _J_ = 3.3 Hz, 1H), 7.51 – 7.39 (m, 2H), 1.43 (s, 6H). $^{13}$C NMR (214 MHz, DMSO) δ 169.62, 152.83, 150.90, 150.85, 145.68, 144.46, 143.06, 142.46, 142.36, 125.88, 111.38, 77.99, 23.54. LCMS calculated for $C_{13}H_{12}ClFN_5O_2$ [M + H]$^+$: 324.06. Found: 324.13.

**Ethyl 2-(3-((4-((2,2-dimethyl-3-oxo-3,4-dihydro-2_H_-pyrido[3,2-_b_][1,4]oxazin-6-yl)amino)-5-fluoropyrimidin-2-yl)amino)phenoxy)acetate (14)**

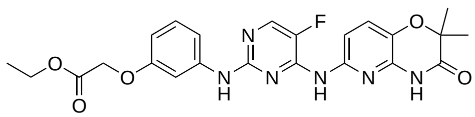

To a microwave vial was added ethyl 2-(3-aminophenoxy)acetate (110 mg, 1.2 equiv., 563 µmol), 6-((2-chloro-5-fluoropyrimidin-4-yl)amino)-2,2-dimethyl-2*H*-pyrido[3,2-*b*][1,4]oxazin-3(4*H*)-one (152 mg, 1.0 equiv., 470 µmol), and EtOH (2 mL). The reaction was degassed with Ar and heated to 80 °C for 16 hrs. The reaction was cooled and the precipitate filtered with ethanol to yield the desired product as a white solid (132 mg, 58%). $^{1}$H NMR (400 MHz, DMSO) δ 11.09 (s, 1H), 9.26 (s, 2H), 8.19 – 8.12 (m, 1H), 7.65 – 7.60 (m, 1H), 7.43 – 7.33 (m, 2H), 7.31 – 7.26 (m, 1H), 7.14 – 7.06 (m, 1H), 6.48 – 6.43 (m, 1H), 4.72 – 4.64 (m, 2H), 4.22 – 4.13 (m, 2H), 1.48 – 1.40 (m, 6H), 1.27 – 1.17 (m, 3H). $^{13}$C NMR (214 MHz, DMSO) δ 169.69, 168.76, 157.85, 155.27, 149.17, 144.29, 141.97, 140.04, 139.87, 133.81, 129.05, 125.75, 111.79, 110.85, 106.28, 105.28, 77.85, 64.60, 60.59, 23.47, 14.05. LCMS calculated for $C_{23}H_{24}FN_6O_5$ [M + H]$^+$: 483.17. Found: 482.87.

**2-(3-((4-((2,2-dimethyl-3-oxo-3,4-dihydro-2*H*-pyrido[3,2-*b*][1,4]oxazin-6-yl)amino)-5-fluoropyrimidin-2-yl)amino)phenoxy)acetic acid (15)**

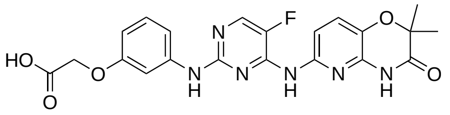

To a flask was added ethyl 2-(3-((4-((2,2-dimethyl-3-oxo-3,4-dihydro-2*H*-pyrido[3,2-*b*][1,4]oxazin-6-yl)amino)-5-fluoropyrimidin-2-yl)amino)phenoxy)acetate (128 mg, 265 µmol, 1.0 equiv.), and LiOH (10 mg, 398 µmol, 1.5 equiv.) in THF (1 mL) and $H_2O$ (239 µL, 50.0 equiv.) and the reaction stirred at 50 ºC for 16 hrs. The reaction was cooled, and HCl (1M) was added. The organics were extracted with EtOAc, washed with brine, and dried with sodium sulfate to yield the desired product as a white solid (66 mg, 55%). $^{1}$H NMR (400 MHz, DMSO) δ 11.08 (s, 1H), 9.29 (d, *J* = 9.1 Hz, 2H), 8.15 (d, *J* = 3.5 Hz, 1H), 7.63 (d, *J* = 8.5 Hz, 1H), 7.40 (d, *J* = 8.5 Hz, 1H), 7.34 – 7.25 (m, 2H), 7.10 (t, *J* = 8.1 Hz, 1H), 6.49 – 6.41 (m, 1H), 4.58 (s, 2H), 1.43 (s, 6H). LCMS calculated for $C_{21}H_{20}FN_6O_5$ [M + H]$^+$: 455.14. Found: 455.08.

***Tert*-butyl (2-(2-(3-((4-((2,2-dimethyl-3-oxo-3,4-dihydro-2*H*-pyrido[3,2-*b*][1,4]oxazin-6-yl)amino)-5-fluoropyrimidin-2-yl)amino)phenoxy)acetamido)ethyl)carbamate (16)**

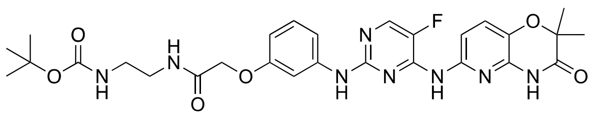

To a flask was added 2-(3-((4-((2,2-dimethyl-3-oxo-3,4-dihydro-2*H*-pyrido[3,2-*b*][1,4]oxazin-6-yl)amino)-5-fluoropyrimidin-2-yl)amino)phenoxy)acetic acid (25 mg, 54 µmol, 1.0 equiv.), DIPEA (30 µL, 173 µmol, 3.2 equiv.), and TBTU (23 mg, 70 µmol, 1.3 equiv.) in DMF (540 µL, 0.1 M). The reaction was allowed to stir at rt for 20 mins followed by addition of *tert*-butyl (2-aminoethyl)carbamate (9 mg, 54 µmol, 1.0 equiv.). The reaction was stirred at rt for 72 hrs. The crude material was concentrated *in vacuo* and purified via column chromatography (silica, 0-20% MeOH in $CH_2Cl_2$) to yield the desired product as a white solid (16 mg, 51%). $^{1}$H NMR (400 MHz, DMSO) δ 11.07 (s, 1H), 9.27 (s, 1H), 9.21 (s, 1H), 8.14 (d, *J* = 3.4 Hz, 1H), 7.64 (d, *J* = 8.5 Hz, 1H), 7.41 – 7.35 (m, 2H), 7.29 (d, *J* = 8.2 Hz, 1H), 7.11 (t, *J* = 8.2 Hz, 1H), 6.49 (dd, *J* = 8.2, 2.4 Hz, 1H), 4.38 (s, 2H), 3.16 (t, *J* = 6.0 Hz, 2H), 3.05 – 2.,99 (m, 2H), 1.43 (s, 6H), 1.37 (s, 9H). LCMS calculated for $C_{28}H_{34}FN_8O_6$ [M + H]$^+$: 597.25. Found: 597.34.

***N*-(2-aminoethyl)-2-(3-((4-((2,2-dimethyl-3-oxo-3,4-dihydro-2*H*-pyrido[3,2-*b*][1,4]oxazin-6-yl)amino)-5-fluoropyrimidin-2-yl)amino)phenoxy)acetamide (17)**

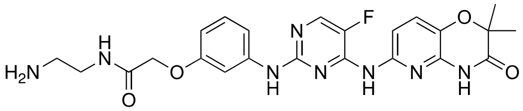

To a flask was added *tert*-butyl (2-(2-(3-((4-((2,2-dimethyl-3-oxo-3,4-dihydro-2*H*-pyrido[3,2-*b*][1,4]oxazin-6-yl)amino)-5-fluoropyrimidin-2-yl)amino)phenoxy)acetamido)ethyl)carbamate (17 mg, 28 µmol, 1.0 equiv.) and TFA (277 µL) in $CH_2Cl_2$ (1 mL) and the reaction stirred at rt for 16 hrs. The reaction was concentrated *in vacuo* to yield the desired product (7 mg, 41%). $^1$H NMR (400 MHz, DMSO) δ 11.08 (s, 1H), 9.33 (d, *J* = 5.7 Hz, 2H), 8.20 (t, *J* = 5.9 Hz, 1H), 8.15 (d, *J* = 3.5 Hz, 1H), 7.70 (s, 2H), 7.61 (d, *J* = 8.5 Hz, 1H), 7.41 – 7.35 (m, 2H), 7.29 (d, *J* = 8.2 Hz, 1H), 7.12 (t, *J* = 8.2 Hz, 1H), 6.52 (dd, *J* = 8.2, 2.4 Hz, 1H), 4.42 (s, 2H), 3.38 (q, *J* = 6.2 Hz, 2H), 2.89 (q, *J* = 5.9 Hz, 2H), 1.42 (s, 6H). LCMS calculated for $C_{23}H_{26}FN_8O_6$ $[M + H]^+$: 497.20. Found: 497.23.

**3-(5,5-difluoro-7-(1*H*-pyrrol-2-yl)-5*H*-5λ$^4$,6λ$^4$-dipyrrolo[1,2-*c*:2',1'-*f*][1,3,2]diazaborinin-3-yl)-*N*-(2-(2-(3-((4-((2,2-dimethyl-3-oxo-3,4-dihydro-2*H*-pyrido[3,2-*b*][1,4]oxazin-6-yl)amino)-5-fluoropyrimidin-2-yl)amino)phenoxy)acetamido)ethyl)propenamide (6)**

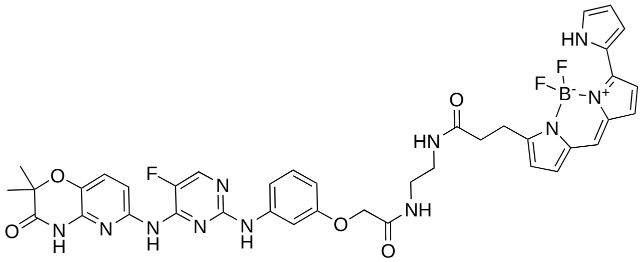

To a flask was added *N*-(2-aminoethyl)-2-(3-((4-((2,2-dimethyl-3-oxo-3,4-dihydro-2*H*-pyrido[3,2-*b*][1,4]oxazin-6-yl)amino)-5-fluoropyrimidin-2-yl)amino)phenoxy)acetamide 2,2,2-trifluoroacetate (7 mg, 0.01 mmol, 1.0 equiv.), 2,5-dioxopyrrolidin-1-yl 3-(5,5-difluoro-7-(1*H*-pyrrol-2-yl)-5*H*-5λ$^4$,6λ$^4$-dipyrrolo[1,2-*c*:2',1'-*f*][1,3,2]diazaborinin-3-yl)propanoate (5 mg, 0.01 mmol, 1.0 equiv.), and DIPEA (10 µL, 6.0 equiv.) in DMF (0.1 mL, 0.1 M). The reaction was stirred at rt for 10 mins, concentrated *in vacuo*, and purified by prep-HPLC (10-100% MeOH in $H_2O$ (0.01% TFA) to yield the desired product as a purple solid (3.7 mg, 40 %). $^1$H NMR (400 MHz, MeOD-*d*$_4$) δ 7.95 (d, *J* = 8.7 Hz, 1H), 7.93 (d, *J* = 3.5 Hz, 1H), 7.41 – 7.39 (m, 1H), 7.23 (d, *J* = 6.3 Hz, 1H), 7.20 (d, *J* = 5.2 Hz, 1H), 7.18 – 7.14 (m, 2H), 7.14 – 7.10 (m, 3H), 6.95 (d, *J* = 4.6 Hz, 1H), 6.83 (d, *J* = 3.9 Hz, 1H), 6.65 – 6.59 (m, 1H), 6.31 (dd, *J* = 3.9, 2.5 Hz, 1H), 6.24 (d, *J* = 4.0 Hz, 1H), 4.49 (s, 2H), 3.42 – 3.35 (m, 2H), 3.22 (t, *J* = 7.8 Hz, 2H), 2.60 – 2.52 (m, 2H), 1.49 (s, 6H). LCMS calculated for $C_{39}H_{38}BF_3N_{11}O_5$ $[M + H]^+$: 808.30. Found: 808.6.

***N*-(2-(2-(2-(2-aminoethoxy)ethoxy)ethoxy)ethyl)-2-(3-((4-((2,2-dimethyl-3-oxo-3,4-dihydro-2*H*-pyrido[3,2-*b*][1,4]oxazin-6-yl)amino)-5-fluoropyrimidin-2-yl)amino)phenoxy)acetamide (18)**

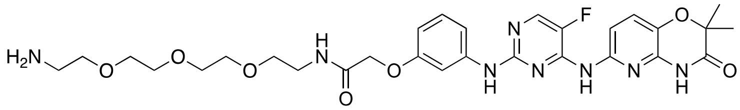

To a flask was added 2-(3-((4-((2,2-dimethyl-3-oxo-3,4-dihydro-2*H*-pyrido[3,2-*b*][1,4]oxazin-6-yl)amino)-5-fluoropyrimidin-2-yl)amino)phenoxy)acetic acid (20 mg, 1.0 equiv., 44 µmol), DIPEA (17 mg, 23 µL, 3.0 equiv., 0.13 mmol), and TBTU (18 mg, 1.3 equiv., 57 µmol) in DMF (0.44 mL, 0.1 M) and the reaction stirred at rt for approx. 5 mins. To this was added *tert*-butyl (2-(2-(2-(2-

aminoethoxy)ethoxy)ethoxy)ethyl)carbamate (13 mg, 1.0 equiv., 44 µmol) and the reaction stirred for 16 hrs. The reaction was concentrated *in vacuo* and purified by column chromatography (silica, 0-10% MeOH in $CH_2Cl_2$) to yield the desired product as a yellow amorphous solid *tert*-butyl (1-(3-((4-((2,2-dimethyl-3-oxo-3,4-dihydro-2*H*-pyrido[3,2-*b*][1,4]oxazin-6-yl)amino)-5-fluoropyrimidin-2-yl)amino)phenoxy)-2-oxo-6,9,12-trioxa-3-azatetradecan-14-yl)carbamate (28.6 mg, 89 %). LCMS calculated for $C_{34}H_{46}FN_8O_9$ $[M + H]^+$: 729.33. Found: 729.25.

To a flask was added *tert*-butyl (1-(3-((4-((2,2-dimethyl-3-oxo-3,4-dihydro-2*H*-pyrido[3,2-*b*][1,4]oxazin-6-yl)amino)-5-fluoropyrimidin-2-yl)amino)phenoxy)-2-oxo-6,9,12-trioxa-3-azatetradecan-14-yl)carbamate (28.6 mg, 39 µmol, 1.0 equiv.) and 20% TFA in $CH_2Cl_2$ (1 mL), and the reaction stirred at rt for 16 hrs. The reaction was concentrated *in vacuo* to yield the desired product as a yellow solid (25 mg, quant). LCMS calculated for $C_{29}H_{38}FN_8O_7$ $[M + H]^+$: 629.28. Found: 629.17.

**3-(5,5-difluoro-7-(1*H*-pyrrol-2-yl)-5*H*-5λ⁴,6λ⁴-dipyrrolo[1,2-*c*:2',1'-*f*][1,3,2]diazaborinin-3-yl)-*N*-(1-(3-((4-((2,2-dimethyl-3-oxo-3,4-dihydro-2*H*-pyrido[3,2-*b*][1,4]oxazin-6-yl)amino)-5-fluoropyrimidin-2-yl)amino)phenoxy)-2-oxo-6,9,12-trioxa-3-azatetradecan-14-yl)propenamide (5)**

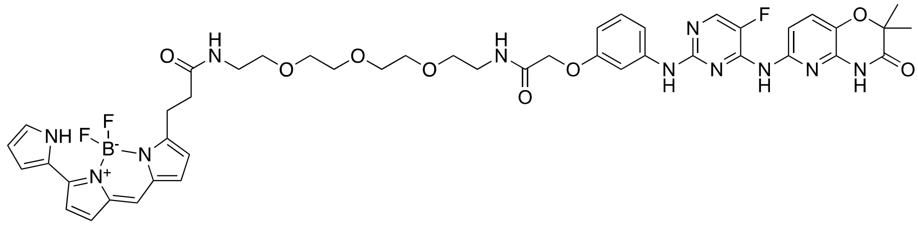

To a flask was added *N*-(2-(2-(2-(2-aminoethoxy)ethoxy)ethoxy)ethyl)-2-(3-((4-((2,2-dimethyl-3-oxo-3,4-dihydro-2*H*-pyrido[3,2-*b*][1,4]oxazin-6-yl)amino)-5-fluoropyrimidin-2-yl)amino)phenoxy)acetamide 2,2,2-trifluoroacetate (5 mg, 7 µmol, 1.0 equiv.), 2,5-dioxopyrrolidin-1-yl 3-(5,5-difluoro-7-(1*H*-pyrrol-2-yl)-5*H*-5$\lambda^4$,6$\lambda^4$-dipyrrolo[1,2-*c*:2',1'-*f*][1,3,2]diazaborinin-3-yl)propanoate (3 mg, 7 µmol, 1.0 equiv.), and DIPEA (7 µL, 42 µmol, 6.0 equiv.) in DMF (0.5 mL) and the reaction stirred at rt for 30 mins. The reaction was concentrated *in vacuo* and purified by prep-HPLC (10-100% MeOH in $H_2O$ (0.01% TFA) to yield the desired product as a purple solid (2 mg, 30 %). $^1H$ NMR (400 MHz, $CDCl_3$) δ 10.38 (s, 1H), 8.99 (s, 1H), 8.02 (s, 1H), 8.00 (d, *J* = 3.0 Hz, 1H), 7.90 (d, *J* = 8.7 Hz, 1H), 7.33 (d, *J* = 2.3 Hz, 1H), 7.24 – 7.19 (m, 2H), 7.16 (d, *J* = 9.8 Hz, 2H), 7.00 (d, *J* = 4.6 Hz, 1H), 6.94 (s, 1H), 6.83 (d, *J* = 4.6 Hz, 1H), 6.79 (d, *J* = 4.0 Hz, 1H), 6.56 (d, *J* = 8.2 Hz, 1H), 6.38 (s, 1H), 6.37 – 6.33 (m, 1H), 6.26 (d, *J* = 4.0 Hz, 1H), 4.49 (s, 2H), 3.58 (s, 4H), 3.54 – 3.48 (m, 10H), 3.43 (q, *J* = 5.2 Hz, 2H), 3.31 (t, *J* = 7.7 Hz, 2H), 2.96 (s, 3H), 2.88 (d, *J* = 0.6 Hz, 3H), 2.60 (t, *J* = 7.7 Hz, 2H). LCMS calculated for $C_{45}H_{50}BF_3N_{11}O_8$ $[M + H]^+$: 940.38. Found: 940.0.

**(1*S*,4*R*)-*N*-(15-(5,5-difluoro-7-(1*H*-pyrrol-2-yl)-5*H*-5λ⁴,6λ⁴-dipyrrolo[1,2-*c*:2',1'-*f*][1,3,2]diazaborinin-3-yl)-13-oxo-3,6,9-trioxa-12-azapentadecyl)-4-hydroxy-2,2-dimethyl-4-(5-(3-methyl-5-((4-methylpyrimidin-2-yl)amino)phenyl)thiazol-2-yl)cyclohexane-1-carboxamide (7)**

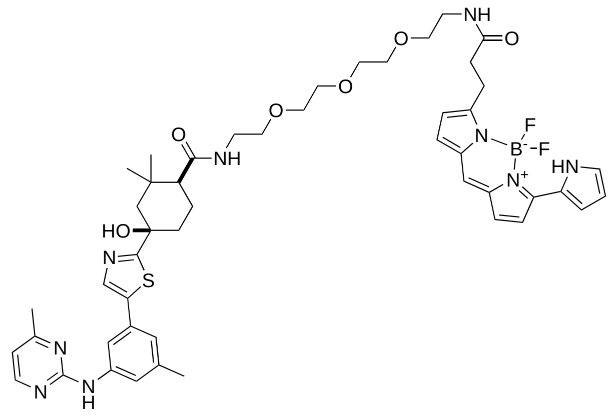

To a flask was added (1*S*,4*R*)-4-hydroxy-2,2-dimethyl-4-(5-(3-methyl-5-((4-methylpyrimidin-2-yl)amino)phenyl)thiazol-2-yl)cyclohexane-1-carboxylic acid (5.0 mg, 11 μmol, 1.0 equiv.), DIPEA (6.2 μL, 35 μmol, 3.2 equiv.), and HATU (5.5 mg, 14 μmol, 1.3 equiv.), in DMF (110 μL, 0.1 M) and the reaction allowed to stir at rt for 5 mins. To this was added *tert*-butyl (2-(2-(2-(2-aminoethoxy)ethoxy)ethoxy)ethyl)carbamate (3.2 mg, 11 μmol, 1.0 equiv.) and the reaction left to stir at rt for 16 hrs to yield the intermediate *tert*-butyl (1-((1*S*,4*R*)-4-hydroxy-2,2-dimethyl-4-(5-(3-methyl-5-((4-methylpyrimidin-2-yl)amino)phenyl)thiazol-2-yl)cyclohexyl)-1-oxo-5,8,11-trioxa-2-azatridecan-13-yl)carbamate. The reaction was concentrated *in vacuo* and deprotected with 20% TFA in $CH_2Cl_2$ at rt for 1 hr. The intermediate (1*S*,4*R*)-*N*-(2-(2-(2-(2-aminoethoxy)ethoxy)ethoxy)ethyl)-4-hydroxy-2,2-dimethyl-4-(5-(3-methyl-5-((4-methylpyrimidin-2-yl)amino)phenyl)thiazol-2-yl)cyclohexane-1-carboxamide 2,2,2-trifluoroacetate was stirred at rt for 30 mins with 2,5-dioxopyrrolidin-1-yl 3-(5,5-difluoro-7-(1*H*-pyrrol-2-yl)-5*H*-5λ4,6λ4-dipyrrolo[1,2-*c*:2',1'-*f*][1,3,2]diazaborinin-3-yl)propanoate (5.6 mg, 13 μmol, 1.0 equiv.) and DIPEA (14 μL, 35 μmol, 3.2 equiv.) in DMF (130 μL, 0.1 M). The reaction was concentrated *in vacuo* and purified by preparative HPLC (10-100% MeOH in $H_2O$ (+0.01% TFA)) to yield the desired product as a purple solid (6.1 mg, 50% over 3 steps). $^1$H NMR (400 MHz, MeOD-*d*$_4$) δ 10.67 (s, 1H), 8.24 (d, *J* = 5.4 Hz, 1H), 7.87 (s, 1H), 7.84 – 7.82 (m, 1H), 7.37 – 7.34 (m, 1H), 7.19 – 7.11 (m, 5H), 6.96 (d, *J* = 4.6 Hz, 1H), 6.87 (d, *J* = 4.0 Hz, 1H), 6.77 (d, *J* = 5.4 Hz, 1H), 6.34 – 6.30 (m, 1H), 6.29 (d, *J* = 4.0 Hz, 1H), 3.62 – 3.55 (m, 8H), 3.51 (dt, *J* = 11.2, 5.4 Hz, 4H), 3.37 (t, *J* = 5.5 Hz, 3H), 3.25 (d, *J* = 8.1 Hz, 3H), 2.67 – 2.58 (m, 2H), 2.44 (s, 3H), 2.38 – 2.33 (m, 3H), 2.15 (dd, *J* = 12.6, 2.4 Hz, 1H), 1.97 – 1.89 (m, 3H), 1.71 (d, *J* = 14.3 Hz, 1H), 1.57 – 1.49 (m, 1H), 1.20 (s, 3H), 0.99 (s, 3H). LCMS calculated for $C_{48}H_{59}BF_2N_9O_6S$ [M + H]$^+$: 938.43. Found: 938.1.

**(1*S*,4*R*)-*N*-(3-(3-(5,5-difluoro-7-(1*H*-pyrrol-2-yl)-5*H*-5λ4,6λ4-dipyrrolo[1,2-*c*:2',1'-*f*][1,3,2]diazaborinin-3-yl)propanamido)propyl)-4-hydroxy-2,2-dimethyl-4-(5-(3-methyl-5-((4-methylpyrimidin-2-yl)amino)phenyl)thiazol-2-yl)cyclohexane-1-carboxamide (8)**

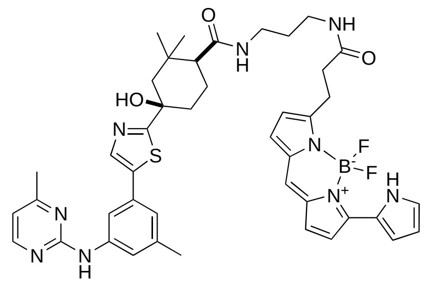

To a flask was added (1*S*,4*R*)-4-hydroxy-2,2-dimethyl-4-(5-(3-methyl-5-((4-methylpyrimidin-2-yl)amino)phenyl)thiazol-2-yl)cyclohexane-1-carboxylic acid (5.0 mg, 11 μmol, 1.0 equiv.), DIPEA (6.2 μL, 35 μmol, 3.2 equiv.), and HATU (5.5 mg, 14 μmol, 1.3 equiv.), in DMF (110 μL, 0.1 M) and the reaction allowed to stir at rt for 5 mins. To this was added *tert*-butyl (3-aminopropyl)carbamate (1.9 mg, 11 μmol, 1.0 equiv.) and the reaction left to stir at rt for 16 hrs to yield the intermediate *tert*-butyl (3-((1*S*,4*R*)-4-hydroxy-2,2-dimethyl-4-(5-(3-methyl-5-((4-methylpyrimidin-2-yl)amino)phenyl)thiazol-2-yl)cyclohexane-1-carboxamido)propyl)carbamate. The reaction was concentrated *in vacuo* and deprotected with 20% TFA in $CH_2Cl_2$ at rt for 1 hr. The intermediate (1*S*,4*R*)-*N*-(3-aminopropyl)-4-hydroxy-2,2-dimethyl-4-(5-(3-methyl-5-((4-methylpyrimidin-2-yl)amino)phenyl)thiazol-2-yl)cyclohexane-1-carboxamide 2,2,2-trifluoroacetate was stirred at rt for 30 mins with 2,5-dioxopyrrolidin-1-yl 3-(5,5-difluoro-7-(1*H*-pyrrol-2-yl)-5*H*-5λ4,6λ4-dipyrrolo[1,2-*c*:2',1'-*f*][1,3,2]diazaborinin-3-yl)propanoate (5.6 mg, 13 μmol, 1.0 equiv.) and DIPEA (14 μL, 35 μmol, 3.2 equiv.) in DMF (130 μL, 0.1 M). The reaction was concentrated *in vacuo* and purified by preparative HPLC (10-100% MeOH in $H_2O$ (+0.01% TFA)) to yield the desired product as a purple solid (5.9 mg, 65% over 3 steps). $^1$H NMR (400 MHz, MeOD-$d_4$) δ 10.71 (s, 1H), 8.28 (d, *J* = 5.1 Hz, 1H), 7.91 – 7.90 (m, 1H), 7.89 (s, 1H), 7.44 – 7.39 (m, 1H), 7.25 – 7.16 (m, 4H), 7.13 – 7.08 (m, 1H), 7.01 (d, *J* = 4.6 Hz, 1H), 6.92 (d, *J* = 4.0 Hz, 1H), 6.74 (dd, *J* = 5.2, 0.5 Hz, 1H), 6.39 – 6.31 (m, 2H), 3.28 – 3.21 (m, 4H), 3.19 – 3.15 (m, 1H), 2.65 (t, *J* = 7.7 Hz, 2H), 2.44 (s, 3H), 2.37 (s, 3H), 2.33 – 2.20 (m, 1H), 2.16 (dd, *J* = 12.6, 2.6 Hz, 1H), 2.05 – 1.93 (m, 3H), 1.74 – 1.67 (m, 3H), 1.57 (dd, *J* = 13.0, 3.0 Hz, 1H), 1.24 (s, 3H), 1.03 (s, 3H). LCMS calculated for $C_{43}H_{49}BF_2N_9O_3S$ $[M + H]^+$: 820.37. Found: 820.7.

#### 2.5 Purity traces and spectra

##### *Tert*-butyl 4-(4-((6-bromoimidazo[1,2-*a*]pyrazin-8-yl)amino)phenyl)piperazine-1-carboxylate (9)

##### $^{1}$H NMR

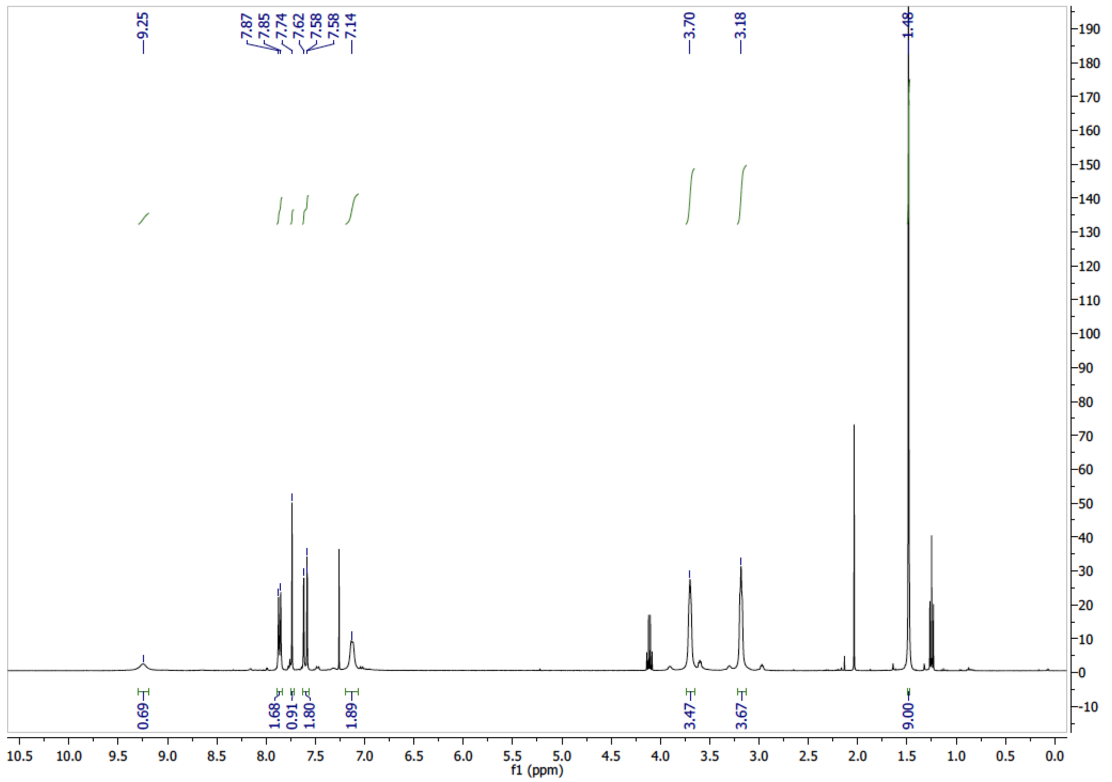


##### LCMS

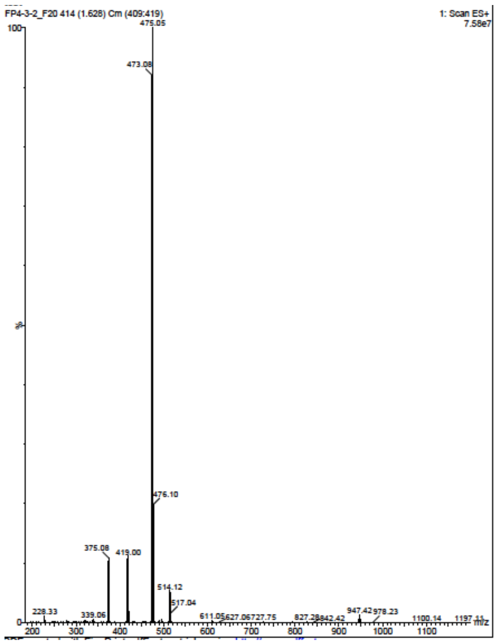

**_Tert_-butyl 4-(4-((6-(1_H_-indazol-6-yl)imidazo[1,2-_a_]pyrazin-8-yl)amino)phenyl)piperazine-1-carboxylate (10)**

**$^{1}$H NMR**

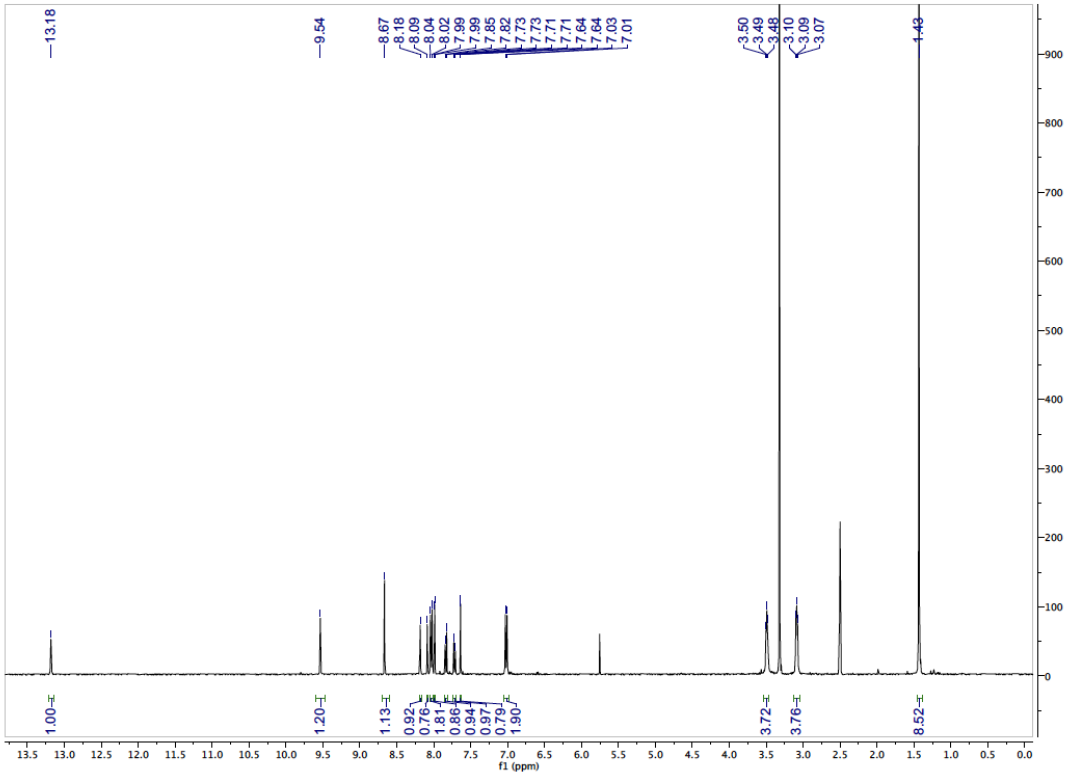

**$^{13}$C NMR**

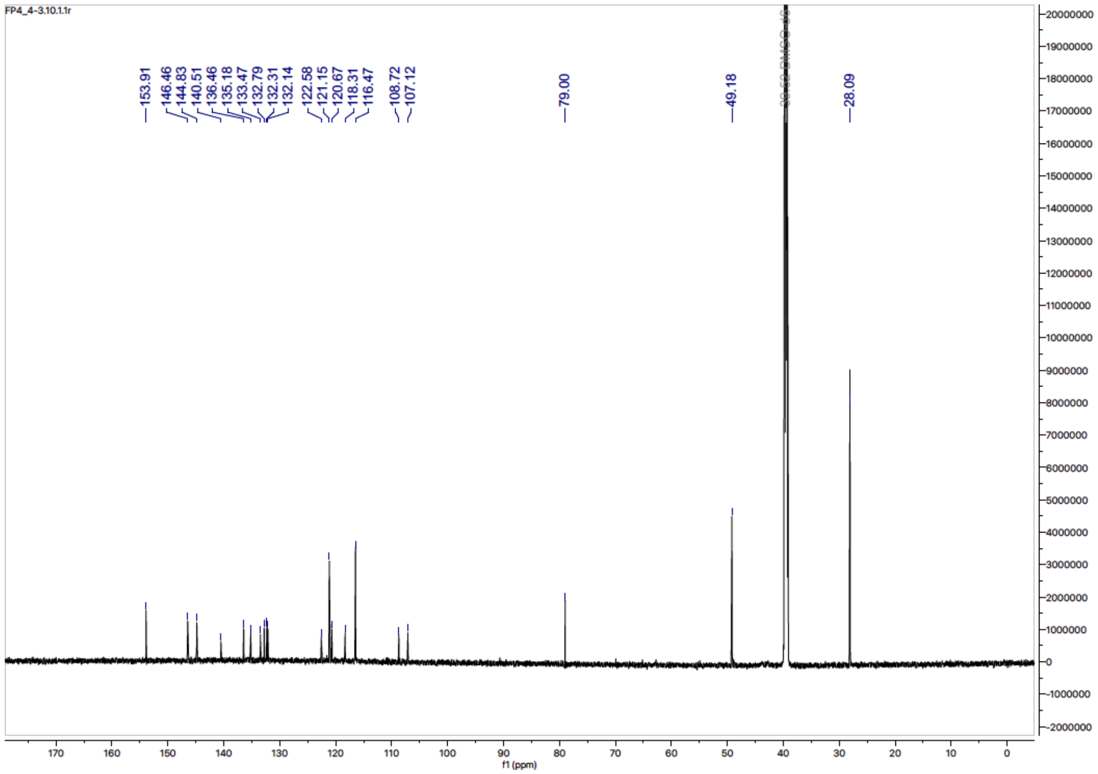

**6-(1-methyl-1*H*-indazol-6-yl)-*N*-(4-(piperazin-1-yl)phenyl)imidazo[1,2-*a*]pyrazin-8-amine (3)**

**$^{1}$H NMR**

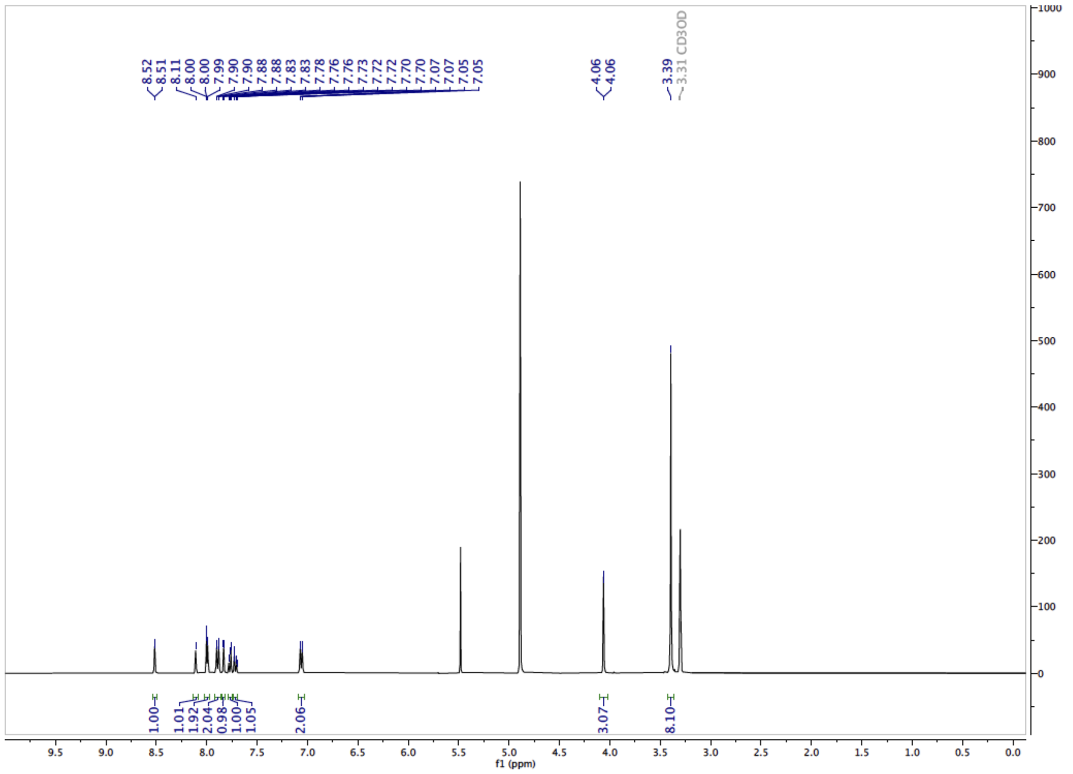

**$^{13}$C NMR**

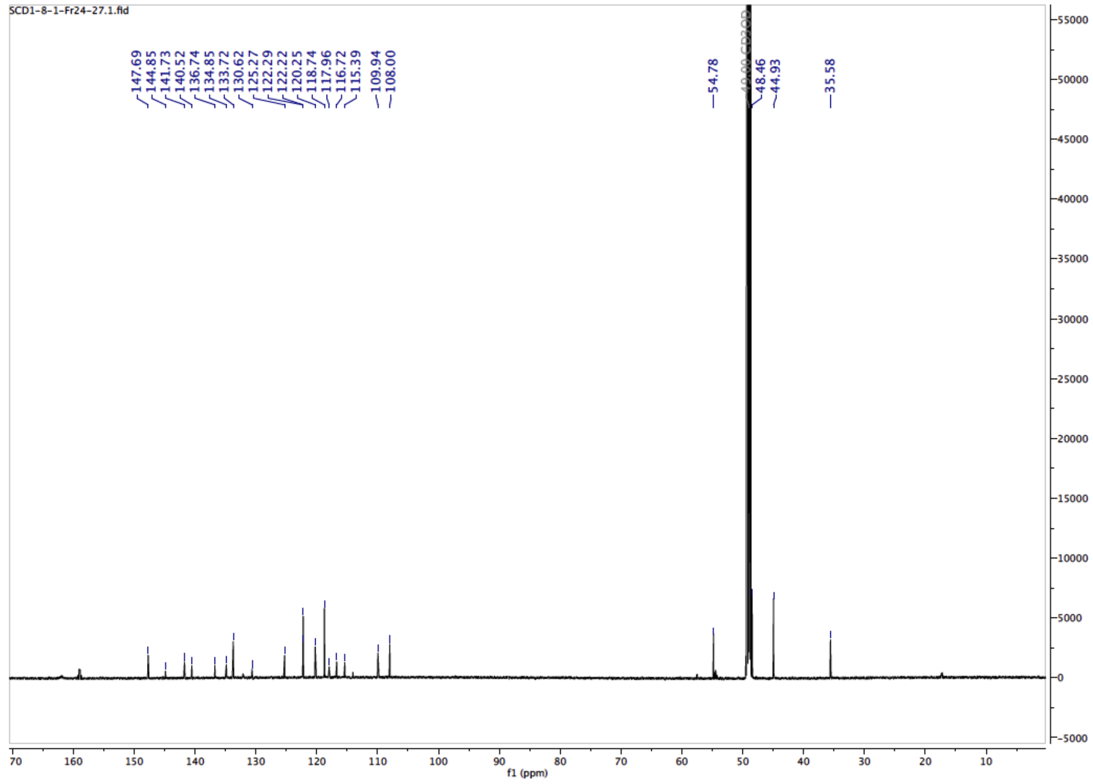

#### LCMS

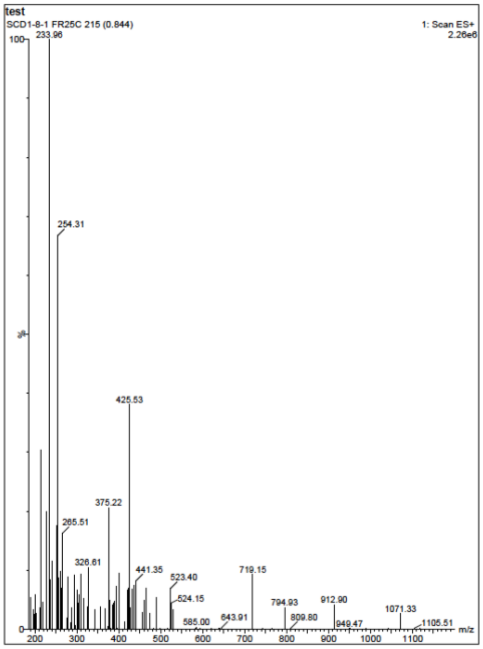


**3-(5,5-difluoro-7-(1*H*-pyrrol-2-yl)-5*H*-5λ$^4$,6λ$^4$-dipyrrolo[1,2-*c*:2',1'-*f*][1,3,2]diazaborinin-3-yl)-1-(4-(4-((6-(1-methyl-1*H*-indazol-6-yl)imidazo[1,2-*a*]pyrazin-8-yl)amino)phenyl)piperazin-1-yl)propan-1-one (4)**

#### $^1$H NMR

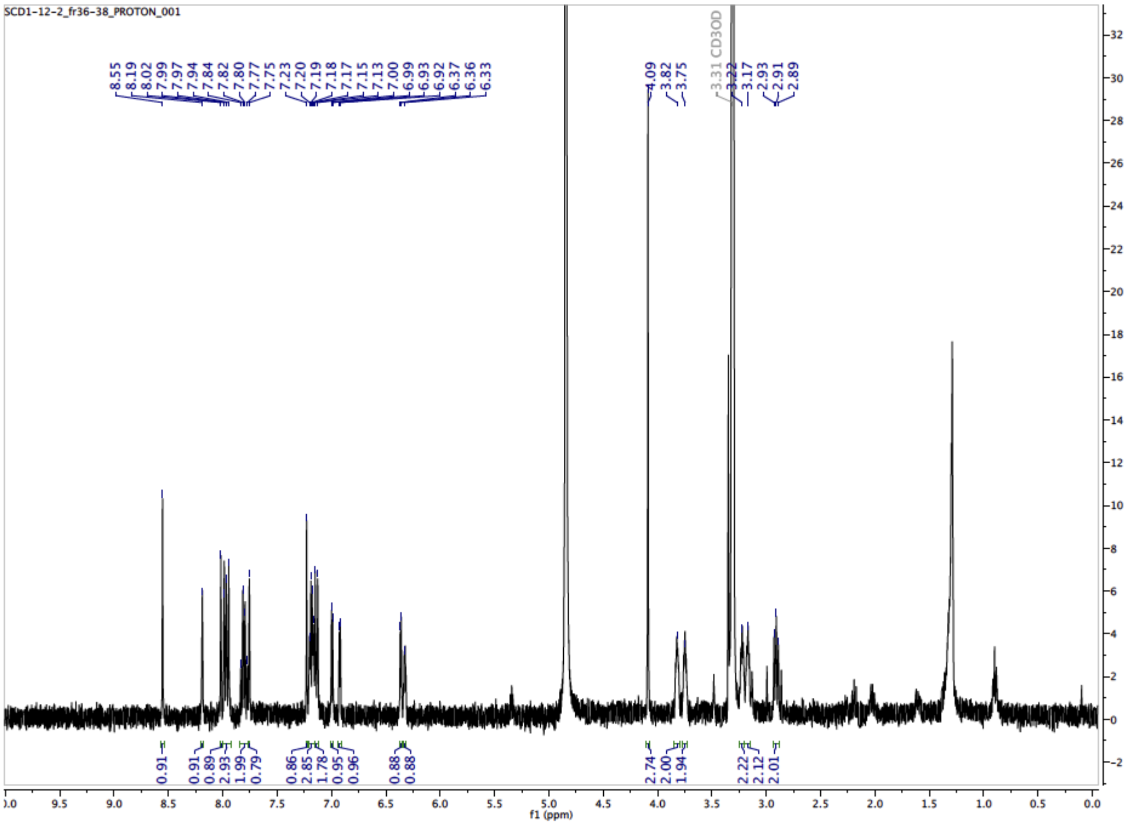

#### LCMS

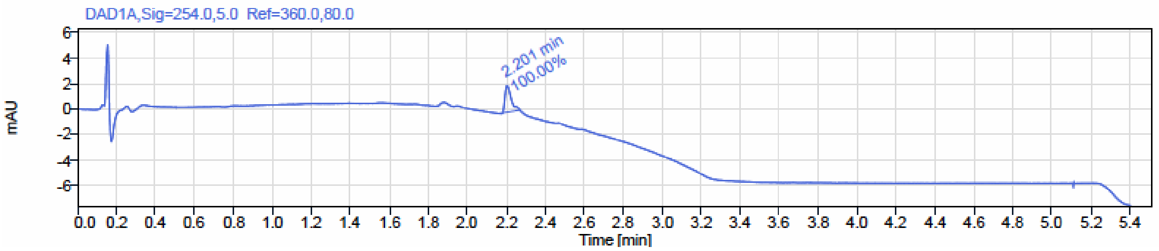


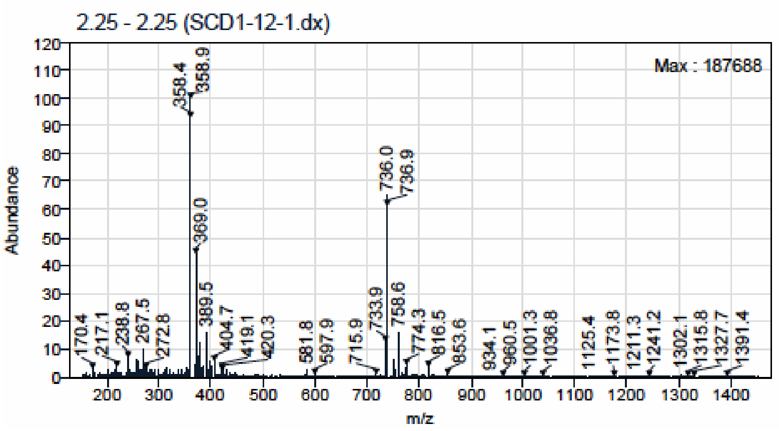


**Tert-butyl 1-(5,5-difluoro-7-(1*H*-pyrrol-2-yl)-5*H*-5λ⁴,6λ⁴-dipyrrolo[1,2-*c*:2',1'-*f*][1,3,2]diazaborinin-3-yl)-3-oxo-7,10,13-trioxa-4-azahexadecan-16-oate (11)**

#### $^{1}$H NMR

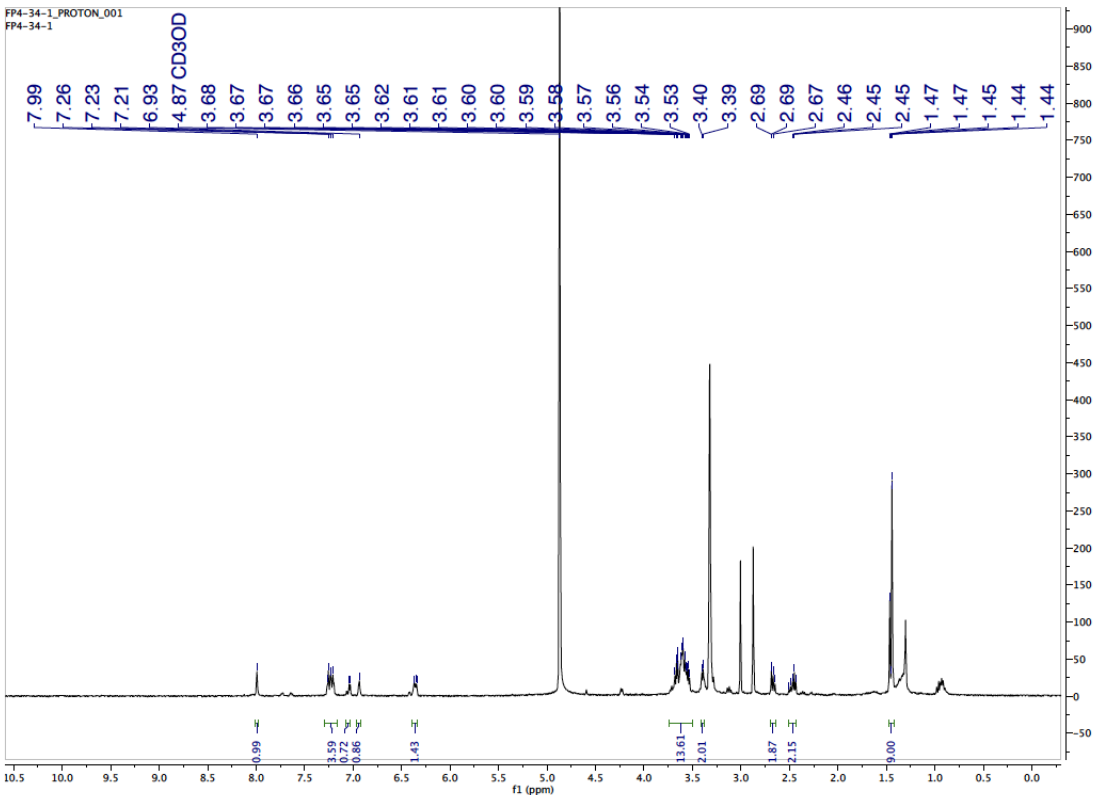

**N-(2-(2-(2-(3-(4-(4-((6-(1*H*-indazol-6-yl)imidazo[1,2-*a*]pyrazin-8-yl)amino)phenyl)piperazin-1-yl)-3-oxopropoxy)ethoxy)ethoxy)ethyl)-3-(5,5-difluoro-7-(1*H*-pyrrol-2-yl)-5*H*-5$\lambda^4$,6$\lambda^4$-dipyrrolo[1,2-*c*:2',1'-*f*][1,3,2]diazaborinin-3-yl)propenamide (2)**

**$^1$H NMR**

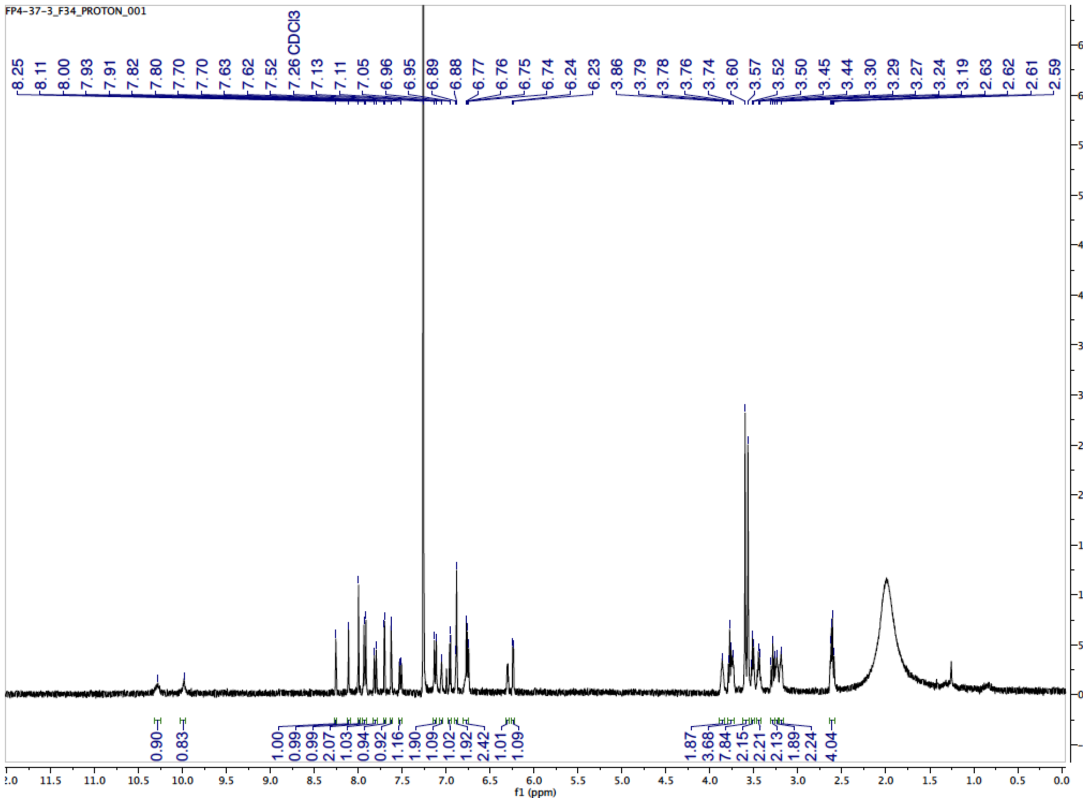


**LCMS**

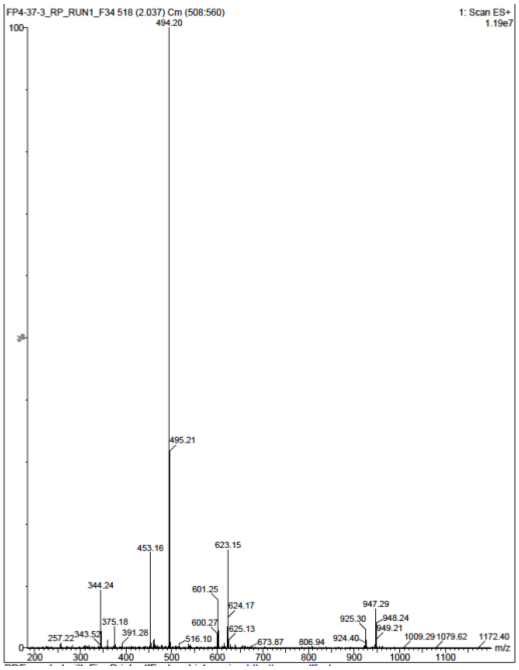

**6-((2-chloro-5-fluoropyrimidin-4-yl)amino)-2,2-dimethyl-2*H*-pyrido[3,2-*b*][1,4]oxazin-3(4*H*)-one (13)**

**$^{1}$H NMR**

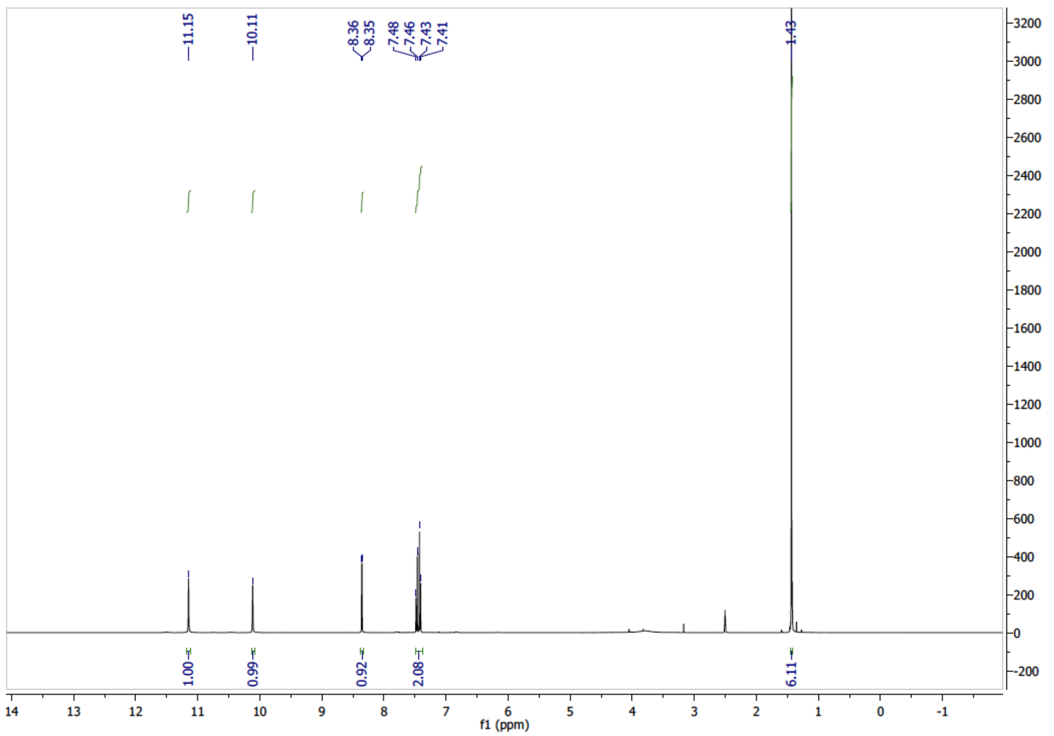

**$^{13}$C NMR**

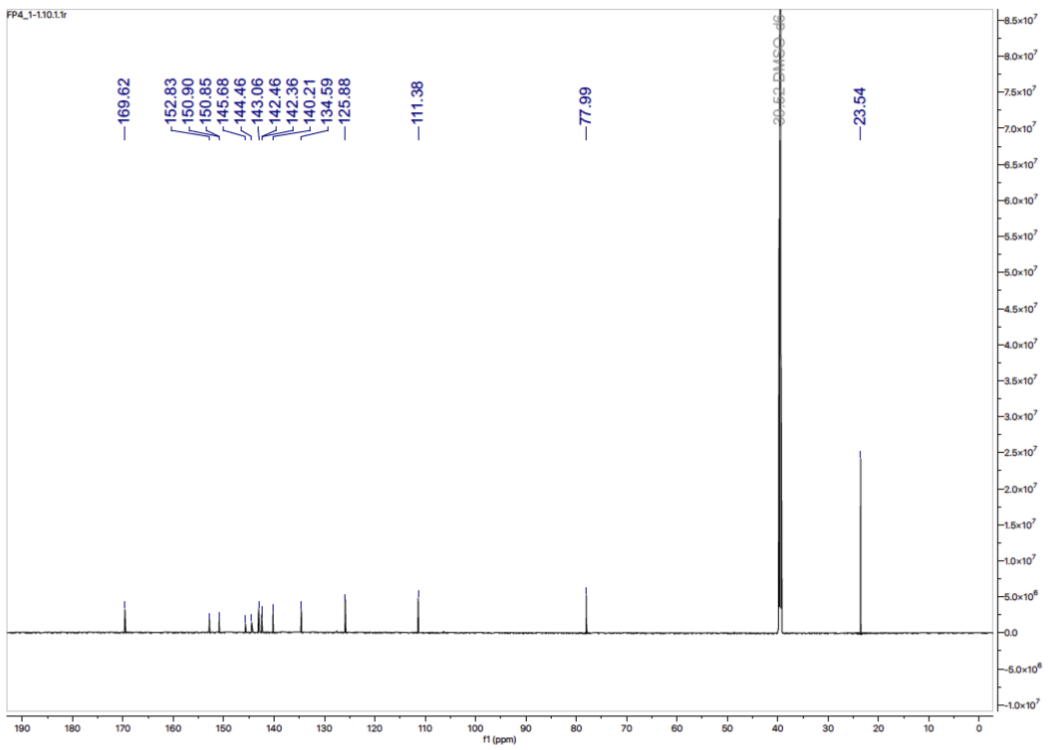

**LCMS**

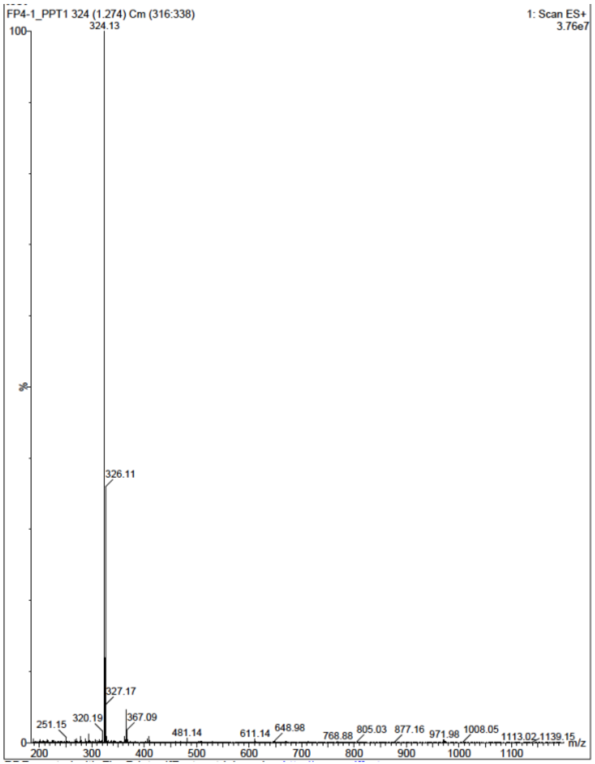


**Ethyl 2-(3-((4-((2,2-dimethyl-3-oxo-3,4-dihydro-2*H*-pyrido[3,2-*b*][1,4]oxazin-6-yl)amino)-5-fluoropyrimidin-2-yl)amino)phenoxy)acetate (14)**

**$^{1}$H NMR**

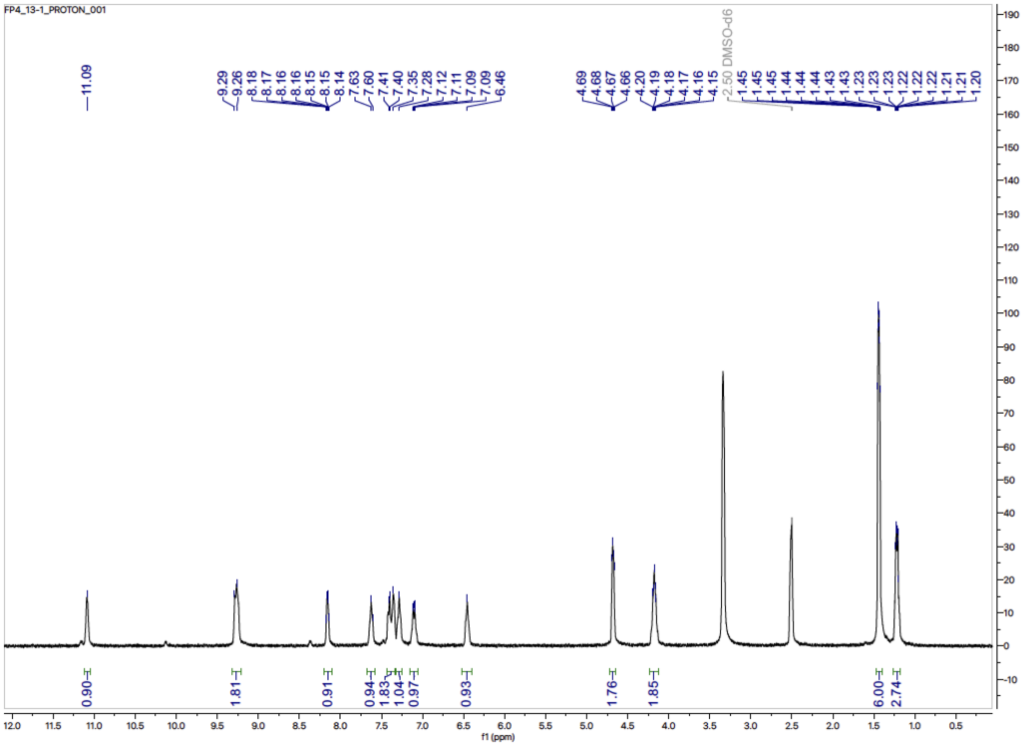

**$^{13}$C NMR**

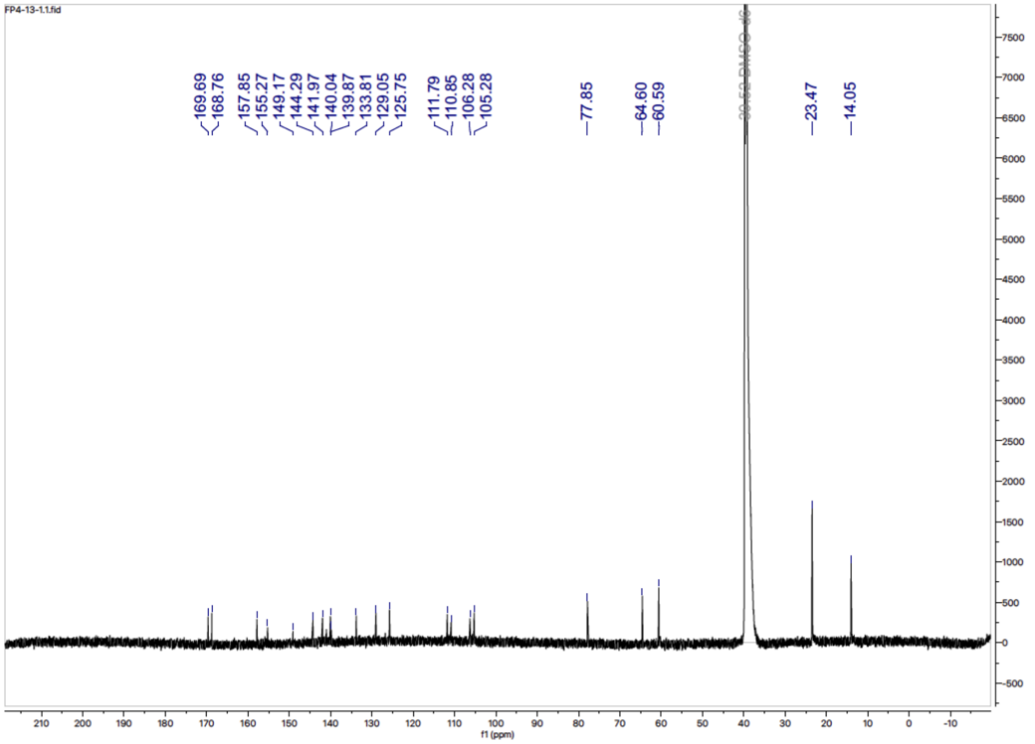

FP4-13-1.1.fid
169.69
168.76
157.85
155.27
149.17
144.29
141.97
140.04
139.87
133.81
129.05
125.75
111.79
110.85
106.28
105.28
77.85
64.60
60.59
23.47
14.05
f1 (ppm)


**LCMS**

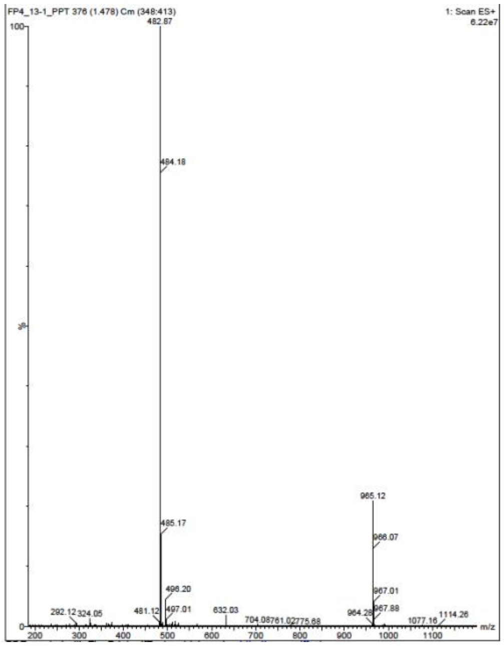

FP4_13-1_PPT 376 (1.478) Cm (348:413)
1: Scan ES+
6.22e7
482.87
484.18
485.17
496.20
497.01
481.12
292.12
324.05
632.03
965.12
966.07
967.01
967.88
964.28
1114.26
1077.16
m/z

**2-(3-((4-((2,2-dimethyl-3-oxo-3,4-dihydro-2*H*-pyrido[3,2-*b*][1,4]oxazin-6-yl)amino)-5-fluoropyrimidin-2-yl)amino)phenoxy)acetic acid (15)**

**$^{1}$H NMR**

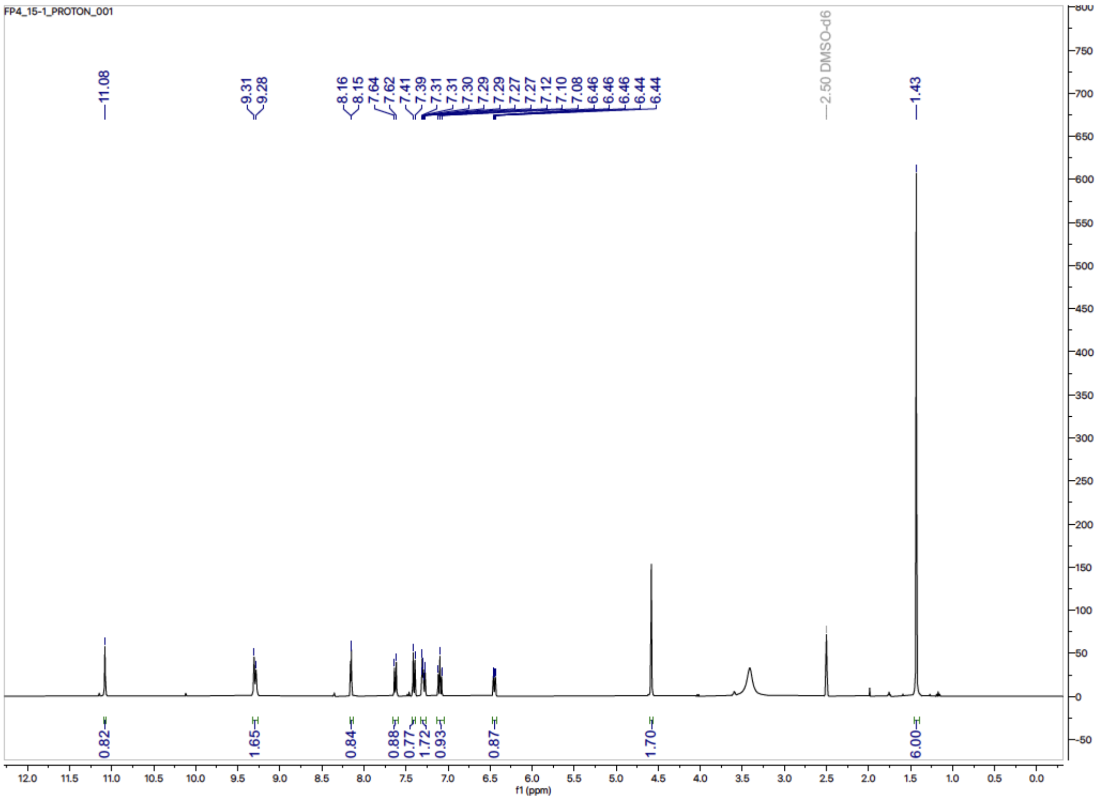


**LCMS**

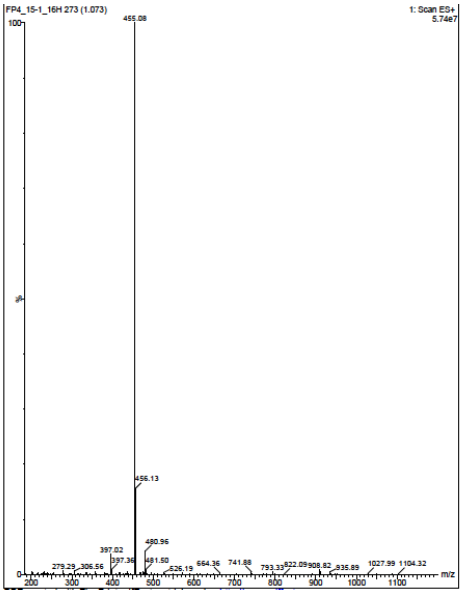

***Tert*-butyl (2-(2-(3-((4-((2,2-dimethyl-3-oxo-3,4-dihydro-2*H*-pyrido[3,2-*b*][1,4]oxazin-6-yl)amino)-5-fluoropyrimidin-2-yl)amino)phenoxy)acetamido)ethyl)carbamate (16)**

**[1]H NMR**

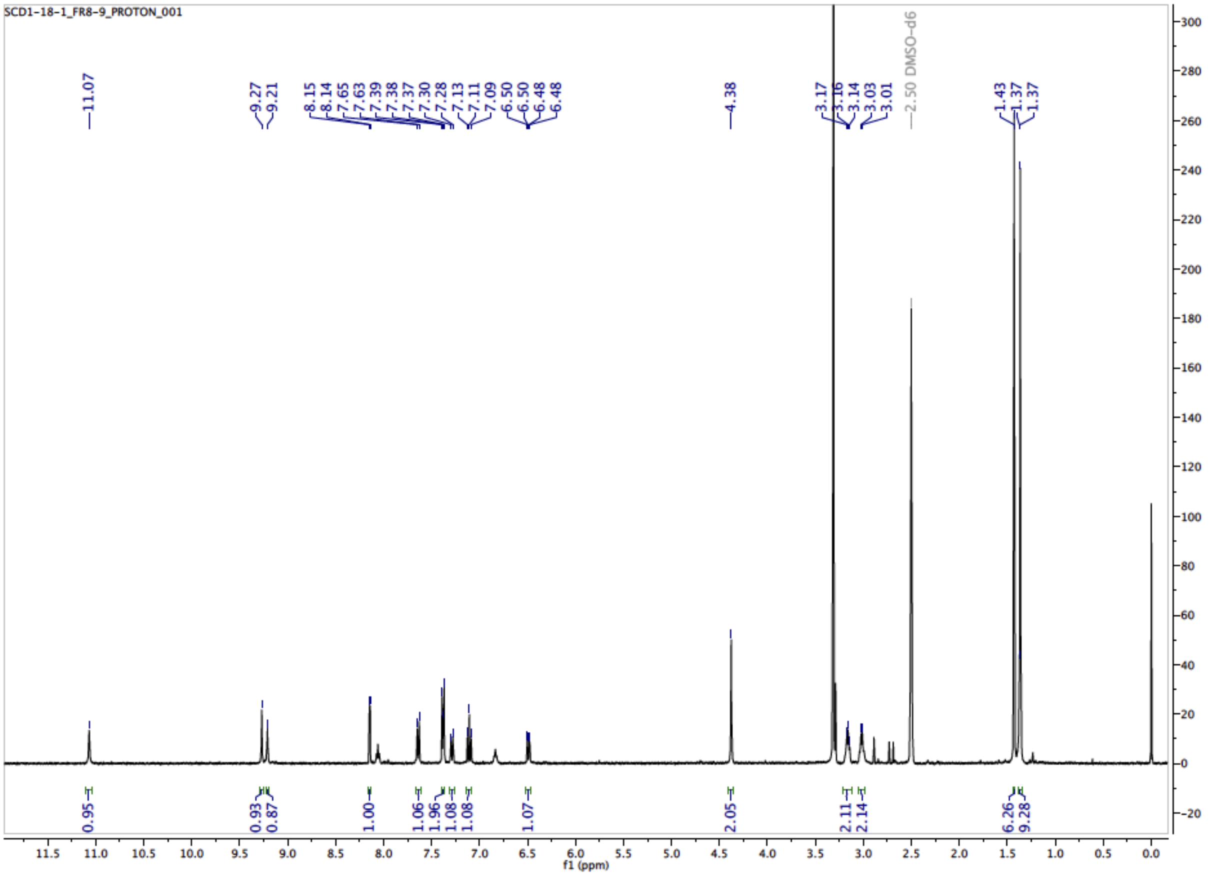


**LCMS**

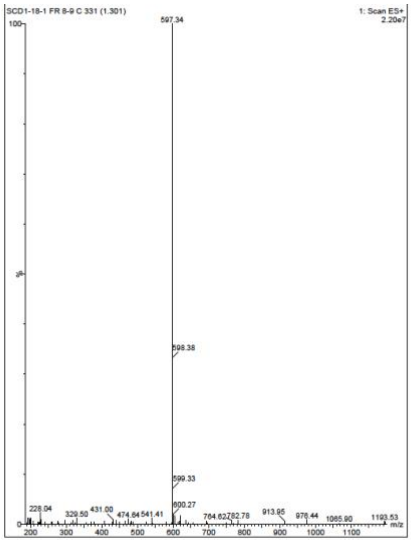

***N*-(2-aminoethyl)-2-(3-((4-((2,2-dimethyl-3-oxo-3,4-dihydro-2*H*-pyrido[3,2-*b*][1,4]oxazin-6-yl)amino)-5-fluoropyrimidin-2-yl)amino)phenoxy)acetamide (17)**

**$^{1}$H NMR**

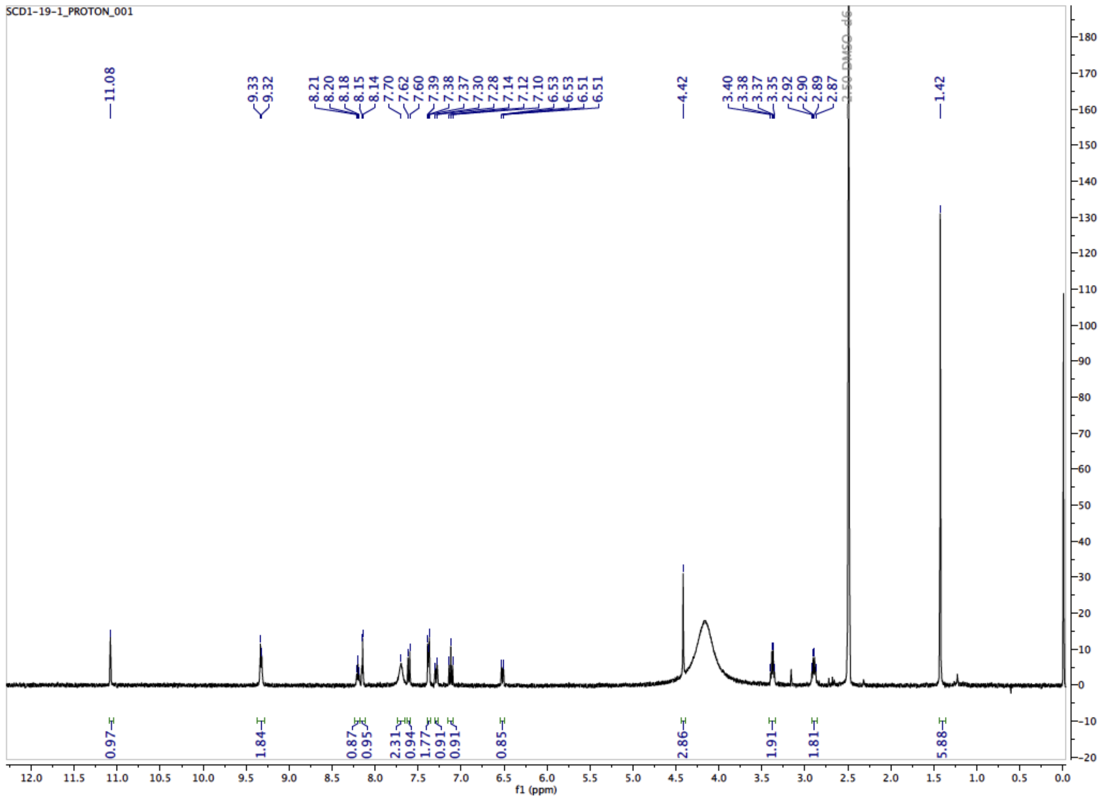


**LCMS**

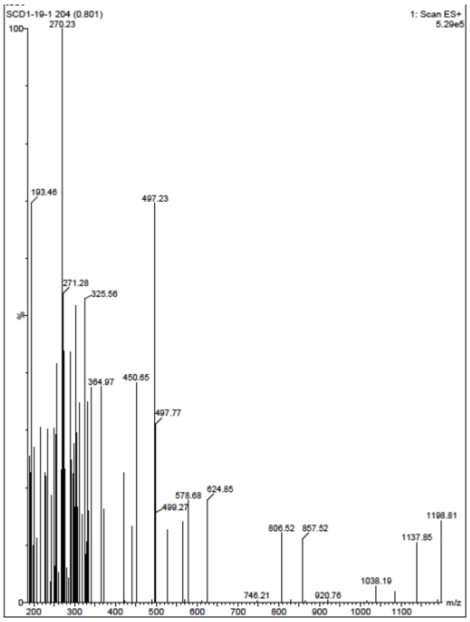

**3-(5,5-difluoro-7-(1*H*-pyrrol-2-yl)-5*H*-5$\lambda^4$,6$\lambda^4$-dipyrrolo[1,2-*c*:2',1'-*f*][1,3,2]diazaborinin-3-yl)-*N*-(2-(2-(3-((4-((2,2-dimethyl-3-oxo-3,4-dihydro-2*H*-pyrido[3,2-*b*][1,4]oxazin-6-yl)amino)-5-fluoropyrimidin-2-yl)amino)phenoxy)acetamido)ethyl)propenamide (6)**

#### $^1$H NMR

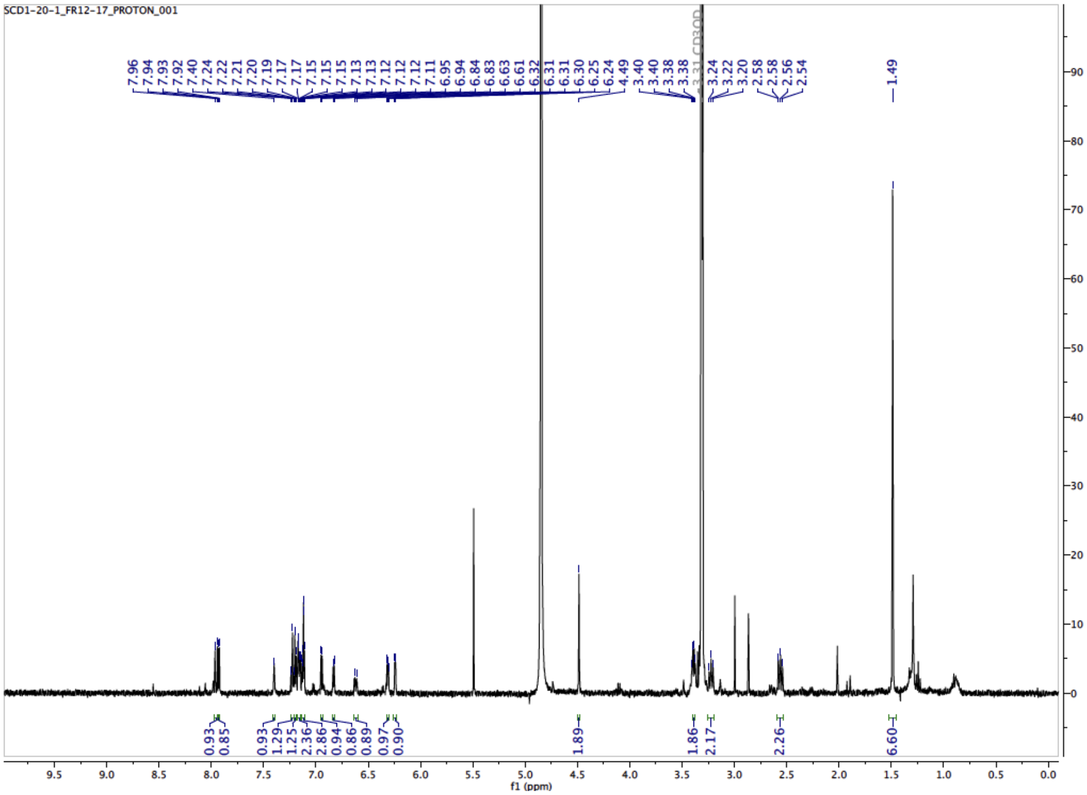


#### LCMS

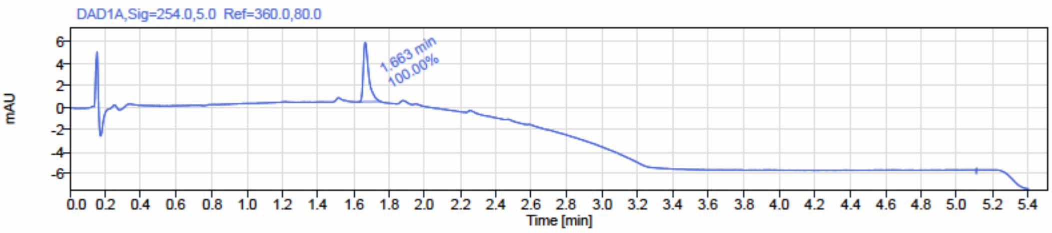


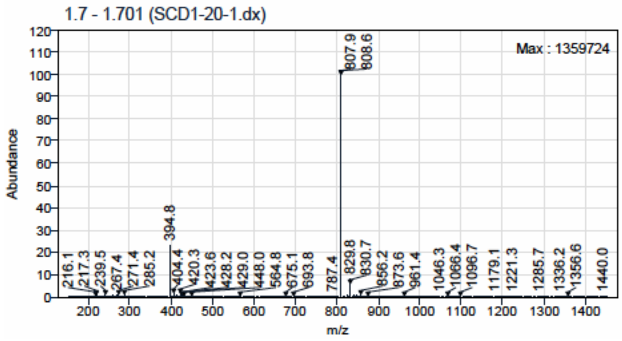

***N*-(2-(2-(2-(2-aminoethoxy)ethoxy)ethoxy)ethyl)-2-(3-((4-((2,2-dimethyl-3-oxo-3,4-dihydro-2*H*-pyrido[3,2-*b*][1,4]oxazin-6-yl)amino)-5-fluoropyrimidin-2-yl)amino)phenoxy)acetamide (18)**

**LCMS (Intermediate :** *tert*-butyl (1-(3-((4-((2,2-dimethyl-3-oxo-3,4-dihydro-2*H*-pyrido[3,2-*b*][1,4]oxazin-6-yl)amino)-5-fluoropyrimidin-2-yl)amino)phenoxy)-2-oxo-6,9,12-trioxa-3-azatetradecan-14-yl)carbamate)

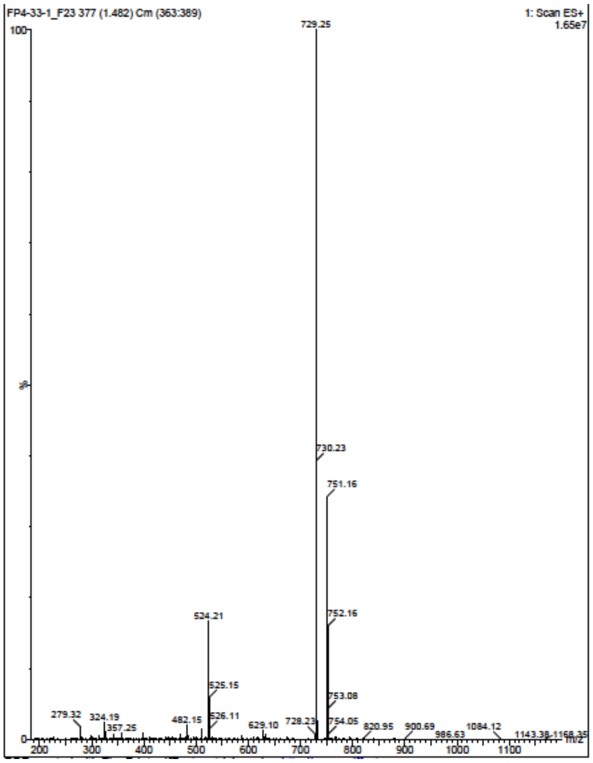


**LCMS**

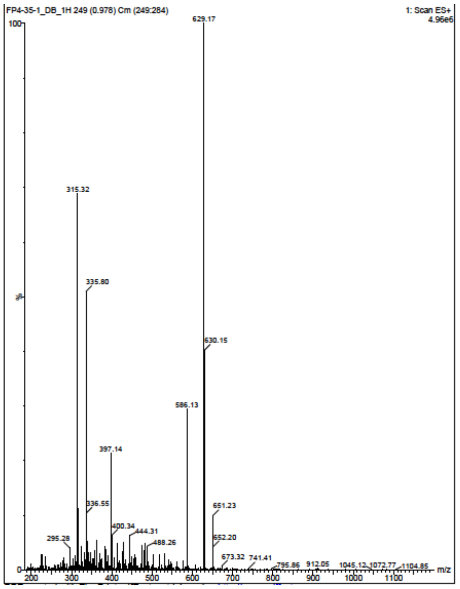

**3-(5,5-difluoro-7-(1*H*-pyrrol-2-yl)-5*H*-5$\lambda^4$,6$\lambda^4$-dipyrrolo[1,2-*c*:2',1'-*f*][1,3,2]diazaborinin-3-yl)-*N*-(1-(3-((4-((2,2-dimethyl-3-oxo-3,4-dihydro-2*H*-pyrido[3,2-*b*][1,4]oxazin-6-yl)amino)-5-fluoropyrimidin-2-yl)amino)phenoxy)-2-oxo-6,9,12-trioxa-3-azatetradecan-14-yl)propenamide (5)**

**$^1$H NMR**

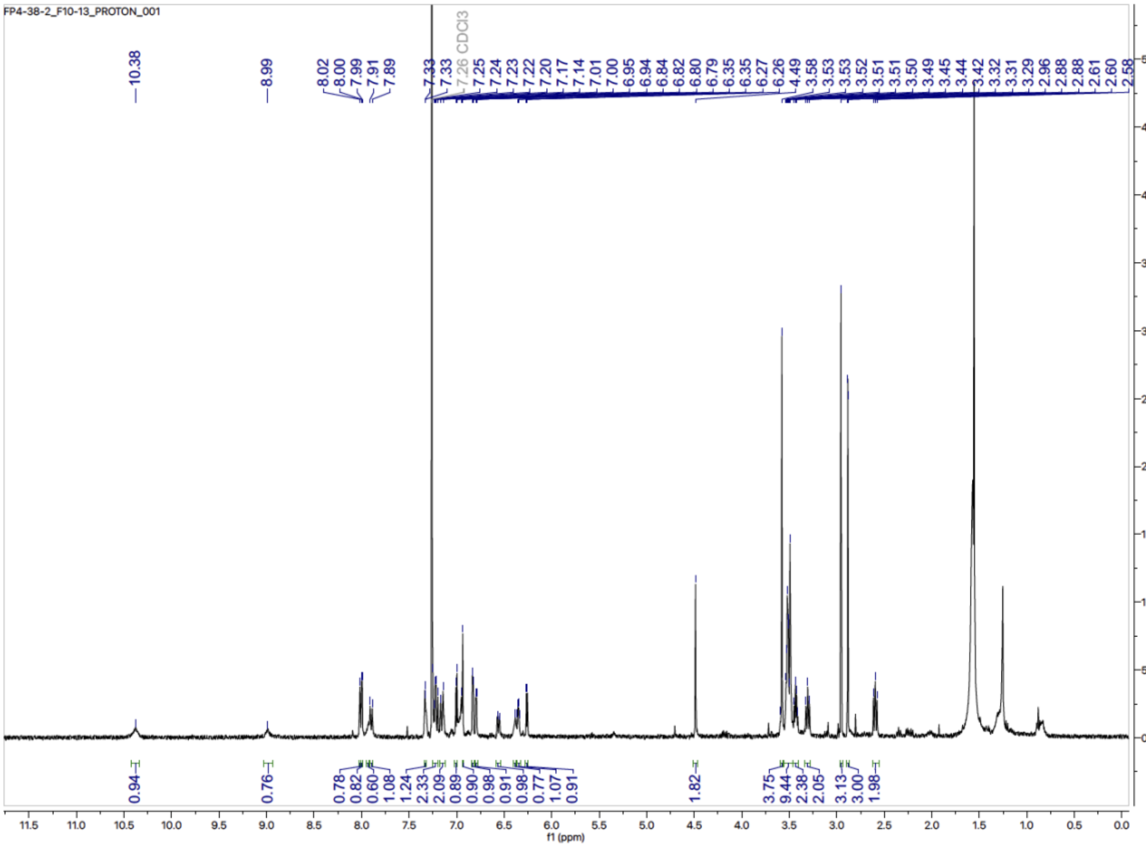


**LCMS**

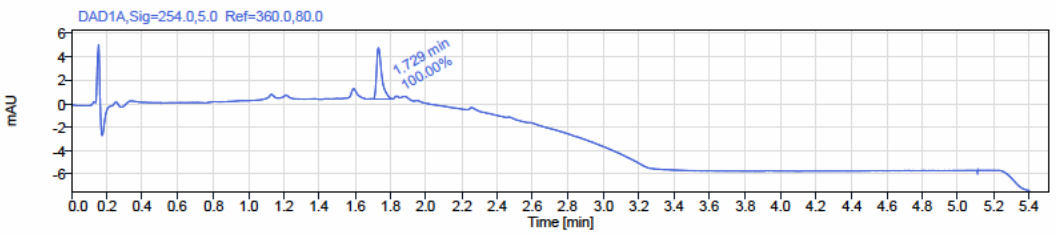


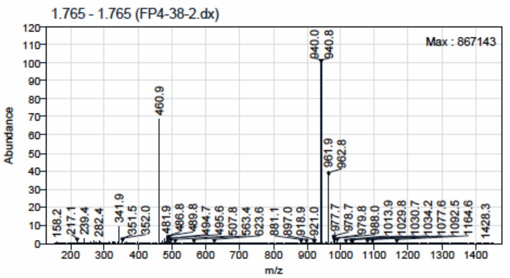

**(1*S*,4*R*)-*N*-(15-(5,5-difluoro-7-(1*H*-pyrrol-2-yl)-5*H*-5λ$^4$,6λ$^4$-dipyrrolo[1,2-*c*:2',1'-*f*][1,3,2]diazaborinin-3-yl)-13-oxo-3,6,9-trioxa-12-azapentadecyl)-4-hydroxy-2,2-dimethyl-4-(5-(3-methyl-5-((4-methylpyrimidin-2-yl)amino)phenyl)thiazol-2-yl)cyclohexane-1-carboxamide (7)**

**$^1$H NMR**

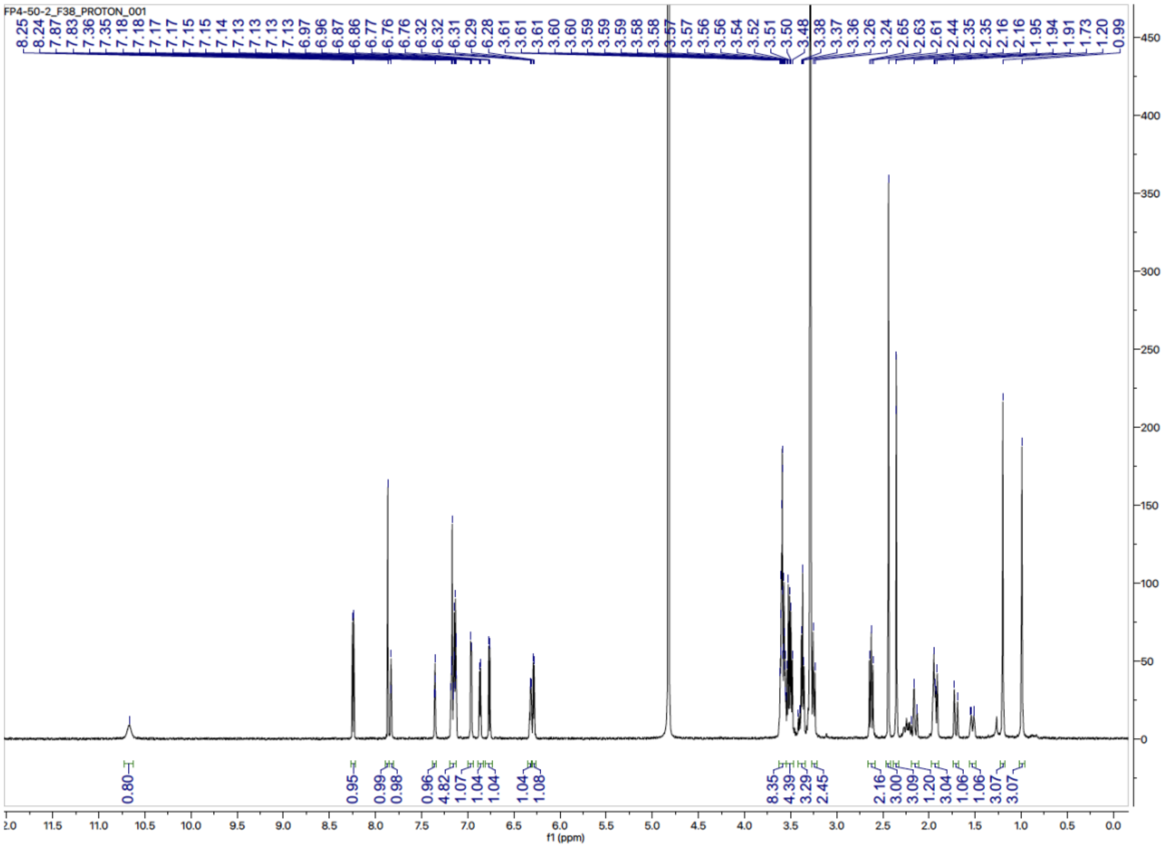


**LCMS**

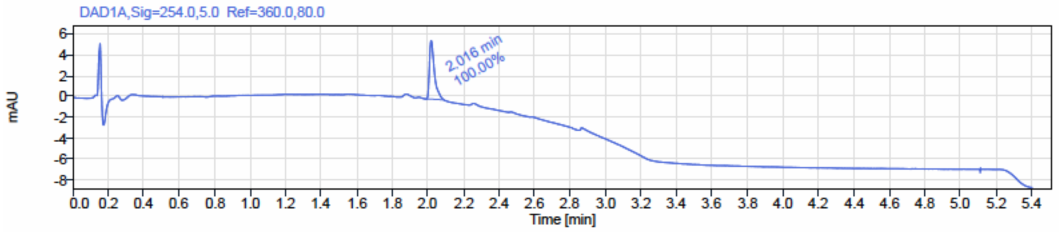

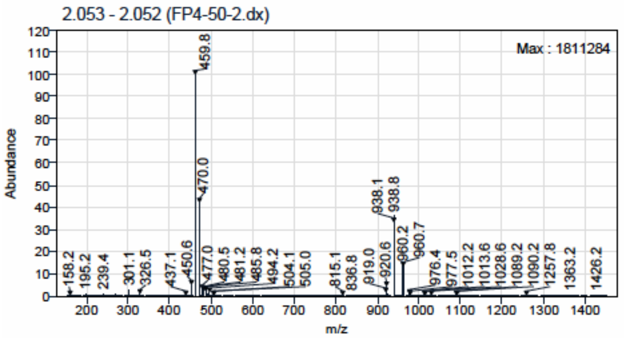


**(1*S*,4*R*)-*N*-(3-(3-(5,5-difluoro-7-(1*H*-pyrrol-2-yl)-5*H*-5λ⁴,6λ⁴-dipyrrolo[1,2-*c*:2',1'-*f*][1,3,2]diazaborinin-3-yl)propanamido)propyl)-4-hydroxy-2,2-dimethyl-4-(5-(3-methyl-5-((4-methylpyrimidin-2-yl)amino)phenyl)thiazol-2-yl)cyclohexane-1-carboxamide (8)**

#### $^{1}$H NMR

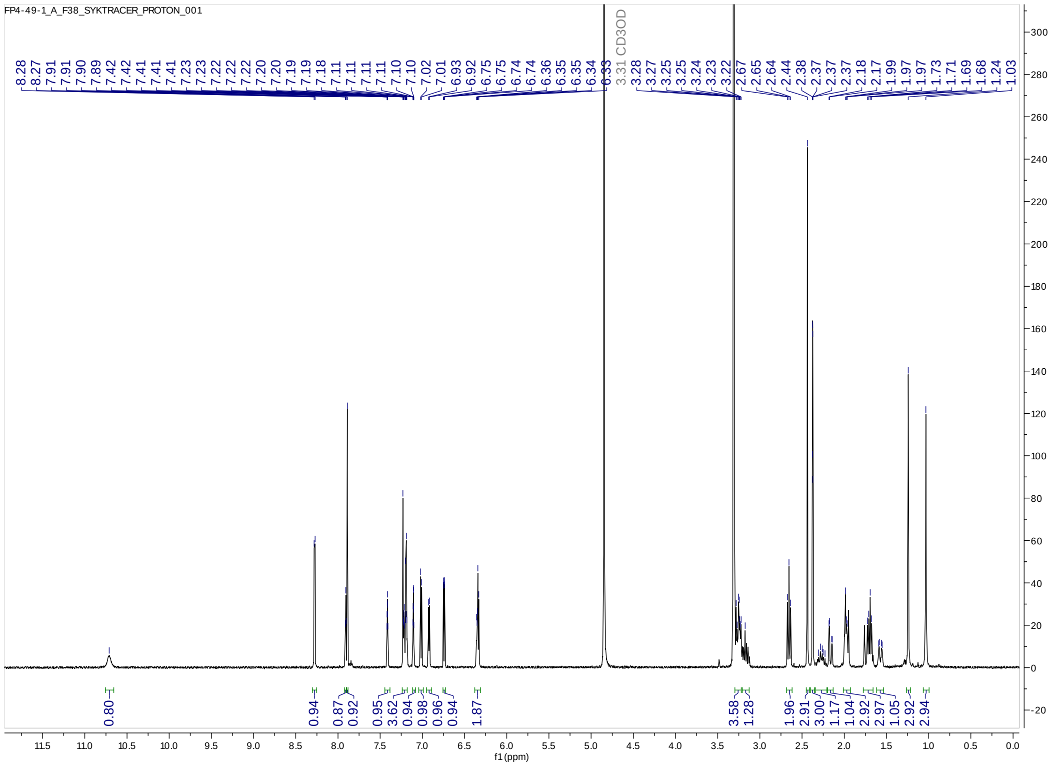


#### LCMS

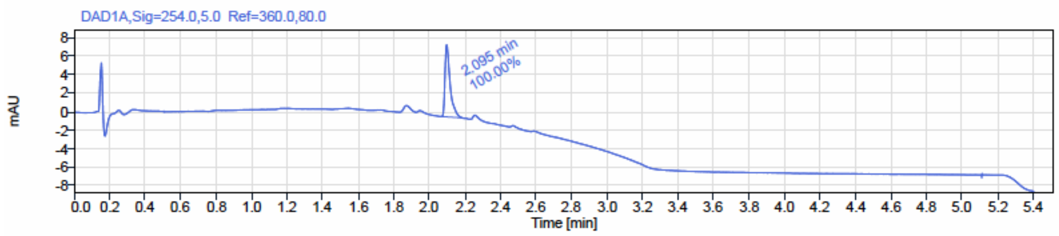

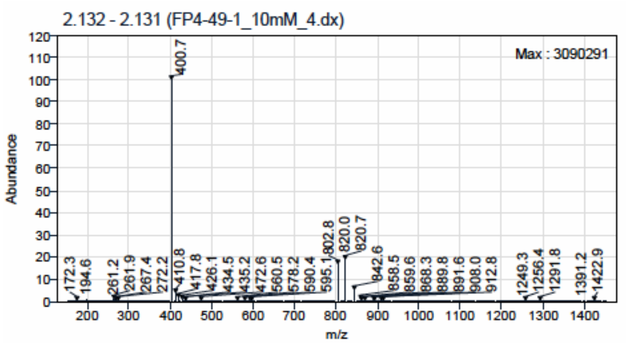
2.132 - 2.131 (FP4-49-1_10mM_4.dx)
Max : 3090291
Abundance
m/z
400.7
802.8
820.0
820.7
842.6
172.3
194.6
261.2
261.9
267.4
272.2
410.8
417.8
426.1
434.5
435.2
472.6
560.5
578.2
590.4
595.1
858.5
859.6
868.3
889.8
891.6
908.0
912.8
1249.3
1256.4
1291.8
1391.2
1422.9
